# *TET2*-mutant clonal hematopoiesis prevents T-cell exhaustion and suppresses cancer metastasis

**DOI:** 10.64898/2026.03.05.709727

**Authors:** Ai Nhan Thi Le, Manabu Fujisawa, Giacomo Campo, Yen T M Nguyen, Tran B. Nguyen, Sakurako Suma, Yasuhito Suehara, Fumihito Miura, Hiromitsu Araki, Kenichi Makishima, Tatsuhiro Sakamoto, Mizuho Nakayama, Masanobu Oshima, Miwako Kakiuchi, Shumpei Ishikawa, Chizu Tanikawa, Koichi Matsuda, Ryunosuke Saiki, Seishi Ogawa, Robert J. Vanner, Mamiko Sakata-Yanagimoto

## Abstract

Clonal hematopoiesis (CH) is widely regarded as a risk factor for age-associated disease, yet its influence on solid tumor progression remains poorly understood. Here, we uncover an unexpected tumor-suppressive role for *TET2*-mutant CH in solid tumors. Through an analysis of over 16,000 cancer patients, we demonstrate that *TET2* mutations are associated with a significant reduction in metastatic burden. Mechanistically, we show that *Tet2* deficiency prevents terminal exhaustion in CD8^+^ tumor-infiltrating lymphocytes (CD8^+^ TILs). This resistance is driven by DNA hypermethylation at regulatory elements of the transcription factor *Tox*, which prevents the induction of the terminal exhaustion program. Consequently, *Tet2*-deficient CD8^+^ TILs maintain stem-like, effector-competent states that mediate durable anti-tumor immunity and suppress metastatic outgrowth. Our findings establish *TET2*-mutant CH as a natural epigenetic constraint on T-cell exhaustion and uncover a mechanistically defined pathway with translational potential for enhancing anti-tumor immune responses.

## INTRODUCTION

Metastasis represents the principal cause of cancer-related mortality. Clinical metastatic disease is present in 4% to 65% of patients with solid malignancies at the time of cancer diagnosis^1^. Metastasis occurs through a complex and multi-step cascade wherein malignant cells spread beyond the primary tumor through the blood and lymph systems, colonize distant organs, and eventually manifest as metastatic lesions^2^. Immune cells recruited to metastatic sites undergo extensive reprogramming, and contribute to the formation of tumor-promoting or tumor-restraining immune microenvironments^3^. Indeed, metastatic tumor-infiltrating leukocytes exhibit dual effects on metastasis. Cytotoxic CD8^+^ tumor-infiltrating lymphocytes (TILs) and natural killer (NK) cells can restrain metastatic tumor cells under certain conditions^4–6^. However, their anti-tumor function is frequently compromised by exhaustion, characterized by upregulation of inhibitory receptors (PD-1, TIM3, LAG3) and impaired effector functions within the immunosuppressive tumor microenvironment^7–9^. Similarly, while macrophages, neutrophils, IL-17-producing γδ T cells, and regulatory T cells are commonly associated with promoting metastasis outgrowth^10–12^, specific subsets within these populations can exert anti-metastatic effects depending on their polarization state and local microenvironmental cues. For instance, M1-polarized macrophages exhibit anti-tumor activity through production of pro-inflammatory cytokines and direct cytotoxicity^13,14^, and presentation of tumor- antigens^15^, N1-polarized neutrophils can suppress tumor progression through reactive oxygen species production and immune activation^16,17^, and certain γδ T cell subsets demonstrate potent cytotoxic activity against tumor cells and support anti-tumor immunity^18–20^. These observations indicate that the phenotypic heterogeneity and functional state of tumor-infiltrating immune cells, shaped by local microenvironmental cues, critically influence metastatic colonization.

Clonal hematopoiesis (CH), the age-associated acquisition of somatic mutations in hematopoietic stem and progenitor cells, has emerged as a major source of systemic inflammation and immune dysregulation. Analysis of patient samples and animal models indicates that CH-derived immune cells underlie these inflammatory changes^21^. The prevalence of CH in solid cancer patients is 20-30%, which is significantly higher than in age-matched healthy individuals^22,23^. *DNMT3A*, *TET2*, and *ASXL1* are the most commonly mutated genes driving CH in untreated solid cancer patients and in healthy individuals^23^, whereas exposure to cancer therapy heightens the frequency of mutations in DNA damage response genes, such as *TP53*, *CHEK2,* and *PPM1D*^22^. Indeed, CH-derived mutant alleles are frequently detected in tumor-infiltrating immune cells of solid cancer patients with CH^23–26^. Tumor infiltration by CH-derived mutant immune cells shapes the local immune microenvironment, which has been associated with tumor inflammation and poor prognosis^27,28^. However, the clinical and biological implications of CH-derived immune cells on solid cancer remain incompletely understood. Mouse studies revealed conflicting results regarding whether CH-associated immune alterations accelerate or restrain tumor growth across different cancer types^29,30^. Pan-human cancer analyses suggest CH-outcome associations vary based on cancer type^22,23,31,32^, and CH-driver specific outcome analyses are lacking. Recently, *TET2*-mutant CH was shown to potentiate response to immune checkpoint blockade in colorectal cancer (CRC), non-small cell lung cancer, melanoma, and animal models^15,29,33^. Therefore, TET2 inactivation in tumor-infiltrating myeloid and lymphoid cells can enhance anti-tumor immunity. Intriguingly, the potential roles of CH-derived immune cells on cancer metastasis have not been clarified.

Here, we examined the association between CH and metastasis in a large clinical cohort and further investigated how *Tet2*-deficient CH-derived immune cells regulate CRC liver metastasis using conditional *Tet2* knockout mouse models.

## RESULTS

### *TET2*-mutant CH is associated with lower metastatic cancer burden

We analyzed CH in 16,744 non-hematological cancer patients from the MSK-IMPACT cohort with matching demographic and clinical data including treatment history, cancer stage, and metastatic burden^22,34^. Driver clonal hematopoiesis (CH) was defined as putative CH-driver mutations with a variant allele frequency (VAF) ≥ 0.02 (Methods). Reproducing the observations from the larger cohort of Bolton et al. 2020 ^22^, overall CH prevalence was 19.6%, most commonly with *DNMT3A* (10.2%), *TET2* (3.1%), and *PPM1D* (2.6%) driver mutations (Figure S1A). Prevalence increased with age, and patients with driver CH were significantly older (Figures S1B and S1C). Known associations between *ASXL1* and smoking history, and with *CHEK2*, *TP53*, *PPM1D* and prior exposure to cancer therapy, were recapitulated (Figures S1D and S1E). Metastatic cases were more likely to have received therapy (Figure S1F), and CH prevalence patterns varied by cancer type in accordance with prior studies (Figures S1G and S1H).

To investigate a potential association between CH and cancer metastasis, we compared CH prevalence in patients with and without metastatic cancer. In multivariate adjusted analysis, there was a trend towards higher prevalence of driver CH in patients without metastatic disease (OR 0.89, 95%CI [0.78–1.01], FDR-adjusted *P* = 0.109, Figure 1A). CH with a *TET2*-driver mutation was significantly more common in patients with local compared to metastatic cancers (OR 0.64, 95%CI [0.46–0.90], *P =* 0.029; Figure 1A), while *DNMT3A*-driven CH and other forms of CH with non-*DNMT3A*, non-*TET2* drivers were not associated with metastatic status (Figure 1A). Non-driver somatic *TET2* variants were not associated with metastatic status (Figure S1I). By cancer type, *TET2*-driver CH was associated with lower odds of having metastatic breast cancer (multivariate logistic regression, OR 0.36, *P =* 0.022), with similar trends in non–small cell lung cancer (NSCLC) (OR 0.52, *P* = 0.064) and in CRC (OR 0.53, *P =* 0.419) (Figure 1B, Table S1). *TET2*-CH was numerically but not statistically more common in patients with metastatic prostate cancer (multivariate logistic regression, OR 1.98, *P =* 0.419). When examining metastatic status at specific sites, patients with *TET2*-driver CH showed significantly reduced odds of having liver (OR 0.67, *P =* 0.016) or brain metastasis (OR 0.54, *P =* 0.016), and showed a trend towards lower odds of bone metastasis (OR 0.77, *P =* 0.114; Figure 1C).

**Figure 1.**
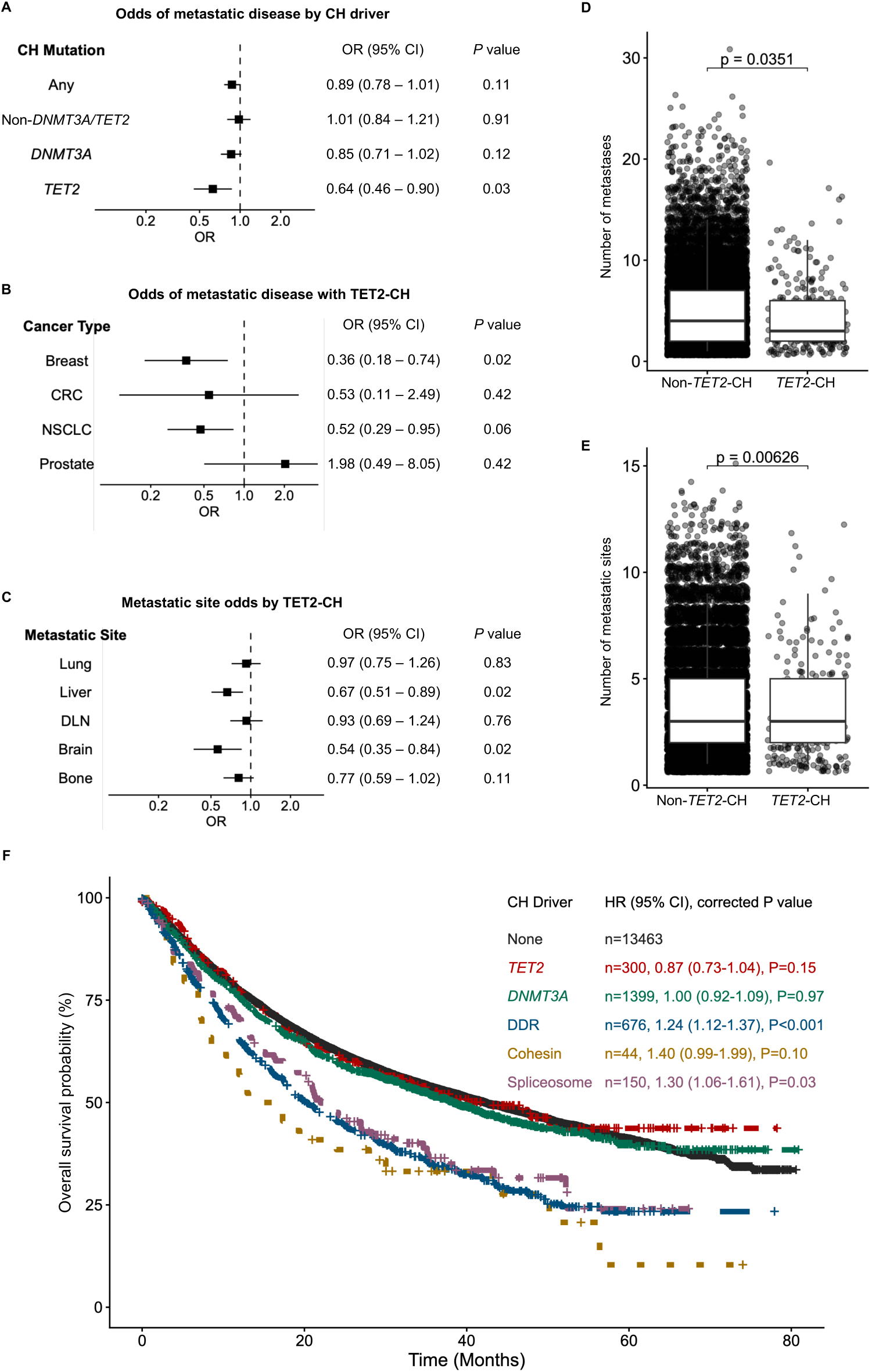
*TET2*-mutant CH is associated with reduced cancer metastasis. (A) Adjusted odds of patient having metastatic versus local cancer by CH-mutant status with any CH-driver mutation, or stratified by (bottom) only *DNMT3A* driver, only *TET2* driver, and non-*TET2* non-*DNMT3A* mutant CH, with the reference of no CH. ORs and 95% confidence intervals are reported from logistic GLMs, and *P* values are FDR-adjusted. (B) Adjusted odds of patient having metastatic versus local cancer given exposure to *TET2*-mutant CH for the four most prevalent cancer types. ORs and 95% confidence intervals are reported from logistic GLMs for each cancer type, and *P* values are FDR-adjusted (see also Table S1). (C) Adjusted odds of metastasis occurring at specific sites given exposure to *TET2*-mutant CH. ORs and 95% confidence intervals are reported from logistic GLMs for each metastatic site, and *P* values are FDR-adjusted. (D) The number of metastases in patients with metastatic cancer stratified by *TET2*-mutant CH presence. Differences in distributions are quantified both univariately by Wilcoxon rank sum tests, and multivariately adjusted with IRRs from negative binomial generalized linear models (Supplementary Figure1J). (E) The distribution of the number of metastatic sites in patients with metastatic cancer stratified by *TET2*-mutant CH presence. Differences in distributions are quantified both univariately by Wilcoxon rank sum tests, and multivariately adjusted with IRRs from negative binomial generalized linear models (Supplementary Figure1K). (F) Kaplan-Meier (KM) survival curves for overall survival of all patients, stratified by no CH mutation, and *DNMT3A*-, *TET2*-, DNA-damage-response (DDR)-, cohesin-, and spliceosome-mutant CH. Adjusted hazard ratios (HR) are reported from a Cox proportional hazards model, with the reference of no CH, and *P* values are FDR-adjusted. CH, clonal hematopoiesis; OR, Odds ratio; GLM, generalized linear model; FDR, false discovery rate; IRR, incidence risk ratio. Detailed statistical outputs, test statistics, and model parameters for all panels in Figure 1 are provided in Table S2.

When examining patients with metastatic disease, *TET2*-driver CH was linked to lower metastatic disease burden as quantified by number of metastases (median and interquartile range [IQR] metastasis number for *TET2*-CH, 3 [2-6], versus no CH and non-*TET2*-CH, 4 [2-7], *P =* 0.035; multivariate negative binomial generalized linear model IRR = 0.92, *P =* 0.052; Figures 1D and S1J) and involvement of fewer metastatic sites (median and IQR metastasis site number for *TET2*-CH, 3 [2-5], versus no CH and non-*TET2*-CH, 3 [2-5], *P =* 0.006; multivariate negative binomial generalized linear model IRR = 0.92, *P =* 0.035; Figure 1E and S1K).

Overall, patients with driver CH mutations experienced modestly worse survival in multivariate analysis (HR 1.06, *P =* 0.0498; Figure S1L). Strikingly, this was not driven by *DNMT3A-* and *TET2-* driver CH (Figure 1F). While patients with *TET2*-driver CH trended towards improved OS in multivariate analysis compared with no CH (HR 0.87, *P* = 0.152; Figure 1F), patients with DDR- (HR 1.24, P < 0.001), cohesin- (HR 1.40, P = 0.096), and spliceosome-driver CH (HR 1.30, P = 0.034) experienced worse survival (Figure 1F). Therefore, solid tumor patients with *TET2*-CH are more likely to have local versus metastatic cancer, those with metastatic cancer tend to have fewer metastases, and in contrast to other forms of CH, *TET2*-mutant CH is not associated with worse overall survival. Based on these results, we explored whether *TET2*-mutant CH could suppress cancer metastasis in animal models.

### *Tet2* deficiency in T cells inhibits CRC liver metastasis in mice

We investigated the effects of hematopoietic *Tet2* deficiency on CRC metastasis to the liver, since *TET2*-CH was prominently associated with reduced metastasis to this site (Figure 1C) and CRC is one of the most common causes of liver metastasis. To explore the role *of Tet2* loss of function in different types of immune cells on CRC metastasis, we crossed mice carrying *LoxP*-flanked *Tet2* alleles (*Tet2^fl/fl^*)^35^ with *Vav1^Cre^ ^36^*, *Cd4^Cre^ ^37^*, *Cd19^Cre^ ^38^*, and *LysM^Cre^ ^39^*to generate mice with targeted *Tet2* knockout (KO) in all hematopoietic cells (*Vav1^Cre^Tet2KO* mice), B lymphoid cells (*Cd19^Cre^Tet2KO* mice), T lymphoid cells (*Cd4^Cre^Tet2KO* mice), and myeloid cells (*LysM^Cre^Tet2KO* mice), respectively. Venus-labeled mouse intestinal tumor-derived organoid cells carrying *Apc^Δ716^*, *Kras^+/G12D^*, *Tgfbr2^-/-^* and *Trp^R270H/LOH^* mutations (Venus-*AKTP^M/LOH^* cells)^40^, which acquired the ability to metastasize to the liver, were injected into the spleen of various *Tet2* conditional KO mice and *Tet2^fl/fl^* mice to generate CRC liver metastases (Figure 2A). After 30 days, liver metastatic tumor burden (LMTB) was lower in *Vav1^Cre^Tet2KO* mice and *Cd4^Cre^Tet2KO* mice than in *Tet2^fl/fl^* mice, while LMTB of *Cd19^Cre^Tet2KO* and *LysM^Cre^Tet2KO* mice was comparable with that of *Tet2^fl/fl^*mice (Figures 2B and 2C; Table S3). This result suggests that *Tet2* deficiency in T cells, but not in B cells or myeloid cells, is sufficient to restrict CRC liver metastasis in mice.

**Figure 2.**
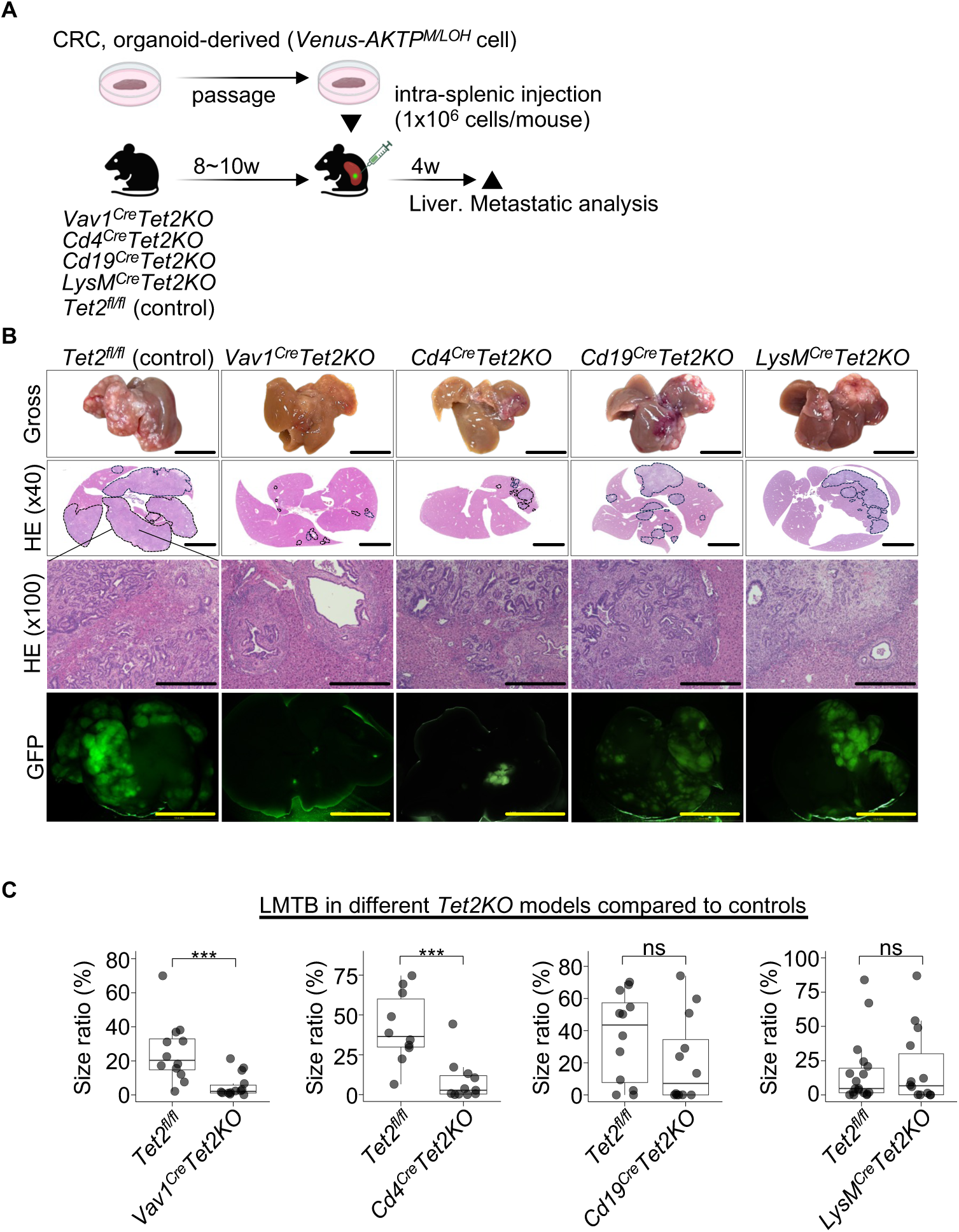
*Tet2* deficiency in T cells, but not in myeloid and B cells, inhibits the liver metastasis of CRC in mice. (A) Schematic of the CRC liver metastasis model using cell-type-specific *Tet2* knockout mice. CRC organoid-derived Venus-*AKTP^M/LOH^* cells (1×10^6^ cells/mouse) were intrasplenically injected into *Vav1^Cre^Tet2KO, Cd4^Cre^Tet2KO*, *Cd19^Cre^Tet2KO*, *LysM^Cre^Tet2KO*, and *Tet2^fl/fl^* control mice. Liver metastases were analyzed 4 weeks post-injection. (B) Representative macroscopic (top row), HE-stained (middle rows; 40× and 100× magnification), and GFP fluorescence (bottom row) images of liver metastases. Scale bars: 1 cm (macroscopic and GFP); 5 mm (HE, 40×); 500 µm (HE, 100×). (C) Quantification and comparisons of LMTB between different Tet2KO models and *Tet2^fl/fl^* controls. See also Table S3 for detailed quantification and statistical comparisons. CRC, colorectal cancer; HE, hematoxylin and eosin; GFP, green fluorescent protein; LMTB, liver metastatic tumor burden. Data in (C) are pooled from 3 independent experiments (total *n* = 11-14 mice per group). Statistical significance was determined by Mann-Whitney U test. \**p <* 0.05, \*\**p <* 0.01, \*\*\**p <* 0.001, \*\*\*\**p <* 0.0001, ns = not significant.

### *Tet2* deficiency reshapes CD8^+^ TILs through the *Tox-*exhaustion axis to enhance anti-tumor immunity

To determine which T cell subsets are regulated by *Tet2* in the anti-tumor response, we compared the transcriptional profiles of CD4^+^ TILs and CD8^+^ TILs isolated from CRC liver metastases of *Vav1^Cre^Tet2KO* mice*, Cd4^Cre^Tet2KO* mice, and *Tet2^fl/fl^* mice (Figure 3A). Principal component analysis (PCA) and heatmap clustering showed that biological replicates of CD8^+^ TILs from either *Vav1^Cre^Tet2KO* mice or *Cd4^Cre^Tet2KO* mice (*Vav1^Cre^/Cd4^Cre^Tet2KO* CD8^+^ TILs) clustered closely together but distinctly from their *Tet2^fl/fl^* counterparts, while *Tet2KO* CD4^+^ TILs showed no significant separation from *Tet2^fl/fl^*CD4^+^ TILs (Figures 3B and S2A, Table S4). Comparing *Vav1^Cre^/Cd4^Cre^Tet2KO* groups to *Tet2^fl/fl^* group identified substantially more differentially expressed genes (DEGs) in CD8^+^ TILs than in CD4^+^ TILs (Figure 3C). In the *Vav1^Cre^Tet2KO* model, CD8^+^ TILs exhibited 226 DEGs compared to 71 DEGs in CD4^+^ TILs. This disparity was even more pronounced in the *Cd4^Cre^Tet2KO* model, where CD8^+^ TILs demonstrated 209 DEGs versus only 8 DEGs in CD4^+^ TILs (FDR< 0.01, |Log2 FC| > 2) (Figure 3C, Table S5-S8). These findings suggest that *Tet2* deficiency affects the transcriptome of CD8^+^ TILs more than CD4^+^ TILs.

**Figure 3.**
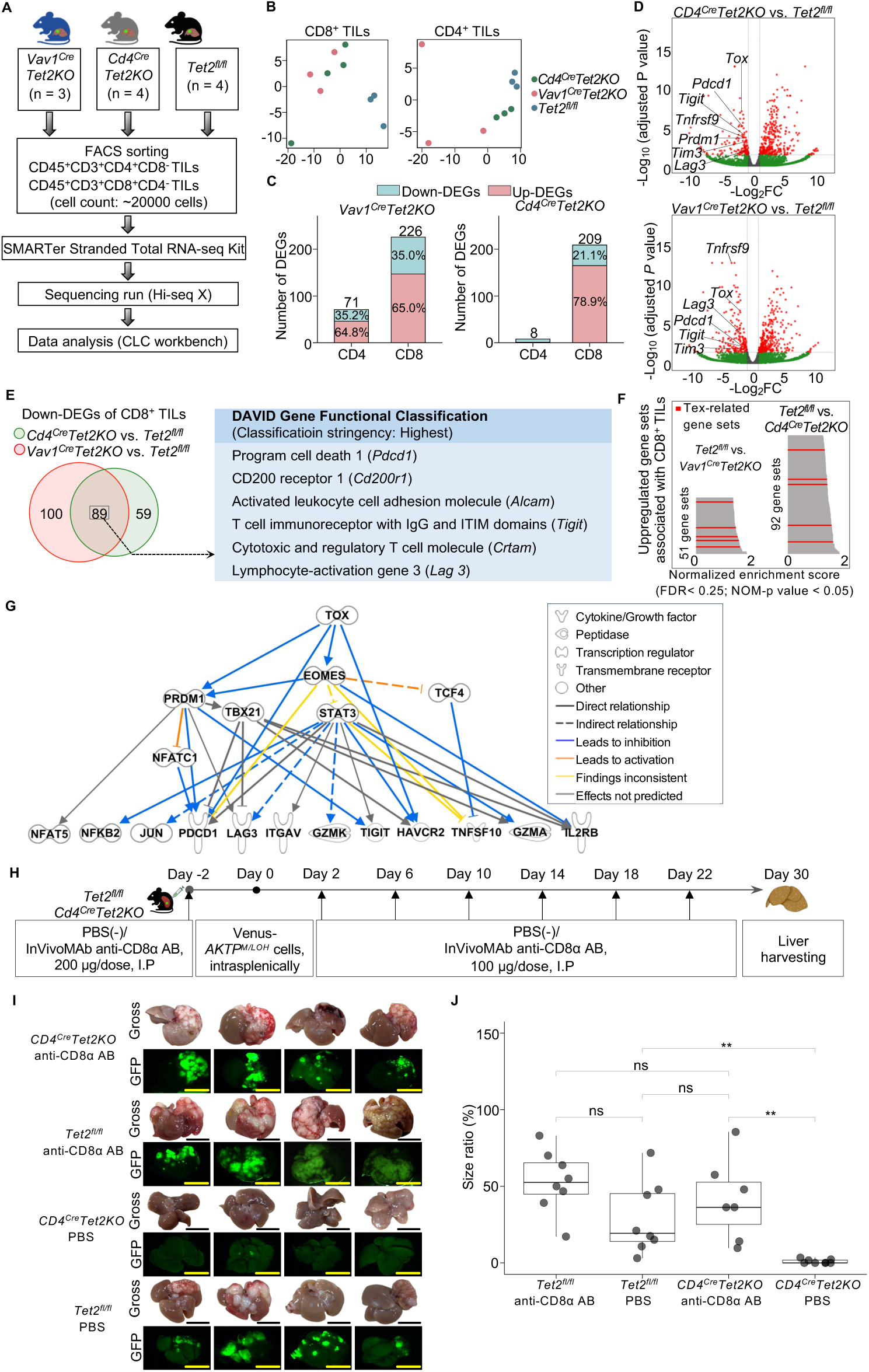
*Tet2* deficiency extensively reshapes CD8^+^ TILs transcriptional profiles through the *Tox-*exhaustion axis to enhance anti-tumor immunity in CRC liver metastasis. (A) Schematic of the experimental workflow for transcriptomic profiling. CD45⁺ tumor-infiltrating leukocytes were isolated via MACS and sorted by FACS into CD4⁺ and CD8⁺ T cells, respectively. Total RNA was extracted from sorted populations and subjected to library preparation using SMARTer Stranded Total RNA-seq Kit, followed by high-throughput sequencing (Hi-seq X) and data analysis (CLC workbench). (B) PCA plots of CD4 and CD8^+^ TILs supervised by the top 200 DEGs. Gene ranking was based on combined FDR calculated by multiplying the two individual FDR from *Vav1^Cre^Tet2KO* vs. *Tet2^fl/fl^*FDR and *Cd4^Cre^Tet2KO* vs. *Tet2^fl/fl^* comparisons (selection criteria: FDR < 0.05, |Log2 FC| > 1 in both comparisons) (see also Table S4). (C) Stacked bar plots show the number of upregulated and downregulated genes in CD4 and CD8^+^ TILs from the comparison between *Vav1^Cre^/Cd4^Cre^Tet2KO* groups to *Tet2^fl/fl^* group (FDR < 0.01, |Log2 FC| > 2). (D) Volcano plots show DEGs of *Vav1^Cre^/Cd4^Cre^Tet2KO* CD8^+^ TILs in comparison with *Tet2^fl/fl^* CD8^+^ TILs (FDR < 0.05, |Log2 FC| > 1). (E) Venn diagrams display 89 overlapping down-DEGs in the comparison between *Vav1^Cre^Tet2KO* and *Cd4^Cre^Tet2KO* with *Tet2^fl/fl^* CD8^+^ TILs (FDR < 0.05, Log2 FC < -1). Also shown is the DAVID functional classification of these overlapping genes, highlighting a cluster associated with immunologic function (Enrichment Score: 2.59; stringent parameters). (F) GSEA using ImmuneSigDB subset of C7 showing the enrichment of T-cell exhaustion-related signatures in *Tet2^fl/fl^* CD8^+^ TILs relative to *Vav1^Cre^/Cd4^Cre^Tet2KO* CD8^+^ TILs (FDR < 0.25, nominal *p <* 0.05). (G) Interaction network of *Tox* and T-cell exhaustion-related genes identified by IPA. The analysis was based on the 89 overlapping down-DEGs within CD8^+^ TILs from *Vav1^Cre^Tet2KO* and *Cd4*^Cre^*Tet2KO* groups compared to the *Tet2^fl/fl^*groups (FDR < 0.05, Log2 FC < -1). (H) Schematic of the CD8 T-cell depletion experiment. Venus-*AKTP^M/LOH^* cells were intrasplenically transplanted into *Cd4^Cre^Tet2KO* or *Tet2^fl/f^* mice. Anti-CD8α antibody or PBS was initiated 2 days before transplantation, and continued every 4 days throughout tumor progression. (I) Representative macroscopic and GFP fluorescence images of liver metastases 30 days post-injection. Scale bars, 1 cm. (J) Quantification of LMTB based on the ratio of tumor area to total liver area (%) See also Table S15 for detailed quantification and statistical comparisons. TILs, tumor-infiltrating lymphocytes; CRC, colorectal cancer; MACS, magnetic-activated cell sorting; FACS, fluorescence-activated cell sorting; PCA, principal component analysis; DEG, differentially expressed gene; GSEA, Gene Set Enrichment Analysis; IPA, Ingenuity Pathway Analysis; GFP, green fluorescent protein; LMTB, liver metastatic tumor burden; PBS, phosphate-buffered saline. Data in (J) are compiled of 2 independent experiments (total *n =* 7-8 mice per group). Statistical significance in (J) was determined by pairwise Mann-Whitney U tests with Bonferroni correction for multiple comparisons. \**P <* 0.05, \*\**P <* 0.01, \*\*\**P <* 0.001, \*\*\*\**P <* 0.0001, ns = not significant.

Notably, multiple exhaustion-associated genes including *Pdcd1*, *Lag3*, *Tim3*, *Tigit*, *Prdm1*, *Tnfrsf9*, and *Tox* were the most significantly downregulated genes in *Vav1^Cre^/Cd4^Cre^Tet2KO* CD8^+^ TILs relative to *Tet2^fl/fl^* CD8^+^ TILs, suggesting attenuated exhaustion programs in the absence of *Tet2*. (Figure 3D; Table S5 and S6). In CD8^+^ TILs, DEG analysis showed substantial concordance between *Vav1^Cre^Tet2KO* and *Cd4^Cre^Tet2KO* models with 198 commonly upregulated and 89 commonly downregulated genes, including downregulation of exhaustion markers *Tox* and *Pdcd1* (FDR< 0.05, |Log2 FC| > 1) (Figure S2B). Using DAVID gene functional classification^41^, we clustered 89 overlapping downregulated genes shared by *Vav1^Cre^Tet2KO* CD8^+^ TILs and *Cd4^Cre^Tet2KO* CD8^+^ TILs compared to *Tet2^fl/fl^* CD8^+^ TILs (Table S9A). Under stringent parameters (Table S9B), these consistently downregulated genes were classified into a single cluster comprising six genes associated with immunologic function, with an enrichment score of 2.59 (Figure 3E). Notably, this downregulated cluster encompassed key genes linked to T-cell exhaustion, including *Pdcd1*, *Lag3*, and *Tigit* (Figure 3E; Table S9C).

Gene Set Enrichment Analysis (GSEA)^42^ was performed using the ImmuneSigDB subset C7 from the Molecular Signatures Database (MSigDB)^43^ to compare the genome expression profiles of *Tet2KO* CD8^+^ TILs and *Tet2^fl/fl^* CD8^+^ TILs. The analysis identified 92 gene sets significantly enriched in *Tet2^fl/fl^* CD8^+^ TILs compared to *Cd4^Cre^Tet2KO* CD8^+^ TILs, five of which were specifically associated with T-cell exhaustion (Figures 3F and S3A; Table S10). In the same approach, 51 gene sets related to CD8 T-cell function were enriched in *Tet2^fl/fl^* CD8^+^ TILs compared to *Vav1^Cre^Tet2KO* CD8^+^ TILs, including five exhaustion-associated signatures (Figures 3F and S3A, Table S11). Of the 34 gene sets consistently enriched in *Tet2^fl/fl^*CD8⁺ TILs relative to both *Vav1^Cre^Tet2KO* and *Cd4^Cre^Tet2KO* counterparts, four were linked to T-cell exhaustion (Figure S3B).

Ingenuity Pathway Analysis (IPA) of 89 consistently downregulated genes in *Vav1^Cre^Tet2KO* CD8^+^ TILs and *Cd4^Cre^Tet2KO* CD8^+^ TILs (Table S12) identified the transcription factor *Tox* as one of the potential upstream regulators targeting genes associated with T-cell exhaustion (overlap *p*-value = 0.00164) (Figure 3G; Table S12). Consistent with this finding, *Tox* exhibited the greatest differential expression with the lowest *P* value among all tested genes, with levels markedly higher in *Tet2^fl/fl^* CD8^+^ TILs compared to *Vav1^Cre^/Cd4^Cre^Tet2KO* CD8^+^ TILs (*Tet2^fl/fl^*vs. *Vav1^Cre^Tet2KO*, *P =* 0.0002; *Tet2^fl/fl^*vs. *Cd4^Cre^Tet2KO*, *P =* 0.0001; Figure S3C; Table S13).

Furthermore, flow cytometry analysis of CRC metastatic tumors in the liver revealed a significantly higher percentage of CD8^+^ TILs co-expressing exhaustion markers Tim3 and PD-1 of the *Tet2^fl/fl^* group compared with that of the *Vav1^Cre^/Cd4^Cre^Tet2KO* group (Figure S4A; Table S14A). Indirect immunofluorescence staining further demonstrated a higher number of CD8α cells co-expressing Tim3, PD-1 and Tox in tumors from the *Tet2^fl/fl^* group than in the *Vav1^Cre^/Cd4^Cre^Tet2KO* groups (Figure S4B; Table S14B). Collectively, these findings identify *Tet2* as a critical enforcer of the terminal exhaustion program, which is intrinsically linked to high *Tox* expression.

To further validate the role of *Tet2* in regulating the antitumor functions of CD8^+^ TILs, we performed CD8 T-cell depletion experiments in *Cd4^Cre^Tet2KO* and *Tet2^fl/fl^* mice (Figure 3H). Circulating and liver-infiltrating CD8 T cells were efficiently and specifically depleted in anti-CD8α antibody-treated groups compared with PBS controls (Figures S5A and S5B). CD8 T-cell depletion in *Cd4^Cre^Tet2KO* mice drastically increased LMTB compared to their PBS-treated controls (*P =* 0.007), while CD8 T-cell depletion in *Tet2^fl/fl^* mice had no significant effect (*P =* 0.396) (Figures 3I, 3J and S5C; Table S15). As a result, LMTB of anti-CD8α antibody-treated *Cd4^Cre^Tet2KO* mice was not significantly different from that of anti-CD8α antibody-treated *Tet2^fl/fl^* mice (*P =* 1.0) and was comparable to that of PBS-treated *Tet2^fl/fl^* mice (*P =* 1.0), indicating that CD8 T-cell depletion attenuated the tumor-restrictive effect observed in *Tet2*-deficient mice (Figures 3I, 3J and S5C; Table S15). In contrast, PBS-treated *Cd4^Cre^Tet2KO* mice revealed a significantly lower LMTB compared to PBS-treated *Tet2^fl/fl^* mice (*P =* 0.007) (Figures 3I, 3J and S5C; Table S15), consistent with our earlier results. Collectively, these data demonstrate that the enhanced tumor control observed in *Cd4CreTet2KO* mice requires CD8^+^ TILs.

### *Tet2* deficiency preserves effector function and restricts terminal differentiation of CD8^+^ TILs

To comprehensively characterize the transcriptional landscape of *Tet2*-deficient CD8^+^ TILs, we performed single cell RNA sequencing (scRNA-seq) with V(D)J profiling. We profiled 16,790 and 17,367 CD45^+^ tumor-infiltrating leukocytes isolated from liver metastatic tumors of *Tet2^fl/fl^* (*n =* 2) and *Vav1^Cre^Tet2KO* (*n =* 2), respectively (Figure S6A). CD8^+^ TIL clusters were identified *in silico* from CD45^+^ tumor-infiltrating leukocytes (Figures S6B-S6J). *In silico*-defined CD8^+^ TILs were subjected to unsupervised sub-clustering, resulting in the identification of 9 distinct clusters, consisting of stem-like CD8 T cells (T_SLC_, *Ly6c2*^low^CD8 T_SLC_ and *Ly6c2*^high^CD8 T_SLC_), precursor/progenitor T cells (T_PROG_), cytotoxic/effector-like exhausted T cells (T_EFF-1_ and T_EFF-2_), intermediate exhausted T cells (T_INT-1_ and T_INT-2_), terminally exhausted T cells (T_TEX_), and proliferative exhausted T cells (P-T_EX_) (Figures 4A and 4B). T_PROG_ was characterized by the high expression of stem-like T-cell markers (*Sell*, *Ccr7*, *Tcf7*, *Il7r* and *Id3*) along with the dual expression of effector-like and exhausted T-cell markers at moderate or low levels (Figure 4C, Table S16). T_EFF_ (T4 and T5) cells displayed moderate to high expression of cytolytic/effector-like and exhaustion markers. T_TEX_ (T8) cells were defined by the high expression of multiple exhaustion markers and reduced expression in effector-like markers (Figure 4C, Table S16). Among the spectrum of CD8^+^ TILs, T_PROG_ (T3) and T_EFF_ (T4 and T5) were predominant in *Vav1^Cre^Tet2KO* mice, while T_INT_ (T6 and T7), T_TEX_ (T8) and P-T_EX_ (T9) were dominant in control *Tet2^fl/fl^* mice (Figure 4B). Among stem-like CD8^+^ TILs, *Ly6c2*^high^T_SLC_ (T2) was markedly enriched in *Tet2^fl/fl^* compared to *Vav1^Cre^Tet2KO* mice (Figure 4B). DEG analysis revealed significant upregulation of exhaustion-associated genes, including *Tox, Tim3, Tnfrsf9, Eomes,* and *Gzmk*, in *Tet2^fl/fl^*CD8^+^ TILs comparing to *Tet2KO* CD8^+^ TILs, consistent with bulk RNA-seq data (Figure 4D; Table S17). In contrast, cytolytic and effector-related genes, such as *Klrc1, Klrk1, Ifng, Tnf, Cd160, Ly6a, Cxcr3, Id2, Nr4a,* and *Ccl4*, were significantly upregulated in *Tet2KO* CD8^+^ TILs (Figure 4D; Table S17). Gene-set activity measured by AUCell scoring^44^ further demonstrated significant enrichment of effector-versus exhausted-T-cell signatures (*p =* 0.013), as well as memory- versus exhausted-T-cell signatures (*p* < 0.0001), in *Tet2KO* CD8^+^ TILs compared with *Tet2^fl/fl^*CD8^+^ TILs (Figure 4E).

**Figure 4.**
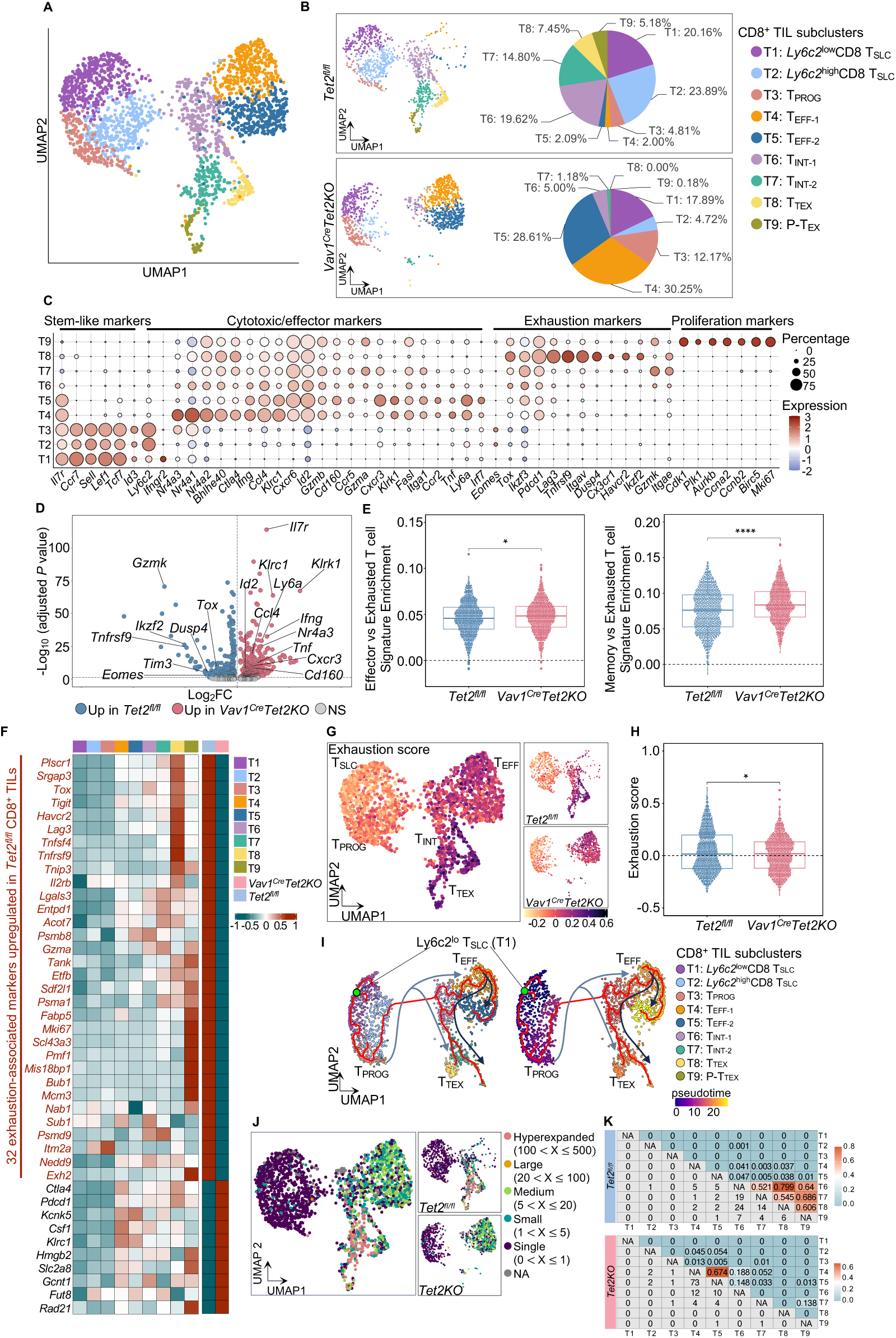
*Tet2* deficiency preserves effector function and restricts terminal differentiation of CD8^+^ TILs in CRC liver metastasis. (A) Nine subclusters were labeled with different colors on the UMAP plot of the total 2,202 CD8^+^ TILs from both *Tet2^fl/fl^*group (*n =* 2) and *Vav1^Cre^Tet2KO* group (*n =* 2). (B) UMAP plot of CD8^+^ TIL subclusters of each mouse group (1,101 CD8^+^ TILs for each group). The pie plots show percentages of each CD8^+^ TIL subcluster in each group. (C) Bubble plot showing the expression of selected marker genes among the top 50 most significant DEGs across defined clusters. The list of the top 50 most significant DEGs for each cluster are provided in Table S16. (D) Volcano plot displaying DEGs between *Tet2^fl/fl^* and *Vav1^Cre^Tet2KO* CD8^+^ TILs, highlighting T-cell exhaustion markers and cytolytic/effector-associated genes (adjusted *P* value < 0.05). List of DEGs is provided in Table S17. (E) Box plots comparing the enrichment scores of effector versus exhausted T-cell signatures (*p =* 0.013) and memory versus exhausted T-cell signatures (*p <* 0.0001) between *Tet2^fl/fl^* and *Vav1^Cre^Tet2KO* CD8^+^ TILs. (F) Heatmap displaying the average expression of 42 exhaustion-associated marker genes across subclusters and genotypes. (G) UMAP plot with cells labeled by exhaustion score characterize the spectrum of exhausted T cells with positive scoring values increasing toward T_TEX._ (H) Box plot comparing exhaustion scores between *Tet2^fl/fl^* and *Tet2KO* CD8^+^ TILs (*p =* 0.046). (I) Pseudotime trajectory of CD8^+^ TIL subclusters inferred by Monocle3 (left), with cells ordered along the trajectory using *Ly6c2^low^*T_SLC_ as the root (right). Arrows indicate inferred differentiation paths toward T_EFF_ and T_TEX_ states. (J) UMAP plot with cells colored by TCR clonal size categories, showing the distribution of clones across states in each group. (K) TCR repertoire overlap analysis using the Morisita overlap index between progenitor-like, effector, and exhausted states across groups. TILs, tumor-infiltrating lymphocytes; CRC, colorectal cancer; UMAP, Uniform Manifold Approximation and Projection; DEG, differentially expressed gene; T_TEX_, terminally exhausted T cell; T_EFF_, cytotoxic/effector-like exhausted T cells; T_SLC_, stem-like CD8 T cell; TCR, T-cell receptor. Statistical significance in (E) and (H) was determined by Mann-Whitney U test. \**p <* 0.05, \*\**p <* 0.01, \*\*\**p <* 0.001, \*\*\*\**p <* 0.0001, ns = not significant.

To characterize the exhaustion phenotype, we analyzed the expression of a signature comprising 42 genes consistently associated with exhausted T cells^45^ (Table S18) via heatmap analysis (Figure 4F). T_SLC_ (T1 and T2) clusters exhibited lower average expression of most exhaustion-associated markers, consistent with their progenitor state. In contrast, T_TEX_ (T8) and P-T_EX_ (T9) clusters displayed high expression across multiple exhaustion markers (Figure 4F). Notably, 32 out of the 42 exhaustion-associated genes were highly expressed in *Tet2^fl/fl^*CD8^+^ TILs but downregulated in *Vav1^Cre^Tet2KO* CD8^+^ TILs (Figure 4F). To quantify this, we calculated the exhaustion score using this 42 gene signature. The T-cell exhaustion score delineated the spectrum of exhausted T cells with positive scoring values increasing toward the T_TEX_ cluster (Figure 4G). *Tet2^fl/fl^* CD8^+^ TILs exhibited a significantly higher exhaustion score than *Tet2KO* CD8^+^ TILs (Figure 4H).

Using *Monocle 3*^46^, we reconstructed the pseudotime trajectory of CD8^+^ TILs to investigate differentiation dynamics within the metastatic tumor microenvironment (Figure 4I). *Ly6c2*^low^T_SLC_ cells were located at the root of the trajectory, consistent with their progenitor-like identity. Along this trajectory, T_PROG_ cells differentiated toward T_EFF-1_ or diverged toward T_TEX_ after transitioning through T_INT_ states. A subset of T_EFF-1_ cells retained cytolytic capacity as inferred by effector signatures and progressed into T_EFF-2_, while others gradually lost effector function and transitioned to T_TEX_ (Figure 4I). We next examined the T-cell receptor (TCR) clonotype landscape using strict clonotype calling based on the combination of V(D)J genes and the nucleotide sequence of the CDR3 region^47^ (Figure 4J). Clonal expansion states were classified by TCR clone size: single (1 cell), small (2–5 cells), medium (6–20 cells), large (21–100 cells), and hyperexpanded (>100 cells). Hyperexpanded clones were predominantly enriched within T_INT_ and T_TEX_ clusters of *Tet2^fl/fl^*CD8^+^ TILs, reflecting a biased differentiation toward terminal exhaustion (Figures 4J, S7A and S7B).

Clonal structure analysis revealed significantly greater repertoire diversity in Tet2KO mice, whereas *Tet2^fl/fl^* mice exhibited marked oligoclonal dominance, particularly within exhausted T cell states (Figures S7C and S7D). Clone size distribution across differentiation states demonstrated that hyperexpanded clones in *Tet2^fl/fl^* were predominantly concentrated in exhaustion states (T_INT_ [T6-T7], T_TEX_ [T8], and P-T_EX_ [T9]), while medium and large clones were underrepresented in effector states (T_EFF-1_ and T_EFF-2_ [T4-T5]). Conversely, *Tet2KO* CD8^+^ TILs displayed more balanced clone size distribution (Figures 4J and S7E-S7F).

The TCR clonotype overlap analysis highlighted distinct differences in differentiation dynamics between *Tet2^fl/fl^* and *Tet2KO* CD8^+^ TILs. In *Tet2^fl/fl^* CD8^+^ TILs, there was an absence of TCR overlap between the progenitor-like states (*Ly6c2*^high^ T_SLC_ [T2] and T_PROG_ [T3]) and cytolytic/effector states (T_EFF-1_ [T4] and T_EFF-2_ [T5]), indicating a restricted clonal relationship (Figure 4K). Conversely, *Tet2KO* CD8^+^ TILs exhibited robust TCR overlap among these populations (Figure 4K). Collectively, these data demonstrate that *Tet2* deficiency prevents the oligoclonal expansion of terminally exhausted clones, while preserving a broad and diverse TCR repertoire across the progenitor and effector states.

### *Tet2* deficiency cell-intrinsically restrains terminal exhaustion and enables sustained effector generation via progenitor preservation

To directly assess whether *Tet2* regulates T-cell exhaustion in a cell-intrinsic manner, we utilized an *in vitro* chronic TCR stimulation model using purified CD8 T cells from *Cd4^Cre^Tet2KO* and *Tet2^fl/fl^* mice. Following initial activation with plate-bound anti-CD3ε and soluble anti-CD28 antibodies, cells were repeatedly re-stimulated with anti-CD3ε antibody alone to mimic persistent antigen exposure (Figure 5A).

**Figure 5.**
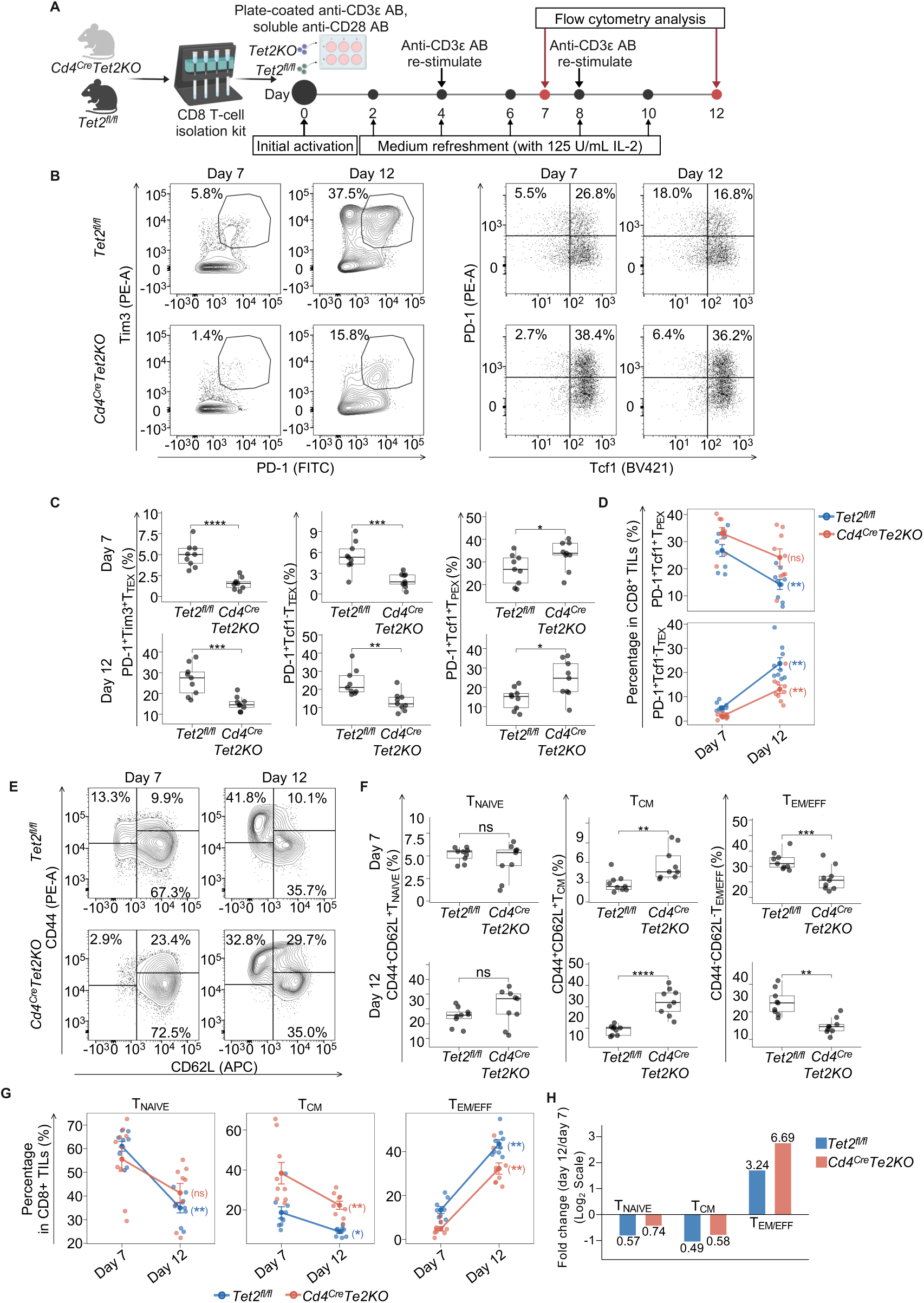
*Tet2* deficiency cell-intrinsically restrains terminal exhaustion while preserving self-renewing progenitor pools under chronic TCR stimulation. (A) Experimental design for *in vitro* chronic TCR stimulation assay. CD8 T cells isolated from *Cd4^Cre^Tet2KO* and *Tet2^fl/fl^* mice (*n =* 9 per group) were activated with anti-CD3ε (2 µg/mL) and soluble anti-CD28 (2 µg/mL) antibodies, then chronically re-stimulated with anti-CD3ε antibody alone (2 µg/mL) on days 4 and 8 to model persistent antigen exposure. Cells were analyzed by flow cytometry at days 7 and 12. (B) Representative flow cytometry plots showing expression of exhaustion markers PD-1 and Tim3 (left), and PD-1 and Tcf1 (right) at days 7 and 12 in *Tet2^fl/fl^* (top row) and *Tet2KO* (bottom row) CD8 T cells. (C) Quantification of exhaustion phenotypes. Box plots comparing frequencies of PD-1^+^Tim3^+^T_TEX_, PD-1^+^Tcf1^-^T_TEX_, and PD-1^+^Tcf1^+^T_PEX_ between *Tet2KO* and *Tet2^fl/fl^* groups at day 7 (top) and day 12 (bottom). See also Table S19A and S19B for detailed quantification and statistical comparisons. (D) Temporal dynamics of exhaustion differentiation. Longitudinal plots showing longitudinal changes from day 7 to day 12 of PD-1^+^Tcf1^+^T_PEX_ (top), and PD-1^+^Tcf1^-^T_TEX_ (bottom). See also Table S19A and S19C for detailed quantification and statistical comparisons. (E) Representative flow cytometry plots showing CD62L and CD44 expression patterns at days 7 and 12 in *Tet2^fl/fl^* (top row) and *Tet2KO* (bottom row) CD8 T cells. (F) Quantification of memory and effector differentiation states. Box plots comparing frequencies of CD44^-^CD62L^+^T_NAIVE_, CD44^+^CD62L^+^T_CM_, and CD44^+^CD62L^-^T_EM/EFF_ between *Tet2KO* and *Tet2^fl/fl^* groups at day 7 and day 12. See also Table S19A and S19B for detailed quantification and statistical comparisons. (G) Temporal dynamics of T_NAIVE_, T_CM_, and T_EM/EFF_ populations. Longitudinal plots show longitudinal changes from day 7 to day 12 of the indicated populations (left to right). See also Table S19A and Table S19C for detailed quantification and statistical comparisons. (H) Fold changes (day 12/day 7) on log2 scale quantifying differentiation kinetics. Numbers on bars indicate actual fold change values; y-axis is log2-transformed to show proportional changes. TCR, T-cell receptor; T_TEX_, terminally exhausted T cells; T_PEX_, progenitor exhausted T cells; T_CM_, central memory T cells; T_EM/EFF_, effector memory/ effector T cells; T_NAIVE_, Naïve T cells; AB, antibody. Data in (C), (D), (F), and (G) are compiled from three independent experiments (total *n =* 9 mice per group). Statistical significance for temporal comparisons (Day 7 vs. Day 12 within each genotype) determined by Wilcoxon signed-rank test (paired, two-tailed). Statistical significance for group comparisons (*Tet2^fl/fl^*vs. *Tet2KO* at each timepoint) determined by Mann-Whitney U test (unpaired, two-tailed). \**p <* 0.05, \*\**p <* 0.01, \*\*\**p <* 0.001, \*\*\*\**p <* 0.0001, ns = not significant.

Flow cytometric analysis at days 7 and 12 revealed that *Tet2* deficiency selectively altered exhaustion fate decisions (Figures 5B and 5C). *Tet2KO* CD8 T cells displayed significantly lower frequencies of PD-1^+^Tim3^+^T_TEX_ and PD-1^+^Tcf1^-^T_TEX_, accompanied by a significant enrichment of PD-1^+^Tcf1^+^ PD-1^+^ progenitor exhausted T (T_PEX_) cells at both timepoints (Figures 5B and 5C, Table S19A and S19B). Notably, longitudinal analysis further demonstrated that *Tet2* is required for progressive depletion of the progenitor exhausted compartment. Between days 7 and 12, the PD-1^+^Tcf1^+^T_PEX_ pool was markedly reduced in *Tet2^fl/fl^* cultures, whereas this population was maintained in *Tet2KO* CD8 T cells, despite accelerated T_TEX_-cell accumulation in both genotypes (Figure 5D; Table S19C). These findings validate our scRNA-seq-based observations and establish *Tet2* as a cell-intrinsic regulator that promotes terminal exhaustion commitment at the expense of the progenitor population under chronic TCR stimulation.

We next examined how *Tet2* deficiency influences memory and effector differentiation states during prolonged TCR signaling. Consistent with their exhaustion profile, *Tet2^fl/fl^* CD8 T cells exhibited significantly higher proportions of CD44^+^CD62L^-^ effector/effector memory (T_EM/EFF_) cells, whereas *Tet2KO* CD8 T cells retained significantly higher frequencies of CD44^+^CD62L^+^ central memory-like (T_CM_) cells at both days 7 and 12 (Figures 5E and 5F; Table S19A and S19B). Notably, although *Tet2KO* cells displayed lower T_EM/EFF_ proportions at individual timepoints, longitudinal analysis revealed robust differentiation capacity. From day 7 to day 12, *Tet2KO* cultures exhibited a 6.69-fold expansion of the T_EM/EFF_ compartment, exceeding the 3.24-fold increase observed in *Tet2^fl/fl^*cells (Figures 5G and 5H). Importantly, enhanced effector output in *Tet2KO* cells was achieved without excessive attrition of self-renewing populations. *Tet2KO* naive CD8 T cells (T_NAIVE_) were not significantly depleted over time, and while T_CM_ frequencies declined in both genotypes, *Tet2KO* cells exhibited lower fold reductions, indicating superior preservation of self-renewing memory compartments (Figures 5G and 5H; Table S19C). Together, these data demonstrate that *Tet2* deficiency decouples effector generation from progenitor depletion, enabling sustained differentiation while preserving Tcf1^+^T_PEX_ and CD62L^+^T_CM_-like reservoirs under chronic TCR stimulation.

### *Tet2* loss induces genome-wide hypermethylation and epigenetic silencing of the exhaustion regulator *Tox* in PD-1^+^CD8^+^ TILs

To determine whether *Tet2* loss impairs active DNA demethylation in CD8^+^ TILs, we performed whole genome bisulfite sequencing (WGBS) on activated PD-1^+^CD8^+^ TILs isolated from liver metastatic tumors in *Vav1^Cre^Tet2KO* and control *Tet2^fl/fl^* mice (Figure 6A). WGBS analysis comparing *Tet2KO* to *Tet2^fl/fl^*PD-1^+^CD8^+^ TILs identified 19,492 differentially methylated regions (DMRs) across all chromosomes, of which 98.8% were hypermethylated in *Tet2KO* cells (Figures S8A-C). Among these, 5,751 DMRs were mapped to annotated genomic loci corresponding to 4,260 genes (Figure S8D; Table S20). A total of 2,644 DMRs were located in regulatory elements, including enhancer, promoter, transcription start site (TSS), or CTCF-binding regions, and were designated as regulatory DMRs (rDMRs) (Figure 6B; Table S21). In addition, the majority of rDMRs were distributed within the enhancer or promoter regions (Figure 6B). Notably, *Tet2KO* PD-1^+^CD8^+^ TILs exhibited a marked predominance of hypermethylated rDMRs (hyper-rDMRs; *n =* 2,519) compared with hypomethylated rDMRs (hypo-rDMRs; *n =* 125) (Figure 6C). Integration of WGBS with scRNA-seq revealed 22 genes with predominantly hypermethylated rDMRs that were also transcriptionally downregulated in *Tet2KO* PD-1⁺CD8^+^ TILs compared to *Tet2^fl/fl^*PD-1⁺CD8^+^ TILs (Figures 6D; Table S22). Among these, *Tox*, a master regulator of T-cell exhaustion, showed one of the most extensive methylation changes genome-wide (Figures 6D and S8E). A methylation impact score (total DMR count × mean absolute methylation difference) (see Methods) similarly ranked *Tox* second and highlighted its widespread hypermethylation across multiple regulatory regions (Figure S8F).

**Figure 6.**
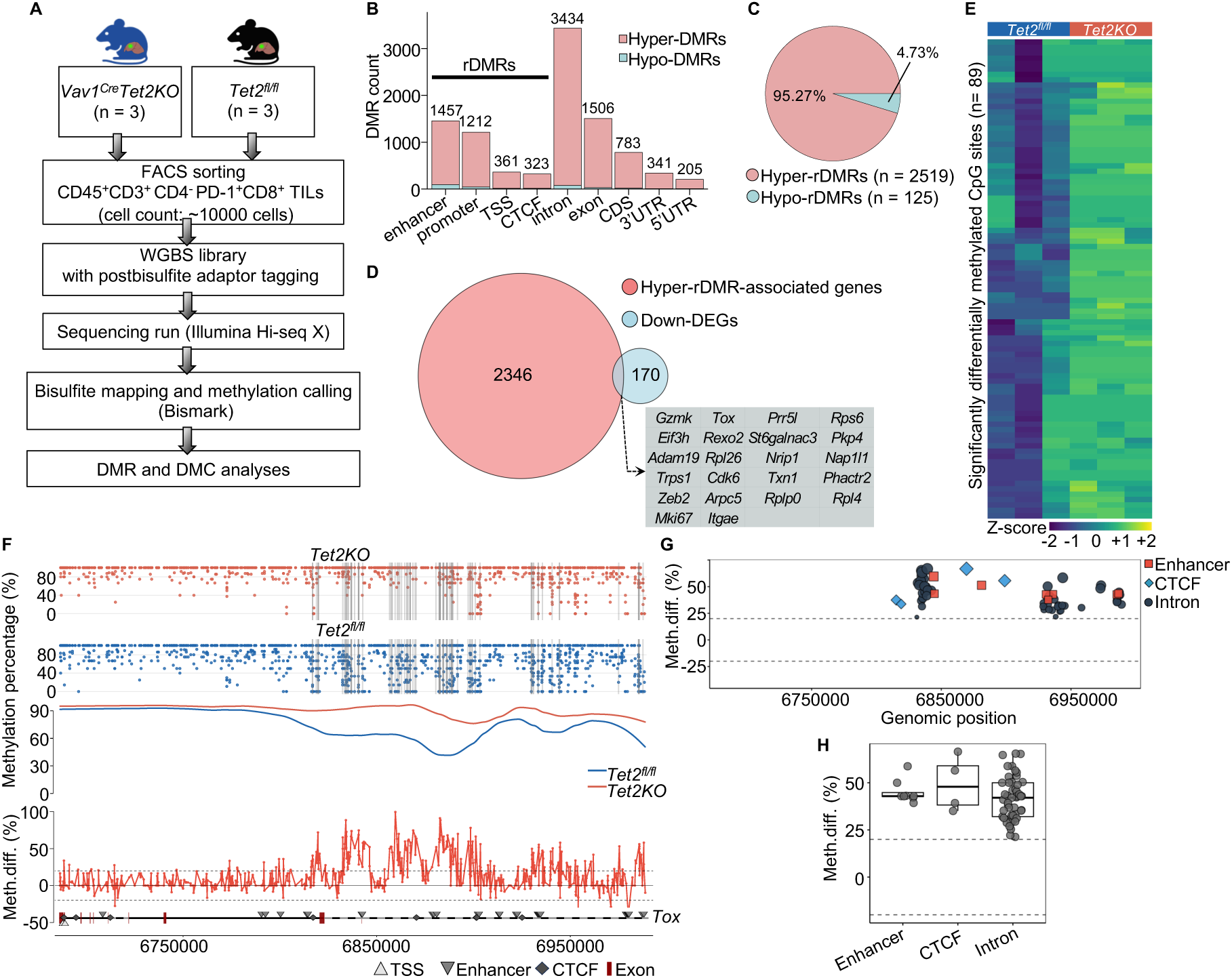
*Tet2* loss induces genome-wide hypermethylation and epigenetic silencing of *Tox* in PD-1⁺CD8^+^ TILs. (A) Schematic overview of WGBS workflow for analyzing DNA methylation in PD-1⁺CD8^+^ TILs. (B) Distribution of DMRs across genomic features in *Tet2KO* vs. *Tet2^fl/fl^* PD-1^+^CD8^+^ TILs. DMRs located at enhancer, promoter, TSS, or CTCF-binding regions, and were designated as regulatory DMRs (rDMRs). (C) Pie chart displaying the proportion of hyper-rDMRs (*n =* 2519, 95.3%) versus hypo-rDMRs (*n =* 125, 4.7%). (D) Venn diagram identifying 22 genes showing coordinated hypermethylation (*q-*value < 0.01, meth.diff > 0.2) and transcriptional downregulation (min.pct = 0.1, adjusted *P* value < 0.05) in *Tet2KO* PD-1^+^CD8^+^ TILs, including the master exhaustion regulator *Tox*. (E) Heatmap displaying 89 significant DMCs across the *Tox* locus identified by DMC analysis, with Z-scores indicating methylation levels in *Tet2^fl/fl^* and *Tet2KO* replications. (F) Genome browser view of the *Tox* locus at CpG resolution showing methylation percentage at 735 CpG sites in *Tet2^fl/fl^* (blue) and *Tet2KO* (red) PD-1^+^CD8^+^ TILs, with smoothed methylation trends and methylation difference plot below, the grey bars highlighting 89 significant DMCs. Gene structure indicates TSS, enhancers, CTCF binding sites, and exons. (G) Scatter plot of methylation percentage differences across the *Tox* gene structure by DMR analysis, displaying 56 rDMRs at enhancer, CTCF, and intronic regions. (H) Box plots of methylation percentage differences at enhancer, CTCF, and intronic regions within the *Tox* locus. TIL, tumor-infiltrating lymphocytes; WGBS, whole genome bisulfite sequencing; DMR, differentially methylated region; DMC, differentially methylated cytosine; TSS, transcription start site; CTCF, CCCTC-binding factor; CDS, coding sequence; UTR, untranslated region; min.pct, minimum percentage of cells. For (B)-(H), statistical significance to identify DMRs and DMCs determined by *methylKit* using *q-*value < 0.01 and |meth.diff| ≥ 20%.

Within the *Tox* locus, differentially methylated CpG site (DMC) analysis identified 89 DMCs among 735 CpGs, all showing increased methylation in *Tet2KO* PD-1^+^CD8^+^ TILs relative to *Tet2^fl/fl^* PD-1^+^CD8^+^ TILs, with a mean gain of 58.2% (Figures 6E, 6F and S8G-S8I; Table S23). DMR analysis focusing on the *Tox* locus using 500bp-window screening revealed 56 significant DMRs with a mean methylation increase of 47.2% (Figures 6G and S8J; Table S24). Notably, *Tox* showed extensive hypermethylation at regulatory elements, including 8 enhancer DMRs with mean methylation difference of 45.2%, and 4 CTCF binding site DMRs with mean methylation difference of 49.3% (Figures 6G and 6H). These findings indicate that *Tet2* deficiency induces broad hypermethylation at critical regulatory elements of *Tox*, leading to transcriptional repression of this central exhaustion-program regulator in *Tet2KO* PD-1⁺CD8^+^ TILs.

### *Tet2KO:Tet2WT* chimeric mouse model of CH demonstrates restricted tumor growth and intrinsic resistance of *Tet2-*deficient CD8^+^ TILs to terminal exhaustion

To model the mosaicism of tumor-infiltrating immune cells observed in solid tumor patients with CH, in which mutant and wild-type hematopoietic cells coexist within the tumor microenvironment, we established a competitive bone marrow transplantation (BMT) mouse model of CRC liver metastasis (Figure 7A). Lethally irradiated CD45.1⁺ wild-type (WT) recipients were reconstituted with mixtures of *Vav1^Cre^⁺Tet2^fl/fl^* CD45.2⁺ BM cells and *Tet2WT* CD45.1⁺/CD45.2⁺ competitor BM cells at ratios of 1:2 or 1:3, generating *Tet2KO:Tet2WT* mixed chimeras. Control *Tet2^fl/fl^:Tet2WT* chimeras received a 1:1 mixture of *Tet2^fl/fl^* CD45.2⁺ and *Tet2WT* CD45.1⁺/CD45.2⁺ BM cells. Four weeks post-BMT, flow cytometric analysis of peripheral blood confirmed successful hematopoietic reconstitution, with donor-derived CD45.2⁺ cells constituting >95% of B220⁺ B lymphocytes, CD11b⁺ myeloid cells, and NK1.1⁺ natural killer cells (Figure S9A). As expected from radio-resistant host T cell survival^48^, approximately 25.9 ± 1.8% recipient-derived CD45.1⁺ T cells remained within the CD3⁺ T cell population (Figure S9A). Subsequently, Venus-*AKTP^M/LOH^* CRC cells were injected intrasplenically to induce liver metastases in these reconstituted chimeric mice (Figure 7A).

**Figure 7.**
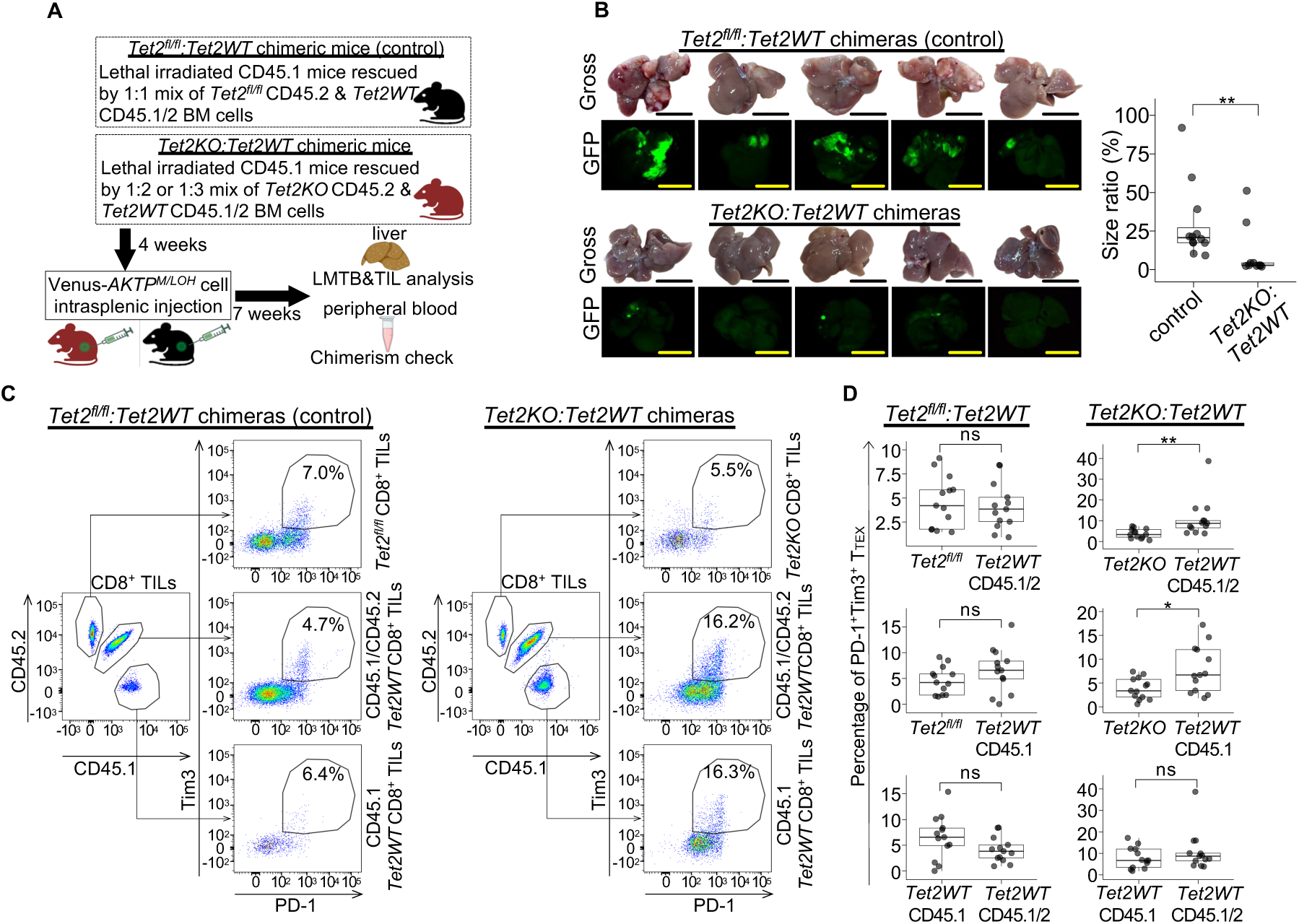
*Tet2KO:Tet2WT* chimeric mouse model of CH demonstrates restricted tumor growth and intrinsic resistance of *Tet2-*deficient CD8^+^ TILs to terminal exhaustion in CRC liver metastasis. (A) Experiment schema of *Tet2KO:Tet2WT* chimeric mouse model with CRC liver metastasis. Lethally irradiated CD45.1⁺ WT-recipients were reconstituted with mixtures of *Vav1^Cre^⁺Tet2^fl/fl^* CD45.2⁺ and *Tet2WT* CD45.1⁺/CD45.2⁺ BM cells at ratios of 1:2 or 1:3 (KO : WT competitor) to generate *Tet2KO:Tet2WT* mixed chimeras. Control *Tet2^fl/fl^*:*Tet2WT* chimeras received a 1:1 mixture of *Tet2^fl/fl^* CD45.2⁺ and *Tet2WT* CD45.1⁺/CD45.2⁺ BM cells. Four weeks post-transplantation, Venus-*AKTP^M/LOH^* colorectal cancer cells were intrasplenically injected. Chimerism was assessed in peripheral blood 8 weeks post-injection, followed by analysis of CD8^+^ TILs and LMTB. (B) Representative liver metastasis samples and box plot comparing LMTB between *Tet2KO:Tet2WT* chimeric mice (*n =* 10) and *Tet2^fl/fl^*:*Tet2WT* controls (*n =* 12) (29.1 ± 24.1% vs. 10.6 ± 16.7%; *p =* 0.0076). Scale bars: 1 cm. (C) Representative flow cytometry plots showing the proportion of PD-1⁺Tim3⁺ CD8 T_TEX_ in *Tet2^fl/fl^* CD45.2⁺, host-derived *Tet2WT* CD45.1⁺, and *Tet2KO* CD45.2⁺ CD8^+^ TILs. (D) Box plots comparing frequencies of PD-1⁺Tim3⁺ CD8 T_TEX_ across donor and host CD8^+^ TIL populations (*n* = 13 per group). See also Table S25D for detailed quantification and statistical comparisons. CH, clonal hematopoiesis; TILs, tumor-infiltrating lymphocytes; CRC, colorectal cancer; BM, bone marrow; LMTB, liver metastatic tumor burden; T_TEX_, terminally exhausted T cells. Data in (B) and (D) are compiled of at least 2 independent experiments and presented as mean ± SD. Statistical significance was determined by Mann-Whitney U test. \**p <* 0.05, \*\**p <* 0.01, \*\*\**p <* 0.001, \*\*\*\**p <* 0.0001, ns = not significant.

Twelve weeks post-BMT, *Tet2KO* cells exhibited progressive competitive expansion relative to *Tet2WT* cells across all immune cell subsets compared with the four-week post-BMT baseline, with the most pronounced expansion in myeloid cell populations (Figure S9B; Table S25A and S25B). In contrast, control chimeras showed no competitive advantage of *Tet2^fl/fl^* cells over *Tet2WT* competitor cells, maintaining stable ratios throughout the observation period (Figure S9B; Table S25A and S25B). This clonal expansion pattern recapitulates the progressive outgrowth of somatic *TET2* mutant-derived CH observed in human patients^49^. Within liver metastases of *Tet2KO:Tet2WT* chimeras, *Tet2KO* T cells accounted for approximately 35–46% of total T cells and 23–32% of the CD8^+^ TIL subset. These frequencies were comparable to the *Tet2^fl/fl^* CD45.2⁺ populations observed in controls (Figure S9C; Table S25C).

Consistent with the restricted LMTB observed in *Vav1^Cre^/Cd4^Cre^Tet2KO* mice, *Tet2KO:Tet2WT* chimeric mice demonstrated significantly reduced LMTB compared to *Tet2^fl/fl^:Tet2WT* controls (Figures 7B and S9D). Flow cytometric analysis of liver TILs revealed a markedly higher proportion of PD-1⁺Tim3⁺ T_TEX_ in both *Tet2^fl/fl^* CD45.2⁺ and residual recipient *Tet2WT* CD45.1⁺ populations compared to that in *Tet2KO* CD45.2⁺ CD8^+^ TILs (Figures 7C and 7D; Table S25D). Collectively, these data demonstrate that even within a competitive environment containing *Tet2WT* counterparts, *Tet2*-deficient CD8^+^ TILs exhibit an intrinsic resistance to terminal exhaustion.

### Anti-PD-1 treatment suppresses CRC liver metastasis in Tet2^fl/fl^ mice

Given the enrichment of exhaustion markers in *Tet2^fl/fl^*CD8^+^ TILs, we evaluated the therapeutic efficacy of PD-1 blockade (Figure S10A). *Tet2^fl/fl^* mice treated with anti-PD-1 antibody exhibited complete tumor eradication, whereas IgG-treated controls developed profound liver metastatic growth (Figures S10B and S10C). Although IgG-treated *Vav1^Cre^Tet2KO* mice displayed reduced LMTB compared to *Tet2^fl/fl^* controls (IgG-treated *Vav1^Cre^Tet2KO* vs. IgG-treated *Tet2^fl/fl^*, 0.54 ± 0.39 mm2 vs. 23.53 ± 12.64 mm2, *p =* 0.008), minimal residual tumors persisted (Figures S10B and S10C). This indicates that targeting the PD-1 axis is sufficient to reverse the metastasis-permissive environment associated with *Tet2-*intact hematopoiesis.

## DISCUSSION

While CH is highly prevalent among patients with solid tumors, its impact on fundamental aspects of cancer biology, including tumor growth, treatment response, and metastasis, has remained poorly defined^21,26,50,51^. In this study, we demonstrate that *TET2*-mutant CH confers protection against cancer metastasis, extending prior observations that *TET2* inactivation can enhance anti-tumor immunity in settings such as chimeric antigen receptor T-cell therapy^52,53^ and immune checkpoint blockade^15,33^. By integrating large-scale clinical data with genetic and functional mouse models, we identify *TET2*-mutant CH as a previously unrecognized tumor-suppressive state that limits metastatic outgrowth.

In a cohort of over 16,000 solid tumor patients, we show that individuals with *TET2*-driver clonal hematopoiesis have significantly lower odds of presenting with metastatic disease and exhibit reduced metastatic burden. Notably, although *TET2*-mutant CH is generally considered an adverse condition due to its association with increased morbidity and mortality from atherosclerotic and fibrotic diseases^54–57^, we did not observe a difference in median survival between solid tumor patients without CH and those with *TET2*-CH. This finding stands in contrast to patients harboring CH driven by mutations in DNA damage response, cohesin, or spliceosome genes, who experienced significantly worse overall survival, consistent with prior studies^22,23,27,58^. Together, these data suggest that the biological consequences of CH are highly driver dependent, and that *TET2*-mutant CH differs fundamentally from other high-risk CH genotypes in the context of solid tumor microenvironments.

Our liver metastasis models provide functional evidence supporting a causal relationship between *TET2*-mutant CH and protection from metastatic disease. Both complete and chimeric pan-hematopoietic *Tet2* inactivation were sufficient to attenuate metastatic outgrowth, demonstrating that this association is not merely correlative. Importantly, we show that *Tet2*-deficient CD8 T cells are necessary and sufficient for the anti-metastatic effect of *TET2*-mutant CH, linking this phenotype directly to adaptive immune function. This finding helps reconcile prior heterogeneous reports on the impact of *Tet2*-mutant CH in syngeneic tumor models, where tumor growth has been reported as promoted, suppressed, or unaffected depending on tumor type, implantation site, and experimental context^29,30,59^. Indeed, our patient-level analyses revealed tumor-type–specific variation, with the strongest protective association observed for liver and brain metastasis.

Mechanistically, our data established that *Tet2* deficiency restrains metastatic progression by limiting terminal CD8 T cell exhaustion. Tet2 is transcriptionally induced following T cell receptor engagement^53,60^ and has been shown to act as a negative regulator of adaptive immunity by catalyzing DNA demethylation at key regulatory loci, thereby favoring differentiation toward exhausted or regulatory states at the expense of durable memory^60,61^. In the context of CH, this negative feedback is diminished. Within metastatic liver lesions, *Tet2*-deficient CD8^+^ TILs exhibit sustained methylation at regulatory elements of the *Tox* locus, resulting in transcriptional repression of TOX, a master regulator of terminal exhaustion^62,63^. This epigenetic constraint preserves progenitor and stem-like CD8 T cell states and enables sustained effector generation under chronic antigen stimulation.

These findings define a TET2–DNA methylation–TOX axis as a central epigenetic checkpoint governing exhaustion fate commitment in CD8 T cells. Similar mechanisms have been implicated in enhanced control of chronic infection and cancer by *Tet2*-mutant CD8 T cells and *Tet2*-deficient CAR-T cells^53^, underscoring the generalizability of this regulatory pathway. However, unrestrained T-cell activation associated with *TET2*-mutant CH may also be pathological in certain contexts, including autoimmunity^64,65^, immune-related toxicities during immunotherapy^66^, or malignant transformation following acquisition of additional mutations^67,68^. Indeed, *TET2*-mutant CH is a recognized risk factor for T cell lymphoma^69^, highlighting the dual-edged nature of sustained immune activation.

In addition to its effects on T cells, *Tet2* loss has been shown to enhance antigen-presenting capacity in macrophages^15^. While myeloid-restricted *Tet2* inactivation was insufficient to suppress liver metastasis in our models, it may still contribute to improved tumor control in concert with CD8 T cells. Consistent with this notion, we and others have previously observed transcriptional signatures of heightened T-cell activation in colorectal cancer and melanoma samples from patients with *TET2*-mutant CH^15,70^. Defining the relative contributions of distinct immune lineages to other CH-associated clinical phenotypes remains an important area for future investigation.

Several limitations of this study should be acknowledged. We were unable to directly assess whether tumor-infiltrating *TET2*-mutant T lymphocytes correlate with metastatic status in patients, as such data were not available. Although *TET2* is a well-established negative regulator of CD8 T cell activity^53,60,71^, the frequency and functional relevance of *TET2*-mutant CD8 T cells within human tumors remain incompletely characterized. Moreover, our experimental models focused on experimental liver metastasis and therefore do not address potential effects of *TET2*-mutant CH on primary tumor growth, metastatic dissemination, or circulation through blood and lymphatic compartments. While the *Cd4Cre* model demonstrates that T cells are necessary and sufficient for the observed phenotype, the potential contribution of other *Tet2*-deficient immune cells warrants further study. Finally, the causes of death in solid tumor patients with high-risk CH are not known and may be unrelated to metastatic disease.

In summary, we show that *TET2*-mutant CH suppresses cancer metastasis in both patients and animal models by enforcing an epigenetic barrier to terminal CD8 T-cell exhaustion. These findings reposition *TET2*-mutant CH as a protective modifier of tumor progression and suggest that CH status may have utility as a prognostic biomarker for metastatic risk, at least in certain cancers. Notably, the magnitude of metastatic suppression conferred by *TET2*-mutant CH was substantial in our experimental models, approaching the level of tumor control observed with immune checkpoint blockade, suggesting that pharmacologic strategies designed to mimic hematopoietic *TET2* inactivation—or to target downstream components of the TET2–TOX axis—could represent novel therapeutic approaches for the prevention or treatment of metastatic disease.

## METHODS

### Experimental model and subject details Human study cohort

Patients who were tested for clonal hematopoiesis using an MSK-IMPACT targeted sequencing panel with somatic mutations from peripheral blood sequencing results curated as putative drivers or non-drivers of CH in the Bolton *et al*. 2020 cohort who were also evaluated for metastatic disease by Nyugen *et al*. 2022 were included in our sub-cohort analysis. Study data containing anonymous patient identifiers were downloaded from cBioPortal on September 4^th^, 2025. Using variants reported by Bolton *et al.* 2020, clonal hematopoiesis (CH) was called as a putative driver mutation with variant allele fraction (VAF) ≥ 2%. Each mutation was defined as either putative (PD) or non-putative based on its role in cancer pathogenesis and recurrence in an MSKCC in-house dataset of myeloid neoplasms as in Bolton *et al.* 2020. It was further was defined as either a non-myeloid PD or myeloid PD based on relevance to myeloid neoplasms specifically.

### Mice

*Vav1^Cre36^*, *Cd4^Cre37^*, *Cd19^Cre38^*, and *LysM^Cre39^* transgenic mice were crossed with *Tet2^fl/fl^*mice^35^ to generate homozygous *Tet2*-knockout mice (*Vav1^Cre^Tet2KO, Cd4^Cre^Tet2KO*, and *LysM^Cre^Tet2KO*). *Tet2^fl/fl^* mice served as *Tet2* wild-type control. All mice were maintained in a specific pathogen-free animal facility in the Laboratory of Animal Center, University of Tsukuba. All animal procedures adhered to the The Regulations for Animal Experiments in University of Tsukuba. Mouse genotyping was performed using genomic DNA extracted either from tails by KAPA kit method (KAPA Biosystems) or from blood by QIAamp DNA Blood Mini Kit (Qiagen). The primers used for genotyping are listed in Table S26.

### Organoid cell line

Venus-labeled mouse intestinal tumor-derived organoid-originated cells simultaneously carrying *Apc^Δ716^*, *Kras^+/G12D^*, *Tgfbr2^-/-^*, and *Trp53^R270H/LOH^* mutations (Venus-*AKTP^M/LOH^* cells) were established as previously described^40^. Venus-*AKTP^M/LOH^* cells were cultured in Advanced DMEM/F-12 medium (Gibco, #2508872) supplemented with 10% Fetal Bovine Serum (FBS, Hyclone, #AE25787263), 1% Penicillin-Streptomycin (PS, Fujifilm, #168-23191), 0.3 μg/ml Rock inhibitor (Y27632, Fujifilm, #235-00513), 0.9 μg/ml GSK-3 inhibitor (CHIR99021, R&D system, #4423), and 21 μg/ml ALK inhibitor (A8301, R&D systems, #5640). For the passage, Venus-*AKTP^M/LOH^* cells were enzymatically dissociated with 0.25% Trypsin at 37°C for 10 minutes and mechanically dissociated by pipetting.

### Method details

#### Clinical cohort data processing and statistical analysis

##### (1) Clonal hematopoiesis mutation grouping

Patients were categorized as having CH if they had a variant annotated as CH PD in Bolton *et al.* 2020^22^. Patients were then categorized as having a *TET2* or *DNMT3A* PD only if they had no other CH-defining PD variants. Using a mutually exclusive hierarchical system, patients were first categorized as having a *TET2* driver, followed by a *DNMT3A*, cohesin (*STAG2, RAD21, SMC1, SMC3, PDS5A, PDS5B*), spliceosome (*SF3B1, SRSF2, ZRSR2, U2AF1*), or DNA-damage-response driver (*PPM1D, CHEK2, TP53, ATM*). For example, if a patient had both a cohesin (e.g. *STAG2*) and spliceosome (*SF3B1*) PD variant, they would be categorized in the cohesin group. A patient with a *TET2* and *STAG2* PD variant would also be categorized in the cohesin group.

##### (2) Quantification of cancer metastatic burden

Patients were labelled as metastatic patient or non-metastatic patient by Nguyen *et al*.. For each patient with metastatic disease, the number of metastases and metastatic sites was quantified. Patients were assigned as positive or negative for cancer metastases to various organs including lung, liver, draining lymph node (DLN), brain, and bone.

##### (3) Clinical cohort statistical analysis

The odds of metastatic disease stratified by each PD variant group was assessed using multivariate logistic models to account for covariates that have previously been linked either to CH prevalence or cancer outcome, including chemotherapy and radiation treatment status, age, sex, smoking status, race, and cancer type. Similar models were used to assess odds of metastasis within cancer types for *TET2*-driver CH. However, due to few occurrences of non-metastatic cancer in *TET2*-driver CRC patients and prostate cancer patients and limitations of multivariate analysis, Fisher’s exact test for count data was applied in addition to the logistic model to ensure robust statistical testing. (Table S1). For odds of metastasis to specific sites, one model for each metastatic site allowed for patients with metastases at multiple sites (such as brain and bone) to be classified as positive for all metastatic sites.

To assess metastatic burden, Wilcoxon rank-sum tests were used to quantify differences in distribution of both number of metastases, and number of metastatic sites. Further, multivariate negative binomial generalized linear models were used to account for the same covariates as above.

Patients were categorized in a mutually exclusive manner according to the criteria outlined above to investigate associations between PD variant groups and survival. Kaplan-Meier survival curves and a corresponding Cox proportional hazards model were used to account for covariates previously linked to survival, including age, sex, race, cancer type, smoking status, chemotherapy and radiation treatment status, and metastatic status.

Difference in distribution of age between CH driver patients and non-CH driver patients was quantified using a multivariate linear model to account for covariates associated with age, including age, sex, race, cancer type, smoking status, chemotherapy and radiation treatment status, and metastatic status. The association between *ASXL1* CH driver mutation prevalence and smoking status was assessed using a multivariate logistic model, as were the associations between DNA-damage-response CH driver mutation prevalence with treatment status, and metastatic disease with treatment status. Odds of metastatic disease in patients with a *TET2* CH driver mutation as opposed to a *TET2* CH non-driver mutation was also assessed using a multivariate logistic model.

All statistical tests are two-tailed, and analyses in which multiple comparisons were made were corrected for multiple hypothesis testing in a false discovery rate (FDR) controlled fashion. An FDR-controlled *P* value threshold of 0.05 was used to denote significance.

#### Venus-AKTP^M/LOH^ cell injection, liver metastasis tumor burden (LMTB) evaluation

Venus-*AKTP^M/LOH^* cells were used for the mouse spleen injection after at least four times of passage. One million (1×10^6^) of thoroughly dissociated Venus-*AKTP^M/LOH^* cells resolved in 30 μl of Matrigel Matrix (Corning) were injected into mouse spleen. Mouse livers were harvested 30 days post-injection and imaged as whole organs using the fluorescence stereo microscope equipped with the digital color microscope camera (M165 FC/DFC450C, Leica Microsystems) at 4.6x magnification to macroscopically assess the presence of Venus-*AKTP^M/LOH^* cell metastasis. LMTB was evaluated as previously described^72^. Briefly, formalin-fixed paraffin-embedded liver sections were stained with hematoxylin and eosin (H&E). The tumor burden was quantified by calculating the ratio of the total tumor area to the total liver section area using a Keyence BZ-X710 microscope and BZ-X analyzer software The two-tailed Mann-Whitney U test with α = 0.05 was used to compare the size ratio between the two groups.

#### Antibody-based depletion of CD8 T cells

Anti-CD8α antibody treatment was initiated two days prior to intrasplenic injection of Venus-*AKTP^M/LOH^* cells. InVivoMAb anti-mouse CD8α antibody (clone 2.43, BioXCell, #BE0061) was diluted with Phosphate-Buffered Saline (PBS) to the concentration of 1μg/μL and intraperitoneally injected at an initial dose of 200 μg/mouse, followed by 6 booster injections of 100 μg/mouse every 4 days throughout the tumor progression course. Control mice received PBS in volumes matching those used in the anti-CD8α antibody treatment schedule. Mice were sacrificed 30 days after the injection of Venus-*AKTP^M/LOH^* cells to collect the whole liver for LMTB evaluation.

#### Enzymatic digestion of liver metastases and isolation of tumor-infiltrating immune cells for flow cytometry

Mouse livers were harvested 30 days post-injection. Liver metastatic tumors and surrounding liver tissues 0.5 cm from the edge of tumors were minced into small pieces < 1mm^3^. The minced tumor tissue was then digested by 15 ml of digestion buffer containing 0.75 mg/ml collagenase A (Sigma-Aldrich, #10103578001) and 50 μg/ml DNase I (Worthington Biochemical, #LS002139) in DMEM (Gibco, #10567-014), and incubated at 37 °C with shaking at 170 rpm for 30 mins. After tissue digestion, the cell suspension was first filtered through a 70-μm cell strainer (Falcon) to remove larger debris. The resulting homogenate was then further filtered through a 30-μm cell strainer (Miltenyi Biotec) to obtain a single-cell suspension. Any remaining undigested lumps were further dissociated by gentle pipetting. DMEM supplemented with 10% FBS (Hyclone, #AE25787263) and 1% Penicillin-Streptomycin (PS, Fujifilm, #168-23191) was added to terminate the digestion. The cell suspension was centrifuged at 163× *g* for 7 mins at 4 °C, and then the cell pellet was resuspended in 5 ml of 40% Percoll (Cytiva, #17089102) in PBS. The cell suspension was then slowly and carefully dropped on top of 3 ml of Histopaque-1083 (Sigma-Aldrich, #10831) without mixing and centrifuged at 1000× *g* for 20 mins in brake off mode. The cells concentrated at the interface layer were carefully aspirated and added into 30 ml of FACS buffer (PBS containing 2% Bovine Serum Albumin (BSA) and 2 mM EDTA) and centrifuged at 800× *g* for 5 mins at 4 °C. To detect exhausted T cells, the yielded cell pellet was resuspended in FACS buffer and incubated with a panel of fluorophore-conjugated anti-mouse antibodies consisting of anti-CD45, anti-CD3e, anti-CD4, anti-CD8α, anti-PD-1, and anti-Tim3 (Havcr2/CD366) antibodies (Table S27A) before being analyzed by BD FACSAria III (BD Biosciences).

#### Bulk-RNA sequencing

Mouse livers were harvested 30 days post-injection. Tumor-infiltrating leukocytes yielded by the same procedure as that used for flow cytometry were utilized for the CD45-positive leukocyte selection. 10 μL of CD45 microbeads (Miltenyi Biotec, #130-052-301) was mixed well and incubated with each 1×10^7^ cells in 90 μL of FACS buffer for 30mins at 4°C. The CD45-positive selection was then conducted using the MACS LS column and the MidiMACS separator (Miltenyi Biotec), following the column-based magnetic cell-separation strategy established by the manufacturer. CD4^+^ TILs and CD8^+^ TILs were identified using a panel of fluorophore-conjugated anti-mouse antibodies consisting of CD45, CD3e, CD4, and CD8α, and isolated by BD FACSAria III Cell Sorter (BD Biosciences).

Total RNA was isolated using RNeasy Micro Kit (Qiagen). RNA integrity number (RIN) values were measured using the Agilent 2100 Bioanalyzer (Agilent Technologies). Samples with RIN > 7 were subsequently used for library preparation using the SMARTer Stranded Total RNA-seq Kit v2 - Pico Input Mammalian (Takara Inc.) according to the manufacturer’s protocol. The library size distribution and concentration were assessed with the High Sensitivity DNA assay using the Agilent 2100 Bioanalyzer (Agilent Technologies). Sequencing was carried out using the Illumina HiSeq X platform (Macrogen) with a standard 150 bp paired-end read protocol. Subsequently, the gene expression was analyzed from the sequencing data using the CLC workbench version 11.0.1 (Qiagen) according the procedure described previously^73^. DEGs were identified by the Differential Expression for RNA-Seq tool of CLC workbench using the Wald test applying for the Against-control-group comparison with the *Tet2^fl/fl^* group serving as the control. The gene expression datasets were then used to identify significantly enriched gene sets for each group using GSEA^42^ (GSEA 4.3.0, https://www.gsea-msigdb.org/gsea/index.jsp) with the ImmuneSigDB subset of C7 (4872 gene sets, https://www.gsea-msigdb.org/gsea/msigdb/human/collections.jsp) for annotation.

#### Upstream regulator analysis

Overlapping downregulated DEGs within CD8^+^ TILs from *Vav1^Cre^Tet2KO* and *Cd4^Cre^Tet2KO* groups compared to the *Tet2^fl/fl^*group (FDR< 0.05, Log2 FC < -1; Table S8) were used to predict the upstream transcription factors by the Upstream Regulator Analysis function of QIAGEN’s IPA^74^ (QIAGEN Redwood City, www.qiagen.com/ingenuity). The pathway analysis was performed using the core analysis based on gene expression fold change^74^. The upstream transcription regulators were called with overlap p-value < 0.01. The extracted upstream regulators were then used to generate the molecular interaction network by the Network and Pathway tools of IPA.

#### Indirect immunofluorescence staining, co-expression cell evaluation

Liver metastasis tissue collected from *Vav1^Cre^Tet2KO, Cd4^Cre^Tet2KO* and *Tet2^fl/fl^* groups were mounted in O.C.T embeding compound (Sakura Finetek Japan Co.), frozen rapidly in hexane-dry ice and stored at -80°C. The frozen tissue blocks were then sectioned at 4 μm-thickness in a cryostat at -20°C. Sections were dried up in air for 60 mins at room temperature (RT), fixed in 4% paraformaldehyde for 15 mins and permeabilized in 0.1% Triton X100 for 10 mins and incubated with protein blocking buffer (Protein block, Abcam, #ab64226). Sections were then incubated with primary antibodies at 4°C for 16 hours, washed 3 times in PBS for 5 mins each and subsequently incubated with secondary antibodies for 30 mins at RT. Antibodies and the dilution used for immunofluorescence staining were provided in the Table S27D. After three washes, tissue sections were incubated with True VIEW Autofluorescence Quenching Kit (Vector Laboratories, #SP-8400-15) for 5 mins at RT and mounted with 4’,6-diamidino-2- 129 phenylindole (DAPI, Vectashield Mounting Medium; Vector Laboratories, #H-1200).

The sections were imaged using the confocal laser scanning microscope (Leica TCS SP8, Leica Microsystems). To quantify exhaustion phenotypes, CD8^+^ TILs co-expressing PD-1, Tim3, or Tox were manually counted in 10 to 20 random intratumoral fields per sample at 40× magnification. Data were expressed as the percentage of each subset relative to the total CD8^+^ TIL population. The one-way ANOVA test with α = 0.05 was used to compare the percentage of co-expressing cells between groups.

#### Single-cell RNA sequencing (scRNA-seq) library preparation, data acquisition and pre-processing

Liver samples from *Vav1^Cre^Tet2KO* mice and *Tet2^fl/fl^*mice were harvested at day 30 post-injection, and tumor-infiltrating leukocytes were harvested using the cell processing procedure that used for flow cytometry. The yielded cells were washed and resuspended in PBS supplemented with 1% BSA. Cells were then incubated with Brilliant Violet 421 (BV421) anti-mouse CD45.2 antibody (clone 104, Biolegend, #109832) for 20 minutes at 4°C, followed by staining with 7-AAD viability solution (ThermoFisher Scientific, 00-6993-50; 1:20) for 10 minutes on ice. CD45-positive live cells were subsequently sorted using the BD FACSAria III (BD Biosciences). Isolated CD45+ live cells were processed to generate barcoded 5′ gene expression (5′ GEX) libraries, together with T-cell receptor (TCR)-enriched libraries, using the Chromium Next GEM Single Cell V(D)J Reagent Kits v1.1 (10x Genomics, Pleasanton, CA, USA). The procedure followed the manufacturer’s instructions (CG000208 Rev G) and aimed to capture 10,000 cells per library. All libraries were subjected to quality 150 control checks and quantified using the 2100 Bioanalyzer High Sensitivity DNA Kit (Agilent 151 Technologies) and KAPA Library Quantification kit for Illumina platforms (Kapa Biosystems, KK4873). The library were sequenced on the Illumina NovaSeqX Plus Sequencing System (Macrogen), then mapped to the mouse genome (GRCm39) and demultiplexed using CellRanger (10X Genomics, v.8.0.1).

#### scRNA-seq data processing of individual samples

Matrix data of each samle was processed using Seurat^75^ (v.5.1.0) in RStudio (v.4.4.1). Doublets were detected and removed using the “scDblFinder” function from the scDblFinder package^76^ (v.1.18.0) with the default estimated doublet rate. Cells with fewer than 200 or more than 2,500 unique genes and those with over 5% of mitochondrial gene reads were excluded. After removing ribosomal and mitochondrial genes, gene expression count data was normalized using the “NormalizeData” function. Normalized data were linearly transformed (scaled) with the “ScaleData” function and subjected to principal component analysis (PCA) via the “RunPCA” function. Graph-based clustering was performed based on gene expression profiles using the “FindNeighbors” and “FindClusters” functions. Clusters were visualized using Uniform Manifold Approximation and Projection (UMAP) technique via the “RunUMAP” and “DimPlot” functions.

Cell clusters were annotated as T lymphocytes, B lymphocytes, NK cells, and myeloid cells based on marker genes identified by the “FindAllMarkers” and “FindConservedMarkers” functions. For a gene to be considered a marker, a minimum of 25% gene-expressing cells per cluster was required, and the minimum Log2 FC in gene expression of each cluster, compared to all other clusters, was set to 0.25.

#### Data integration for scRNA-seq and batch correction

Canonical correlation analysis (CCA) was performed to identify shared features across multiple gene expression datasets using the FindIntegrationAnchors function of Seurat. Four datasets (data from tumor-infiltrating leukocytes of *Tet2^fl/fl^* (*n =* 2) and *Vav1^Cre^Tet2KO* (*n =* 2) mice) were integrated based on the identified anchors using the IntegrateData function with 30 canonical correlation dimensions. The integrated data were scaled, subjected to PCA, and used for graph-based clustering for cell annotation, as described above.

#### scRNA-seq subclustering and cell-type annotation

CD8^+^ TIL clusters were identified and extracted from the integrated data, followed by scaling, PCA-based dimensionality reduction, and unsupervised graph-based subclustering for cell annotation. Subclusters that highly expressed marker genes characteristic of other immune cell subsets, distinct from CD8^+^ TILs, were excluded. Following in silico sorting, the CD8 T-cell dataset comprised 2,251 *Vav1^Cre^Tet2KO* CD8^+^ TILs and 1,101 *Tet2^fl/fl^* CD8^+^ TILs. To minimize discrepancies in cell numbers between the groups, downsampling was performed, resulting in a balanced dataset of 1,101 CD8^+^ TILs per group. The downsampling process was repeated multiple times to ensure consistency with the original data. Cell subclusters were annotated by analyzing the expression levels of canonical gene markers using Seurat’s “FindAllMarkers” and “FindConservedMarkers” functions.

#### DEG analysis of scRNA-seq

DEG analysis between *Tet2^fl/fl^* and *Vav1^Cre^Tet2KO* conditions was performed using “FindMarkers” function of Seurat. Genes were considered differentially expressed if they met the following criteria: (i) expressed in a minimum of 10% of cells per cluster within at least one condition, and (ii) exhibited a Benjamini–Hochberg adjusted p-value (adjusted p-value) less than 0.05.

#### Analysis of single-cell gene set activity

We used the gene set comprising of 42 consistently exhausted T cell associated marker genes identified by Zhang^45^ to calculate the T-cell exhaustion score for single cell CD8^+^ TIL dataset using “AddModuleScore” function. The number of bins of aggregate expression levels for all analyzed features was set to 24 and number of control features selected from the same bin per analyzed feature was set to 100. The differential enrichment of T-cell exhaustion scores between *Tet2^fl/fl^*and *Vav1^Cre^Tet2KO* groups was visualized using boxplots, and statistical significance was assessed via the Mann-Whitney U test.

We used the AUCell package^44^ v.1.26.0 to analyze the Memory versus Exhausted CD8 T-cell signature enrichment, and Effector versus Exhausted CD8 T-cell signature enrichment for CD8^+^ TIL dataset. The gene set “*GSE9650_EXHAUSTED_VS_MEMORY_CD8_TCELL_DN”* and “*GSE9650_EXHAUSTED_VS_MEMORY_CD8_TCELL_UP*” were obtained from the Human MSigDB (v2024.1.Hs) and converted to mouse orthologs using the ‘orthologs” function from the babelgene package (v.22.9). Gene expression rankings were calculated for each cell using the “AUCell_buildRankings” function, followed by computation of the area under the curve (AUC) scores of every cells for each gene set using the “AUCell_calcAUC” function. To create an aggregate Memory versus Exhausted CD8 T-cell signature, the AUC values of the upregulated gene set were subtracted from those of the downregulated gene set. Similarly, to calculate the aggregate Effector versus Exhausted CD8 T-cell signature enrichment, we subtracted the AUC values of the downregulated gene set from those of the upregulated gene set using the gene sets “GSE9650_EFFECTOR_VS_EXHAUSTED_CD8_TCELL_DN” and “GSE9650_EFFECTOR_VS_EXHAUSTED_CD8_TCELL_UP” obtained from the Human MSigDB (v2024.1.Hs). The differential enrichments of these gene signatures between *Tet2^fl/fl^*and *Vav1^Cre^Tet2KO* groups were visualized using boxplots, and statistical significance was assessed via the Mann-Whitney U test.

#### Single-cell trajectory inference

We used *Monocle 3 toolkit*^46^ (v.1.3.7) to construct the cellular pseudotime trajectories for CD8 T-cell subsets. Single cell datasets were preprocessed using PCA, retaining 50 dimensions. Dimensionality reduction was then performed using UMAP technique. Cells were clustered using the “cluster_cells” function with the Leiden algorithm, setting the resolution parameter to 0.008, the number of nearest neighbors (k) to 20, and performing two iterations. A principal graph representing cellular transitions was learned using the “learn_graph” function with default heuristics and without partitioning the data. According to previous knowledge, we selected the root cell cluster to order single cells along the trajectory using the “order_cells” function. Single cell pseudotime and trajectory were visualized using the “plot_cells” function.

#### TCR-seq data analysis

Single cell TCR-seq contig data from all samples were combined and annotated into clonotypes based on shared nucleotide sequences, amino acid sequences, and V(D)J gene segment usage using the “combineTCR” function of scRepertoire package^47^ (v.2.3.0). Clonotypes were defined using the combination of V(D)J genes (TCR/Ig) and the nucleotide sequence of the CDR3 region (strict clonotype calling) throughout the analysis. The clonotype data was integrated with gene expression datasets for clonal frequency calculation using the “combineExpression” function. TCR clonotype similarity across cell clusters was analyzed using the “clonalOverlap” function with the Morisita overlap index^47^.

Clonal diversity was quantified using Shannon entropy and the inverse Simpson index^77,78^. Clonal inequality was assessed using the Gini coefficient and Lorenz curve analysis^79^. All metrics were calculated from TCR clonotype frequencies using R (v.4.4.1) in RStudio.

#### Whole genome bisulfite sequencing (WGBS)

PD-1^+^CD8^+^ TILs isolated from the liver metastatic tumors of *Vav1^Cre^Tet2KO* (*n =* 3) and *Tet2^fl/fl^*(*n =* 3) mice were used for library preparation. Genomic DNA for WGBS was extracted from sorted cells using the QIAamp DNA Blood Mini Kit (Qiagen). The library for WGBS was prepared using the postbisulfite adaptor tagging (PBAT) protocol as described previously^80,81^. After library preparation, library yields were quantified using the library quantification kit (Takara Bio Inc.) and amplified using PrimeStar Max DNA polymerase (Takara Bio Inc.). The amplified libraries were sequenced on the Illumina HiSeq X platform at Macrogen.

WGBS data were aligned to the *Mus musculus* reference genome (GRCm39) using the Bismark pipeline^82^. Differentially methylated region (DMR) analysis was performed using *methylKit* package (v.1.30.0). For the genome-wide analysis, DMRs of *Tet2KO* PD-1^+^CD8^+^ TILs in comparison to *Tet2^fl/fl^* PD-1^+^CD8^+^ TILs were detected by scanning non-overlapping 1000bp-windows across the whole genome locus using the calculateDiffMeth and getMethylDiff functions, applying a q-value < 0.01 and |meth.diff| ≥ 20%. For DMR analysis focusing *Tox* gene, DMRs were detected by screening 500bp-windows at 250bp-step, applying the same statistical thresholds as in the global analysis. Differentially methylated cytosines (DMCs) within the *Tox* locus were identified at single-CpG resolution with the same q-value and methylation difference cutoffs. DMRs were annotated using the annotatePeakInBatch function from the ChIPpeakAnno package (v3.38), with mouse gene annotations from the Gencode vM35 primary assembly and the ENCODE Registry of cCREs (v3, 2021) serving as reference. DMRs overlapping enhancer, promoter, CTCF, or TSS regions were classified as regulatory DMRs (rDMRs).

To quantify the combined regulatory impact of differential methylation on gene expression, we developed a composite methylation impact score that accounts for both the breadth (number of affected regulatory elements) and depth (magnitude of methylation changes) of regulatory disruption. The impact score for each gene was calculated as:

Impact Score = (Total regulatory DMRs) × (Mean absolute methylation difference)

Where:

#### Total regulatory DMRs: the number of differentially methylated regions (qvalue < 0.01, |methylation difference| ≥ 20%) overlapping with any regulatory element (promoter, TSS, enhancer, or CTCF binding site) associated with the gene

Mean absolute methylation difference: the average magnitude of methylation change (%) across all regulatory DMRs for that gene.

This scoring approach identifies genes with extensive regulatory methylation changes that are likely to have substantial effects on transcriptional control. Genes with higher scores have both multiple affected regulatory elements and large-magnitude methylation changes.

#### *In vivo* anti-PD-1 blockade

Anti-PD-1 antibody treatment was started seven days after the intrasplenic injection of Venus-*AKTP^M/LOH^* cells. InVivoMAb anti-mouse PD-1 (RMP1-14, Bio X Cell, #BE0146) was diluted with PBS to a final concentration of 1 μg/μL and intraperitoneally injected at a dose of 10 μg/g of mouse body weight every 3 days for 6 doses in total. Control mice received InVivoMAb rat IgG2a isotype control (2A3, Bio X Cell, #815022F1) following the same dilution and dosage as the anti-PD-1 antibody treatment protocol. Mice were sacrificed 30 days after the injection of Venus-*AKTP^M/LOH^* cells to collect the whole liver for LMTB evaluation.

#### Competitive bone marrow transplantation (BMT) mouse model with CRC liver metastasis

Bone marrow isolation and preparation: Bone marrow cells were harvested from femurs and tibias of 8-10 week-old donor mice. Bones were flushed with ice-cold PBS containing 2% FBS using a 23-gauge needle. Cell suspensions were filtered through 70-μm cell strainers. Viable cell counts were determined by trypan blue exclusion.

Lethal irradiation and competitive transplantation: Recipient CD45.1⁺ C57BL/6J mice (8-12 weeks old) were lethally irradiated using an X-ray irradiator (Hitachi Power Solutions MBR-1520R-3) with a split-dose protocol of 4.5 Gy twice, administered 4 hours apart (total dose of 9.0 Gy) while housed in pie cages to ensure uniform radiation exposure. For *Tet2KO:Tet2WT* chimeras, *Vav1Cre⁺Tet2^fl/fl^* CD45.2⁺ bone marrow cells were mixed with *Tet2WT* CD45.1⁺/CD45.2⁺ competitor bone marrow cells at ratios of 1:2 or 1:3. Control *Tet2^fl/fl^:Tet2WT* chimeras received a 1:1 mixture of *Tet2ᶠˡ/ᶠˡ* CD45.2⁺ and *Tet2WT* CD45.1⁺/CD45.2⁺ bone marrow cells. A total of 2×10⁶ bone marrow cells in 200 μL PBS were injected via retro-orbital venous plexus within 4-6 hours post-irradiation. Mice were maintained in sterilized caging, food, and water were provided during the first 14 days after transplantation.

Chimerism was assessed in peripheral blood collected via facial vein bleeding at 4 weeks and re-evaluated at 12 weeks post-BMT to confirm the proliferative advantage of *Tet2KO* hematopoietic and lymphoid compartments. Donor cell engraftment was quantified by flow cytometry analysis of CD45.1 and CD45.2 expression.

After confirming adequate hematopoietic reconstitution at four weeks post-BMT, Venus-*AKTP^M/LOH^* cells were injected intrasplenically to establish liver metastases. Eight weeks after cancer cell injection, mice were euthanized and livers were harvested and processed for histological analysis. LMTB was quantified by microscopic measurement as described above. Tumor-infiltrating immune cells were isolated from liver metastases by enzymatic digestion as described above and subsequently analyzed by flow cytometry to identify exhausted T-cell populations using a panel of fluorophore-conjugated anti-mouse antibodies, consisting of anti-CD3ε, anti-CD4, anti-CD8α, anti-PD-1, anti-Tim3 (Havcr2/CD366), anti-CD45.1, and anti-CD45.2 antibodies (see Table S27B).

#### *In vitro* modeling of CD8 T-cell exhaustion

CD8 T cells were isolated from spleens of *CD4^Cre^Tet2KO* mice, and *Tet2^fl/fl^* mice using the Mouse CD8 T Cell Isolation Kit (Miltenyi Biotec, # 130-095-236) according to the manufacturer’s instructions. Purity (>90% CD45^+^CD3^+^CD8^+^ cells) was confirmed by flow cytometry.

Purified CD8 T cells were seeded at 2×10^7^ cells per well in 6-well plates and activated with plate-bound anti-CD3ε antibody (2 µg/mL, clone 145-2C11, BioLegend, #100340) and soluble anti-CD28 antibody (2 µg/mL, clone 37.51, BioLegend, #102116) in complete RPMI 1640 medium supplemented with 10% FBS, 1% penicillin-streptomycin, 50 µM β-mercaptoethanol, 10 mM HEPES, and recombinant mouse IL-2 (125 U/mL, PeproTech, #212-12). After 4 days, T cells were harvested and transferred to fresh wells pre-coated with anti-CD3 antibody alone (2 µg/mL) for chronic re-stimulation. This re-stimulation was repeated on day 8. T cells were cultured for a total of 12-14 days at 37°C with 5% CO_2_, with medium containing IL-2 (125 U/mL) refreshed every 2-3 days.

At indicated timepoints (days 7 and 12), T cells were harvested and assessed for exhaustion markers by flow cytometry using panels of fluorophore-conjugated anti-mouse antibodies, consisting of anti-CD3ε, anti-CD8α, anti-PD-1, anti-Tim3 (Havcr2/CD366), and anti-Tcf1 antibodies (see Table S27C). For intracellular staining, cells were first stained with Fixable Viability Dye eFluor 780 (eBioscience, #65-0865-14), followed by cell-surface markers. Cells were then fixed and permeabilized using the Foxp3/Transcription Factor Staining Buffer Set (eBioscience, #00-5523-00) according to the manufacturer’s instructions, and stained with intracellular antibodies for 30 minutes at room temperature. Data were acquired on BD FACSAria III and analyzed using FlowJo software (BD Biosciences).

## Acknowledgements

We thank the staff of the associated institutions and departments for their technical assistance throughout this project. We are grateful to the staff of the University of Tsukuba Animal Resource Center for their support with animal care. The authors also thank Editage (www.editage.com) for language editing. Figures were created using BioRender.com.

This work was supported by Grants-in-Aid for Scientific Research (KAKENHI: JP25K19551 [M.F.], JP25K19567 [S.S.], JP25K19550 [Y.S.], JP23K15293 [K.M.], and JP25H01055 [M.S.-Y.]) from the Ministry of Education, Culture, Sports, and Science of Japan; AMED (JP256f0137008 [M.F.], JP23tk0124002, JP24ck0106908, JP23ck0106797, JP22H04925 (PAGS), JP22zf0127009 (the Moonshot Research and Development Program) and JP25gm2010006 (CREST) [M.S.-Y.]); Kobayashi Foundation for Cancer Research, Japan Leukemia Research Fund, and Takeda Science [S.S.], and The Uehara Memorial Foundation, Takeda Science, Kobayashi Foundation for Cancer Research, Kobayashi Foundation, and the Chemo-Sero Therapeutic Research Institute [M.S.-Y.]. M.F. was supported by Gilead’s Research Scholars Program and a grant from The Leukemia & Lymphoma Society (Grant ID: 3442-25). G.C. was supported by the National Sciences and Engineering Research Council of Canada (Award 618788). R.J.V. reports funding from Canadian Institutes for Health Research (PJT 203869, PJF 204160), The Princess Margaret Cancer Foundation, The Leukemia and Lymphoma Society of Canada (1445410), the Brain Tumour Foundation of Canada, Lymphoma Canada, Marathon of Hope Cancer Centres Network Clinician Scientist Award (3256-04), Brain Canada and Cancer Research Society Translational Research Grant in Brain Cancer, Cancer Research Society (1455586), and a University of Toronto Hold ‘Em For Life Early Career Professorship. The funders had no role in the study design, data collection and analysis, or decision to publish or prepare the manuscript.

## Author contributions

**Conceptualization**: M.S.-Y and R.J.V.. **Data Curation**: A.N.T.L., M.F., G.C., R.J.V., T.B.N., S.S., Y.S., F.M., H.A., Y.T.M.N. **Formal analysis**: A.N.T.L., R.J.V., T.B.N., and M.F. **Funding acquisition**: M.F., Y.S., K.M., R.J.V., and M.S.-Y. **Investigation**: A.N.T.L., T.B.N., Y.T.M.N., G.C. **Methodology**: A.N.T.L., T.B.N., M.F., R.J.V., F.M., H.A., H.O., M.N., M.K., M.O., and M.S.-Y. **Project administration**: A.N.T.L., M.F., R.J.V., and M.S.- Y. **Resources**: A.N.T.L., M.F., T.B.N., G.C., Y.T.M.N., F.M., H.A., S.S., K.M., T.S., H.O., M.N., M.O., S.I., S.S., Y.S., M.K., R.J.V., and M.S.-Y. **Software**: A.N.T.L., R.J.V., M.F., and T.B.N. **Supervision**: M.F., R.J.V., and M.S.-Y. **Validation**: A.N.T.L., R.J.V., and T.B.N. **Visualization**: A.N.T.L., R.J.V., and M.F. **Writing-Original Draft**: A.N.T.L., R.J.V., M.F., and M.S.-Y. **Writing-Review & Editing**: A.N.T.L., M.F., R.J.V., G.C., and M.S.-Y.

## Declaration of generative AI and AI-assisted technologies

During manuscript preparation, generative AI tools were used solely for language editing and clarity improvement. The authors reviewed and validated all content and take full responsibility for the manuscript.

## RESOURCE AVAILABILITY

### Lead contact

Further information and requests for resources and reagents should be directed to and will be fulfilled by the lead contact, Mamiko Sakata-Yanagimoto.

### Materials availability

This study did not generate new unique reagents.

### Data and code availability

Statistical source data and detailed results for human cohort analyses (Figure 1 and Figure S1) are provided in Table S1 and Table S2. All original code will be deposited at GitHub and made publicly available as of the date of publication. The code for scRNA-seq and WGBS analysis will be available at: [Link will be provided upon publication]. The code for clonal hematopoiesis correlation analysis is available at: https://github.com/GiacomoCampo/msk_ch_correlations. Any additional information required to reanalyze the data reported in this paper is available from the lead contact upon request.

## REFERENCES

1. Luyendijk, M., Visser, O., Blommestein, H.M., de Hingh, I., Hoebers, F.J.P., Jager, A., Sonke, G.S., de Vries, E.G.E., Uyl-de Groot, C.A., and Siesling, S. (2023). Changes in survival in de novo metastatic cancer in an era of new medicines. J Natl Cancer Inst 115, 628–635. 10.1093/jnci/djad020.

2. Kitamura, T., Qian, B.Z., and Pollard, J.W. (2015). Immune cell promotion of metastasis. Nat Rev Immunol 15, 73–86. 10.1038/nri3789.

3. Edwards, S.C., Hoevenaar, W.H.M., and Coffelt, S.B. (2021). Emerging immunotherapies for metastasis. British Journal of Cancer 124, 37–48. 10.1038/s41416-020-01160-5.

4. Arias-Badia, M., Chang, R., and Fong, L. (2024). γδ T cells as critical anti-tumor immune effectors. Nature Cancer 5, 1145–1157. 10.1038/s43018-024-00798-x.

5. Dean, I., Lee, C.Y.C., Tuong, Z.K., Li, Z., Tibbitt, C.A., Willis, C., Gaspal, F., Kennedy, B.C., Matei-Rascu, V., Fiancette, R., et al. (2024). Rapid functional impairment of natural killer cells following tumor entry limits anti-tumor immunity. Nature Communications 15, 683. 10.1038/s41467-024-44789-z.

6. Portale, F., and Di Mitri, D. (2023). NK Cells in Cancer: Mechanisms of Dysfunction and Therapeutic Potential. Int J Mol Sci 24. 10.3390/ijms24119521.

7. Xie, F., Zhou, X., Su, P., Li, H., Tu, Y., Du, J., Pan, C., Wei, X., Zheng, M., Jin, K., et al. (2022). Breast cancer cell-derived extracellular vesicles promote CD8+ T cell exhaustion via TGF-β type II receptor signaling. Nature Communications 13, 4461. 10.1038/s41467-022-31250-2.

8. Baessler, A., and Vignali, D.A.A. (2024). T Cell Exhaustion. Annu Rev Immunol 42, 179–206. 10.1146/annurev-immunol-090222-110914.

9. Gavil, N.V., Scott, M.C., Weyu, E., Smith, O.C., O’Flanagan, S.D., Wijeyesinghe, S., Lotfi-Emran, S., Shiao, S.L., Vezys, V., and Masopust, D. (2023). Chronic antigen in solid tumors drives a distinct program of T cell residence. Sci Immunol 8, eadd5976. 10.1126/sciimmunol.add5976.

10. Coffelt, S.B., Kersten, K., Doornebal, C.W., Weiden, J., Vrijland, K., Hau, C.-S., Verstegen, N.J.M., Ciampricotti, M., Hawinkels, L.J.A.C., Jonkers, J., and de Visser, K.E. (2015). IL-17-producing γδ T cells and neutrophils conspire to promote breast cancer metastasis. Nature 522, 345–348. 10.1038/nature14282.

11. Wu, P., Wu, D., Ni, C., Ye, J., Chen, W., Hu, G., Wang, Z., Wang, C., Zhang, Z., Xia, W., et al. (2014). γδT17 cells promote the accumulation and expansion of myeloid-derived suppressor cells in human colorectal cancer. Immunity 40, 785–800. 10.1016/j.immuni.2014.03.013.

12. Hedrick, C.C., and Malanchi, I. (2022). Neutrophils in cancer: heterogeneous and multifaceted. Nature Reviews Immunology 22, 173–187. 10.1038/s41577-021-00571-6.

13. de Visser, K.E., and Joyce, J.A. (2023). The evolving tumor microenvironment: From cancer initiation to metastatic outgrowth. Cancer Cell 41, 374–403. 10.1016/j.ccell.2023.02.016.

14. Xu, J., Ding, L., Mei, J., Hu, Y., Kong, X., Dai, S., Bu, T., Xiao, Q., and Ding, K. (2025). Dual roles and therapeutic targeting of tumor-associated macrophages in tumor microenvironments. Signal Transduction and Targeted Therapy 10, 268. 10.1038/s41392-025-02325-5.

15. Herbrich, S., Chaib, M., Anandhan, S., Andrewes, S.W., Nagarajan, A., Guan, B., Gandhi, N., Gilliam, J., Radovich, M., and Sharma, P. (2026). *TET2* mutant clonal hematopoiesis enhances macrophage antigen presentation and improves immune checkpoint therapy in solid tumors. Cancer Cell 44, 187–202.e187. 10.1016/j.ccell.2025.09.011.

16. Carnevale, S., Di Ceglie, I., Grieco, G., Rigatelli, A., Bonavita, E., and Jaillon, S. (2023). Neutrophil diversity in inflammation and cancer. Front Immunol 14, 1180810. 10.3389/fimmu.2023.1180810.

17. Jaillon, S., Ponzetta, A., Di Mitri, D., Santoni, A., Bonecchi, R., and Mantovani, A. (2020). Neutrophil diversity and plasticity in tumour progression and therapy. Nat Rev Cancer 20, 485–503. 10.1038/s41568-020-0281-y.

18. Sebestyen, Z., Prinz, I., Déchanet-Merville, J., Silva-Santos, B., and Kuball, J. (2020). Translating gammadelta (γδ) T cells and their receptors into cancer cell therapies. Nature Reviews Drug Discovery 19, 169–184. 10.1038/s41573-019-0038-z.

19. Bruni, E., Cimino, M.M., Donadon, M., Carriero, R., Terzoli, S., Piazza, R., Ravens, S., Prinz, I., Cazzetta, V., Marzano, P., et al. (2022). Intrahepatic CD69(+)Vδ1 T cells re-circulate in the blood of patients with metastatic colorectal cancer and limit tumor progression. J Immunother Cancer 10. 10.1136/jitc-2022-004579.

20. Wu, Y., Kyle-Cezar, F., Woolf, R.T., Naceur-Lombardelli, C., Owen, J., Biswas, D., Lorenc, A., Vantourout, P., Gazinska, P., Grigoriadis, A., et al. (2019). An innate-like Vδ1(+) γδ T cell compartment in the human breast is associated with remission in triple-negative breast cancer. Sci Transl Med 11. 10.1126/scitranslmed.aax9364.

21. Jaiswal, S., and Ebert, B.L. (2019). Clonal hematopoiesis in human aging and disease. Science 366, eaan4673. doi:10.1126/science.aan4673.

22. Bolton, K.L., Ptashkin, R.N., Gao, T., Braunstein, L., Devlin, S.M., Kelly, D., Patel, M., Berthon, A., Syed, A., Yabe, M., et al. (2020). Cancer therapy shapes the fitness landscape of clonal hematopoiesis. Nature Genetics 52, 1219–1226. 10.1038/s41588-020-00710-0.

23. Coombs, C.C., Zehir, A., Devlin, S.M., Kishtagari, A., Syed, A., Jonsson, P., Hyman, D.M., Solit, D.B., Robson, M.E., and Baselga, J. (2017). Therapy-related clonal hematopoiesis in patients with non-hematologic cancers is common and associated with adverse clinical outcomes. Cell stem cell 21, 374–382. e374.

24. Kleppe, M., Comen, E., Wen, H.Y., Bastian, L., Blum, B., Rapaport, F.T., Keller, M., Granot, Z., Socci, N., Viale, A., et al. (2015). Somatic mutations in leukocytes infiltrating primary breast cancers. npj Breast Cancer 1, 15005. 10.1038/npjbcancer.2015.5.

25. Severson, E.A., Riedlinger, G.M., Connelly, C.F., Vergilio, J.A., Goldfinger, M., Ramkissoon, S., Frampton, G.M., Ross, J.S., Fratella-Calabrese, A., Gay, L., et al. (2018). Detection of clonal hematopoiesis of indeterminate potential in clinical sequencing of solid tumor specimens. Blood 131, 2501–2505. 10.1182/blood-2018-03-840629.

26. Marshall, C.H., Gondek, L.P., Luo, J., and Antonarakis, E.S. (2022). Clonal Hematopoiesis of Indeterminate Potential in Patients with Solid Tumor Malignancies. Cancer Res 82, 4107–4113. 10.1158/0008-5472.Can-22-0985.

27. Pich, O., Bernard, E., Zagorulya, M., Rowan, A., Pospori, C., Slama, R., Huerga Encabo, H., O’Sullivan, J., Papazoglou, D., Anastasiou, P., et al. (2025). Tumor-Infiltrating Clonal Hematopoiesis. New England Journal of Medicine 392, 1594–1608. 10.1056/NEJMoa2413361.

28. Buttigieg, M.M., Vlasschaert, C., Bick, A.G., Vanner, R.J., and Rauh, M.J. (2025). Inflammatory reprogramming of the solid tumor microenvironment by infiltrating clonal hematopoiesis is associated with adverse outcomes. Cell Rep Med 6, 101989. 10.1016/j.xcrm.2025.101989.

29. Pan, W., Zhu, S., Qu, K., Meeth, K., Cheng, J., He, K., Ma, H., Liao, Y., Wen, X., Roden, C., et al. (2017). The DNA Methylcytosine Dioxygenase Tet2 Sustains Immunosuppressive Function of Tumor-Infiltrating Myeloid Cells to Promote Melanoma Progression. Immunity 47, 284–297.e285. 10.1016/j.immuni.2017.07.020.

30. Nguyen, Y.T.M., Fujisawa, M., Nguyen, T.B., Suehara, Y., Sakamoto, T., Matsuoka, R., Abe, Y., Fukumoto, K., Hattori, K., Noguchi, M., et al. (2021). Tet2 deficiency in immune cells exacerbates tumor progression by increasing angiogenesis in a lung cancer model. Cancer Sci 112, 4931–4943. 10.1111/cas.15165.

31. Krishnan, T., Solar Vasconcelos, J.P., Titmuss, E., Topham, J.T., Schaeffer, D.F., Karsan, A., Lim, H.J., Ho, C., Gill, S., Kennecke, H.F., et al. (2024). Clonal hematopoiesis of indeterminate potential (CHIP), treatment outcomes and adverse events in gastrointestinal cancers: A pooled analysis of clinical trial and real-world data. Journal of Clinical Oncology 42, 169–169. 10.1200/JCO.2024.42.3_suppl.169.

32. Reed, S.C., Croessmann, S., and Park, B.H. (2023). CHIP Happens: Clonal Hematopoiesis of Indeterminate Potential and Its Relationship to Solid Tumors. Clin Cancer Res 29, 1403–1411. 10.1158/1078-0432.Ccr-22-2598.

33. Rondeau, V., Bansal, S., Buttigieg, M.M., Zeng, A.G.X., Chan, D.Y., Chan-Seng-Yue, M., Jin, L., McLeod, J., Kates, M., Donato, E., et al. (2026). Response to Immune Checkpoint Blockade Is Enhanced in the Presence of Hematopoietic TET2 Inactivation. Cancer Research, OF845-OF857. 10.1158/0008-5472.CAN-24-3329.

34. Nguyen, B., Fong, C., Luthra, A., Smith, S.A., DiNatale, R.G., Nandakumar, S., Walch, H., Chatila, W.K., Madupuri, R., Kundra, R., et al. (2022). Genomic characterization of metastatic patterns from prospective clinical sequencing of 25,000 patients. Cell 185, 563–575.e511. 10.1016/j.cell.2022.01.003.

35. Quivoron, C., Couronne, L., Della Valle, V., Lopez, C.K., Plo, I., Wagner-Ballon, O., Do Cruzeiro, M., Delhommeau, F., Arnulf, B., Stern, M.H., et al. (2011). TET2 inactivation results in pleiotropic hematopoietic abnormalities in mouse and is a recurrent event during human lymphomagenesis. Cancer Cell 20, 25–38. S1535-6108(11)00225-X [pii] 10.1016/j.ccr.2011.06.003.

36. Stadtfeld, M., and Graf, T. (2005). Assessing the role of hematopoietic plasticity for endothelial and hepatocyte development by non-invasive lineage tracing. Development 132, 203–213. 10.1242/dev.01558.

37. Lee, P.P., Fitzpatrick, D.R., Beard, C., Jessup, H.K., Lehar, S., Makar, K.W., Pérez-Melgosa, M., Sweetser, M.T., Schlissel, M.S., Nguyen, S., et al. (2001). A critical role for Dnmt1 and DNA methylation in T cell development, function, and survival. Immunity 15, 763–774. 10.1016/s1074-7613(01)00227-8.

38. Rickert, R.C., Roes, J., and Rajewsky, K. (1997). B lymphocyte-specific, Cre-mediated mutagenesis in mice. Nucleic Acids Res 25, 1317–1318. 10.1093/nar/25.6.1317.

39. Clausen, B.E., Burkhardt, C., Reith, W., Renkawitz, R., and Forster, I. (1999). Conditional gene targeting in macrophages and granulocytes using LysMcre mice. Transgenic Res 8, 265–277. 10.1023/a:1008942828960.

40. Nakayama, M., Hong, C.P., Oshima, H., Sakai, E., Kim, S.-J., and Oshima, M. (2020). Loss of wild-type p53 promotes mutant p53-driven metastasis through acquisition of survival and tumor-initiating properties. Nature Communications 11, 2333. 10.1038/s41467-020-16245-1.

41. Huang, D.W., Sherman, B.T., Tan, Q., Collins, J.R., Alvord, W.G., Roayaei, J., Stephens, R., Baseler, M.W., Lane, H.C., and Lempicki, R.A. (2007). The DAVID Gene Functional Classification Tool: a novel biological module-centric algorithm to functionally analyze large gene lists. Genome Biology 8, R183. 10.1186/gb-2007-8-9-r183.

42. Subramanian, A., Tamayo, P., Mootha, V.K., Mukherjee, S., Ebert, B.L., Gillette, M.A., Paulovich, A., Pomeroy, S.L., Golub, T.R., Lander, E.S., and Mesirov, J.P. (2005). Gene set enrichment analysis: a knowledge-based approach for interpreting genome-wide expression profiles. Proc Natl Acad Sci U S A 102, 15545–15550. 0506580102 [pii] 10.1073/pnas.0506580102.

43. Godec, J., Tan, Y., Liberzon, A., Tamayo, P., Bhattacharya, S., Butte, A.J., Mesirov, J.P., and Haining, W.N. (2016). Compendium of Immune Signatures Identifies Conserved and Species-Specific Biology in Response to Inflammation. Immunity 44, 194–206. 10.1016/j.immuni.2015.12.006.

44. Aibar, S., González-Blas, C.B., Moerman, T., Huynh-Thu, V.A., Imrichova, H., Hulselmans, G., Rambow, F., Marine, J.-C., Geurts, P., Aerts, J., et al. (2017). SCENIC: single-cell regulatory network inference and clustering. Nature Methods 14, 1083–1086. 10.1038/nmeth.4463.

45. Zhang, C., Sheng, Q., Zhang, X., Xu, K., Jin, X., Zhou, W., Zhang, M., Lv, D., Yang, C., Li, Y., et al. (2023). Prioritizing exhausted T cell marker genes highlights immune subtypes in pan-cancer. iScience 26, 106484. 10.1016/j.isci.2023.106484.

46. Trapnell, C., Cacchiarelli, D., Grimsby, J., Pokharel, P., Li, S., Morse, M., Lennon, N.J., Livak, K.J., Mikkelsen, T.S., and Rinn, J.L. (2014). The dynamics and regulators of cell fate decisions are revealed by pseudotemporal ordering of single cells. Nature Biotechnology 32, 381–386. 10.1038/nbt.2859.

47. Borcherding, N., Bormann, N.L., and Kraus, G. (2020). scRepertoire: An R-based toolkit for single-cell immune receptor analysis. F1000Res 9, 47. 10.12688/f1000research.22139.2.

48. Bosco, N., Swee, L.K., Bénard, A., Ceredig, R., and Rolink, A. (2010). Auto-reconstitution of the T-cell compartment by radioresistant hematopoietic cells following lethal irradiation and bone marrow transplantation. Experimental Hematology 38, 222–232.e222. 10.1016/j.exphem.2009.12.006.

49. Busque, L., Patel, J.P., Figueroa, M.E., Vasanthakumar, A., Provost, S., Hamilou, Z., Mollica, L., Li, J., Viale, A., Heguy, A., et al. (2012). Recurrent somatic TET2 mutations in normal elderly individuals with clonal hematopoiesis. Nature Genetics 44, 1179–1181. 10.1038/ng.2413.

50. Zehir, A., Benayed, R., Shah, R.H., Syed, A., Middha, S., Kim, H.R., Srinivasan, P., Gao, J., Chakravarty, D., Devlin, S.M., et al. (2017). Mutational landscape of metastatic cancer revealed from prospective clinical sequencing of 10,000 patients. Nature Medicine 23, 703–713. 10.1038/nm.4333.

51. Jaiswal, S. (2020). Clonal hematopoiesis and nonhematologic disorders. Blood 136, 1606–1614. 10.1182/blood.2019000989.

52. Fraietta, J.A., Nobles, C.L., Sammons, M.A., Lundh, S., Carty, S.A., Reich, T.J., Cogdill, A.P., Morrissette, J.J.D., DeNizio, J.E., Reddy, S., et al. (2018). Disruption of TET2 promotes the therapeutic efficacy of CD19-targeted T cells. Nature 558, 307–312. 10.1038/s41586-018-0178-z.

53. Dimitri, A.J., Baxter, A.E., Chen, G.M., Hopkins, C.R., Rouin, G.T., Huang, H., Kong, W., Holliday, C.H., Wiebking, V., Bartoszek, R., et al. (2024). TET2 regulates early and late transitions in exhausted CD8(+) T cell differentiation and limits CAR T cell function. Sci Adv 10, eadp9371. 10.1126/sciadv.adp9371.

54. Wang, Y., Sano, S., Yura, Y., Ke, Z., Sano, M., Oshima, K., Ogawa, H., Horitani, K., Min, K.D., Miura-Yura, E., et al. (2020). Tet2-mediated clonal hematopoiesis in nonconditioned mice accelerates age-associated cardiac dysfunction. JCI Insight 5. 10.1172/jci.insight.135204.

55. Fuster, J.J., MacLauchlan, S., Zuriaga, M.A., Polackal, M.N., Ostriker, A.C., Chakraborty, R., Wu, C.L., Sano, S., Muralidharan, S., Rius, C., et al. (2017). Clonal hematopoiesis associated with TET2 deficiency accelerates atherosclerosis development in mice. Science 355, 842–847. 10.1126/science.aag1381.

56. Jaiswal, S., Natarajan, P., Silver Alexander, J., Gibson Christopher, J., Bick Alexander, G., Shvartz, E., McConkey, M., Gupta, N., Gabriel, S., Ardissino, D., et al. Clonal Hematopoiesis and Risk of Atherosclerotic Cardiovascular Disease. New England Journal of Medicine 377, 111–121. 10.1056/NEJMoa1701719.

57. Sano, S., Oshima, K., Wang, Y., MacLauchlan, S., Katanasaka, Y., Sano, M., Zuriaga, M.A., Yoshiyama, M., Goukassian, D., Cooper, M.A., et al. (2018). Tet2-Mediated Clonal Hematopoiesis Accelerates Heart Failure Through a Mechanism Involving the IL-1β/NLRP3 Inflammasome. J Am Coll Cardiol 71, 875–886. 10.1016/j.jacc.2017.12.037.

58. Bolton, K.L., Gillis, N.K., Coombs, C.C., Takahashi, K., Zehir, A., Bejar, R., Garcia-Manero, G., Futreal, A., Jensen, B.C., Diaz, L.A., Jr., et al. (2019). Managing Clonal Hematopoiesis in Patients With Solid Tumors. J Clin Oncol 37, 7–11. 10.1200/jco.18.00331.

59. Li, S., Feng, J., Wu, F., Cai, J., Zhang, X., Wang, H., Fetahu, I.S., Iwanicki, I., Ma, D., Hu, T., et al. (2020). TET2 promotes anti-tumor immunity by governing G-MDSCs and CD8+ T-cell numbers. EMBO reports 21, e49425. 10.15252/embr.201949425.

60. Carty, S.A., Gohil, M., Banks, L.B., Cotton, R.M., Johnson, M.E., Stelekati, E., Wells, A.D., Wherry, E.J., Koretzky, G.A., and Jordan, M.S. (2018). The Loss of TET2 Promotes CD8(+) T Cell Memory Differentiation. J Immunol 200, 82–91. 10.4049/jimmunol.1700559.

61. Yue, X., Trifari, S., Äijö, T., Tsagaratou, A., Pastor, W.A., Zepeda-Martínez, J.A., Lio, C.W., Li, X., Huang, Y., Vijayanand, P., et al. (2016). Control of Foxp3 stability through modulation of TET activity. J Exp Med 213, 377–397. 10.1084/jem.20151438.

62. Khan, O., Giles, J.R., McDonald, S., Manne, S., Ngiow, S.F., Patel, K.P., Werner, M.T., Huang, A.C., Alexander, K.A., Wu, J.E., et al. (2019). TOX transcriptionally and epigenetically programs CD8+ T cell exhaustion. Nature 571, 211–218. 10.1038/s41586-019-1325-x.

63. Kim, K., Park, S., Park, S.Y., Kim, G., Park, S.M., Cho, J.W., Kim, D.H., Park, Y.M., Koh, Y.W., Kim, H.R., et al. (2020). Single-cell transcriptome analysis reveals TOX as a promoting factor for T cell exhaustion and a predictor for anti-PD-1 responses in human cancer. Genome Med 12, 22. 10.1186/s13073-020-00722-9.

64. Zhang, X., Wang, Y., Xue, H., Zhao, Y., Liu, M., Wei, H., and Liu, Q. (2025). Clonal hematopoiesis of indeterminate potential and risk of autoimmune thyroid disease. BMC Med 23, 237. 10.1186/s12916-025-04077-z.

65. Liu, Q., Wästerlid, T., Smedby, K.E., Xue, H., Boberg, E., Fang, F., and Liu, X. (2025). Clonal hematopoiesis of indeterminate potential and risk of immune thrombocytopenia. J Intern Med 297, 672–682. 10.1111/joim.20092.

66. Jaber Chehayeb, R., Singh, J., Matute-Martinez, C., Chen, N.W., Guajardo, A.F., Lin, D., Jayakrishnan, R., Christofides, A., Leveille, E., Im, Y., et al. (2024). Clonal hematopoiesis of indeterminate potential is associated with increased risk of immune checkpoint inhibitor myocarditis in a prospective study of a cardio-oncology cohort. Cardiooncology 10, 84. 10.1186/s40959-024-00289-z.

67. Gao, Q., Shen, K., and Xiao, M. (2024). TET2 mutation in acute myeloid leukemia: biology, clinical significance, and therapeutic insights. Clin Epigenetics 16, 155. 10.1186/s13148-024-01771-2.

68. An, J., González-Avalos, E., Chawla, A., Jeong, M., López-Moyado, I.F., Li, W., Goodell, M.A., Chavez, L., Ko, M., and Rao, A. (2015). Acute loss of TET function results in aggressive myeloid cancer in mice. Nature Communications 6, 10071. 10.1038/ncomms10071.

69. Muto, H., Sakata-Yanagimoto, M., Nagae, G., Shiozawa, Y., Miyake, Y., Yoshida, K., Enami, T., Kamada, Y., Kato, T., Uchida, K., et al. (2014). Reduced TET2 function leads to T-cell lymphoma with follicular helper T-cell-like features in mice. Blood Cancer J 4, e264. 10.1038/bcj.2014.83.

70. Vanner, R., Zeng, A.G.X., Chan-Seng-Yue, M., Bansal, S., Donato, E., Jin, L., Genta, S., Sanz Garcia, E., Chan, S.M., Trumpp, A., and Dick, J.E. (2023). Somatic TET2 Mutations Prime the Immune System for Response to Immune Checkpoint Blockade. Blood 142, 2689–2689. 10.1182/blood-2023-177740.

71. Carty, S.A., Gohil, M., Koretzky, G.A., and Jordan, M.S. (2014). TET2 Regulates CD8+ T Cell Differentiation. Blood 124, 1423–1423. 10.1182/blood.V124.21.1423.1423.

72. Sakai, E., Nakayama, M., Oshima, H., Kouyama, Y., Niida, A., Fujii, S., Ochiai, A., Nakayama, K.I., Mimori, K., Suzuki, Y., et al. (2018). Combined Mutation of Apc, Kras, and Tgfbr2 Effectively Drives Metastasis of Intestinal Cancer. Cancer Research 78, 1334–1346. 10.1158/0008-5472.CAN-17-3303.

73. Fujisawa, M., Nguyen, T.B., Abe, Y., Suehara, Y., Fukumoto, K., Suma, S., Makishima, K., Kaneko, C., Nguyen, Y.T.M., Usuki, K., et al. (2022). Clonal germinal center B cells function as a niche for T-cell lymphoma. Blood 140, 1937–1950. 10.1182/blood.2022015451.

74. Krämer, A., Green, J., Pollard, J., Jr., and Tugendreich, S. (2014). Causal analysis approaches in Ingenuity Pathway Analysis. Bioinformatics 30, 523–530. 10.1093/bioinformatics/btt703.

75. Hao, Y., Stuart, T., Kowalski, M.H., Choudhary, S., Hoffman, P., Hartman, A., Srivastava, A., Molla, G., Madad, S., Fernandez-Granda, C., and Satija, R. (2024). Dictionary learning for integrative, multimodal and scalable single-cell analysis. Nature Biotechnology 42, 293–304. 10.1038/s41587-023-01767-y.

76. Germain, P.L., Lun, A., Garcia Meixide, C., Macnair, W., and Robinson, M.D. (2021). Doublet identification in single-cell sequencing data using scDblFinder. F1000Res 10, 979. 10.12688/f1000research.73600.2.

77. Venturi, V., Kedzierska, K., Turner, S.J., Doherty, P.C., and Davenport, M.P. (2007). Methods for comparing the diversity of samples of the T cell receptor repertoire. J Immunol Methods 321, 182–195. 10.1016/j.jim.2007.01.019.

78. Thomas, P.G., Handel, A., Doherty, P.C., and La Gruta, N.L. (2013). Ecological analysis of antigen-specific CTL repertoires defines the relationship between naïve and immune T-cell populations. Proceedings of the National Academy of Sciences 110, 1839–1844. 10.1073/pnas.1222149110.

79. Greiff, V., Bhat, P., Cook, S.C., Menzel, U., Kang, W., and Reddy, S.T. (2015). A bioinformatic framework for immune repertoire diversity profiling enables detection of immunological status. Genome Medicine 7, 49. 10.1186/s13073-015-0169-8.

80. Miura, F., Shibata, Y., Miura, M., Sangatsuda, Y., Hisano, O., Araki, H., and Ito, T. (2019). Highly efficient single-stranded DNA ligation technique improves low-input whole-genome bisulfite sequencing by post-bisulfite adaptor tagging. Nucleic Acids Research 47, e85–e85. 10.1093/nar/gkz435.

81. Miura, F., Shibata, Y., Miura, M., Inatomi, K., Suzuki, Y., and Ito, T. (2025). Identification of an enzyme with strong single-stranded DNA ligation activity and its application for sequencing. Nucleic Acids Research 53, gkaf054. 10.1093/nar/gkaf054.

82. Krueger, F., and Andrews, S.R. (2011). Bismark: a flexible aligner and methylation caller for Bisulfite-Seq applications. Bioinformatics 27, 1571–1572. 10.1093/bioinformatics/btr167.

