## Supplementary material for "*TET2*-mutant clonal hematopoiesis prevents T-cell exhaustion and suppresses cancer metastasis": Figures S1-S10

### Figure S1

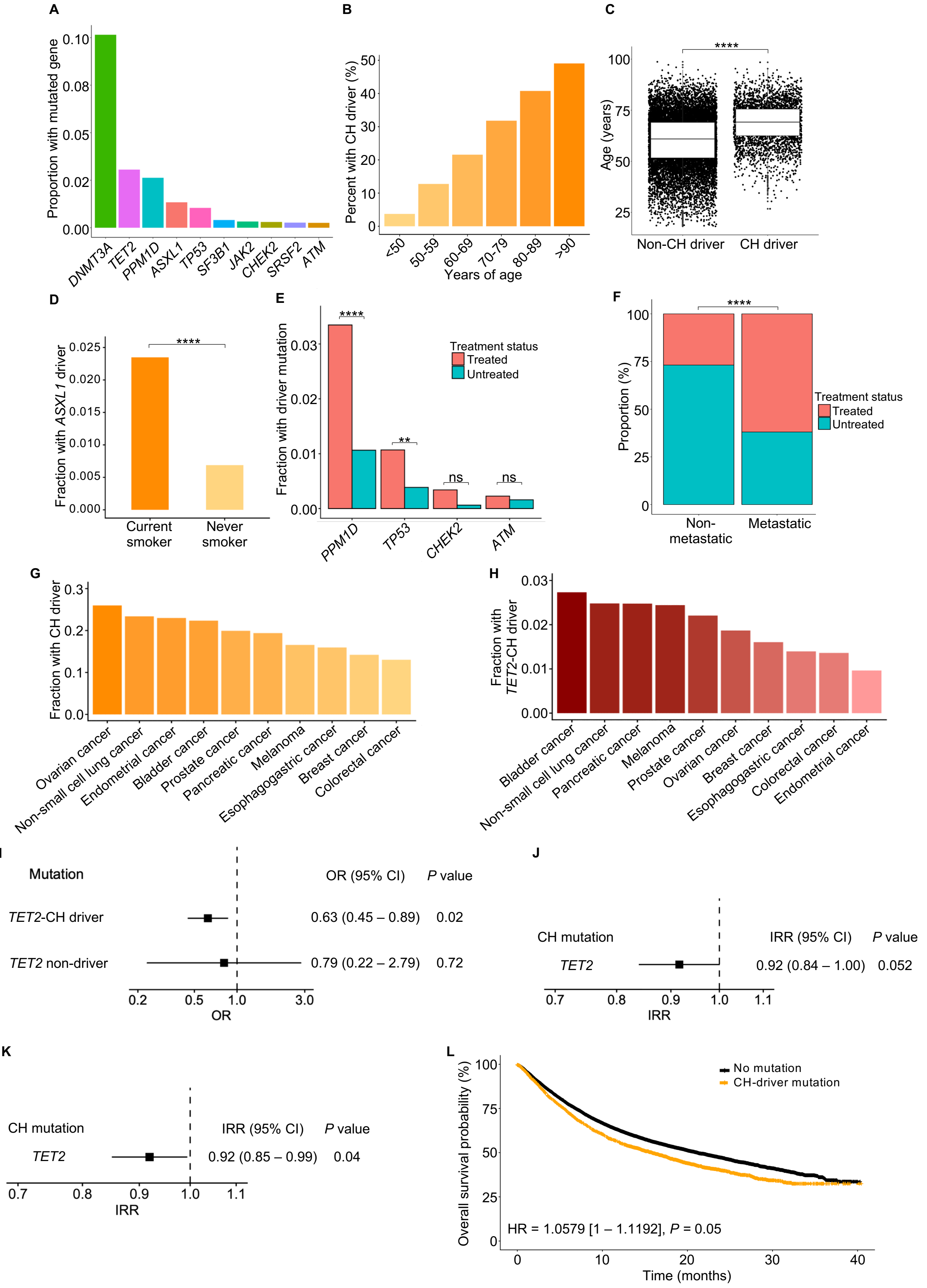
A
Proportion with mutated gene
0.10
0.08
0.05
0.02
0.00
DNMT3A
TET2
PPM1D
ASXL1
TP53
SF3B1
JAK2
CHEK2
SRSF2
ATM
B
Percent with CH driver (%)
50
40
30
20
10
0
<50
50-59
60-69
70-79
80-89
>90
Years of age
C
****
Age (years)
100
75
50
25
Non-CH driver
CH driver
D
****
Fraction with ASXL1 driver
0.025
0.020
0.015
0.010
0.005
0.000
Current smoker
Never smoker
E
****
**
ns
ns
Fraction with driver mutation
0.03
0.02
0.01
0.00
Treatment status
Treated
Untreated
PPM1D
TP53
CHEK2
ATM
F
****
Proportion (%)
100
75
50
25
0
Treatment status
Treated
Untreated
Non-metastatic
Metastatic
G
Fraction with CH driver
0.3
0.2
0.1
0.0
Ovarian cancer
Non-small cell lung cancer
Endometrial cancer
Bladder cancer
Prostate cancer
Pancreatic cancer
Melanoma
Esophagogastric cancer
Breast cancer
Colorectal cancer
H
Fraction with TET2-CH driver
0.03
0.02
0.01
0.00
Bladder cancer
Non-small cell lung cancer
Pancreatic cancer
Melanoma
Prostate cancer
Ovarian cancer
Breast cancer
Esophagogastric cancer
Colorectal cancer
Endometrial cancer
I
Mutation
OR (95% CI)
P value
TET2-CH driver
0.63 (0.45 – 0.89)
0.02
TET2 non-driver
0.79 (0.22 – 2.79)
0.72
0.2
0.5
1.0
3.0
OR
J
CH mutation
IRR (95% CI)
P value
TET2
0.92 (0.84 – 1.00)
0.052
0.7
0.8
0.9
1.0
1.1
IRR
K
CH mutation
IRR (95% CI)
P value
TET2
0.92 (0.85 – 0.99)
0.04
0.7
0.8
0.9
1.0
1.1
IRR
L
Overall survival probability (%)
100
75
50
25
0
No mutation
CH-driver mutation
HR = 1.0579 [1 – 1.1192], P = 0.05
0
10
20
30
40
Time (months)

**Figure S1. Clinical and demographic associations with CH**

(A) Proportion of patients affected with the 10 most frequent CH-driver mutations in the MSK-IMPACT cohort.

(B) Proportions of patients, stratified by age in decades, with a CH mutation.

(C) Distribution of age in patients with and without a CH-driver mutation. Difference in distribution is quantified by a multivariately adjusted linear model (LM), with *P* value reported from the regression coefficient for CH-driver mutation status.

(D) Proportion of patients with an *ASXL1* CH-driver mutation, stratified by smoking status. Difference in proportion is quantified by the regression coefficient of a multivariately adjusted logistic GLM, from which the *P* value is reported.

(E) Proportion of patients with DNA damage response driver mutations (*PPM1D*, *TP53*, *CHEK2*, *ATM*), stratified by treatment status (treated versus untreated). Differences in proportions are quantified by regression coefficients of multivariately adjusted logistic GLMs, from which the FDR-adjusted *P* values are reported.

(F) Proportion of metastatic and non-metastatic patients who have received treatment. Difference in proportion is quantified by the regression coefficient of treatment status in a multivariately adjusted logistic GLM, from which the *P* value is reported.

(G) Proportions of patients with a CH-driver mutation within each of the 10 most prevalent cancer types in the cohort.

(H) Proportions of patients with a *TET2*-CH driver mutation within each of the 10 most prevalent cancer types in the cohort.

(I) Adjusted odds of cancer metastasis stratified by *TET2*-CH driver mutation and *TET2* non-driver mutation, with the reference of no CH. Odds ratios (OR) and 95% confidence intervals are reported from logistic GLMs, and *P* values are FDR-adjusted.

(J) Multivariately adjusted IRRs from negative binomial generalized linear models testing the difference in distribution of the number of metastases in metastatic patients stratified by *TET2*-mutant CH presence.

(K) Multivariately adjusted IRRs from negative binomial generalized linear models testing the difference in distribution of the number of metastatic sites in metastatic patients stratified by *TET2*-mutant CH presence.

(L) Kaplan-Meier (KM) survival curves for overall survival of all patients, stratified by presence or absence of a CH mutation. Adjusted hazard ratios (HR) are reported from a Cox proportional hazards model, with the reference of no CH, from which the *P* value is reported.

CH, clonal hematopoiesis; GLM, generalized linear model; FDR, false discovery rate; IRR, incidence risk ratio. $^*P < 0.05$, $^{**}P < 0.01$, $^{***}P < 0.001$, $^{****}P < 0.0001$, ns = not significant. Detailed statistical outputs, test statistics, and model parameters for panels (C)-(F) and (I)-(L) are provided in Table S2.

#### Figure S2

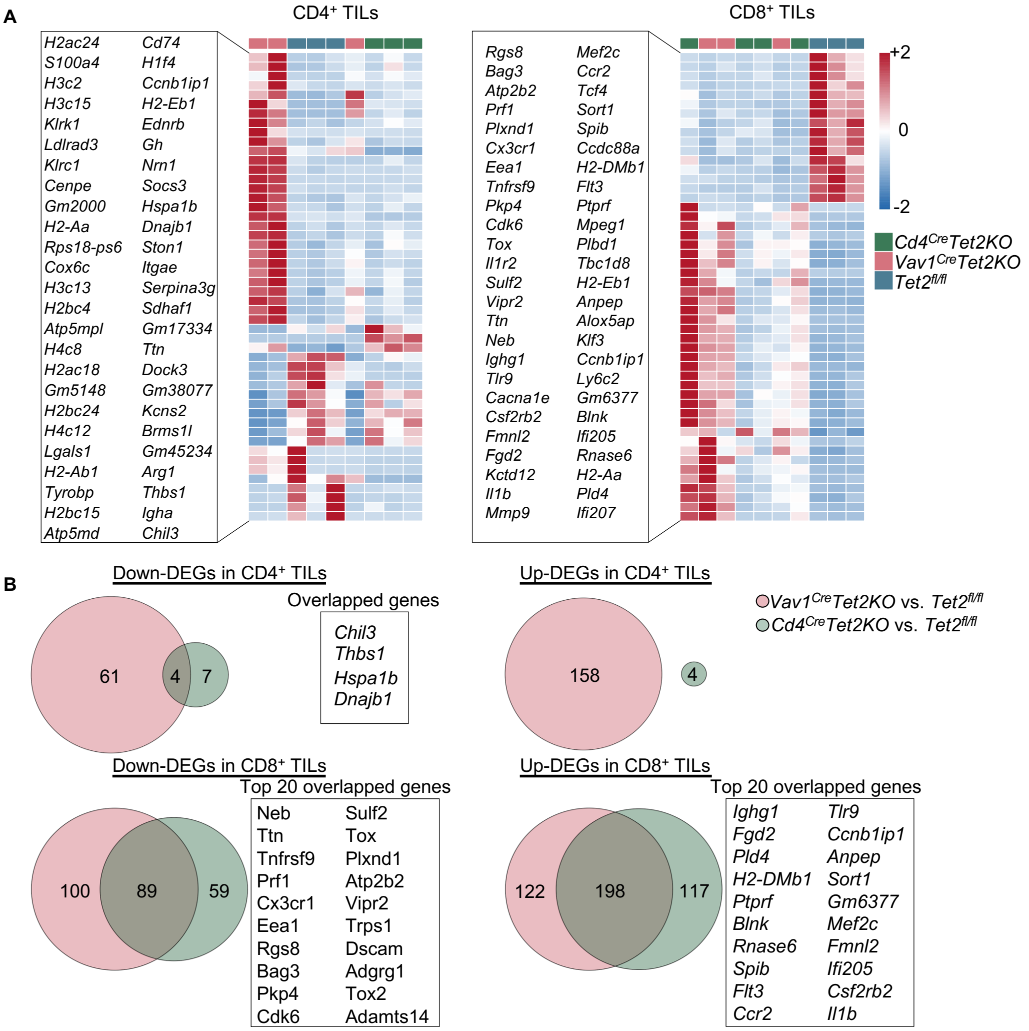


**Figure S2. Bulk RNA-sequencing analysis characterized transcriptional profiles of *Tet2*-deficient $CD4^+$ TILs and $CD8^+$ TILs.**

(A) Heatmap of top 50 DEGs ranked by combined FDR in $CD4^+$ and $CD8^+$ TILs. Color scale represents Z-score normalized expression.

(B) Venn diagrams displaying overlapped DEGs between *$Vav1^{Cre}Tet2KO$* and *$Cd4^{Cre}Tet2KO$* groups relative to *$Tet2^{fl/fl}$* group. The gene lists show top 20 overlapped DEGs ranked by combined FDR.

TILs, tumor-infiltrating lymphocytes; DEG, differentially expressed gene; Up-DEGs, upregulated DEGs; Down-DEGs, downregulated DEGs; FDR, false discovery rate. Statistical significance to identify DEGs was determined as FDR < 0.05 and |Log2 FC| > 1. Combined FDR was calculated by multiplying the two individual FDR from *$Vav1^{Cre}Tet2KO$* vs. *$Tet2^{fl/fl}$* FDR and *$Cd4^{Cre}Tet2KO$* vs. *$Tet2^{fl/fl}$* FDR comparisons.

### Figure S3

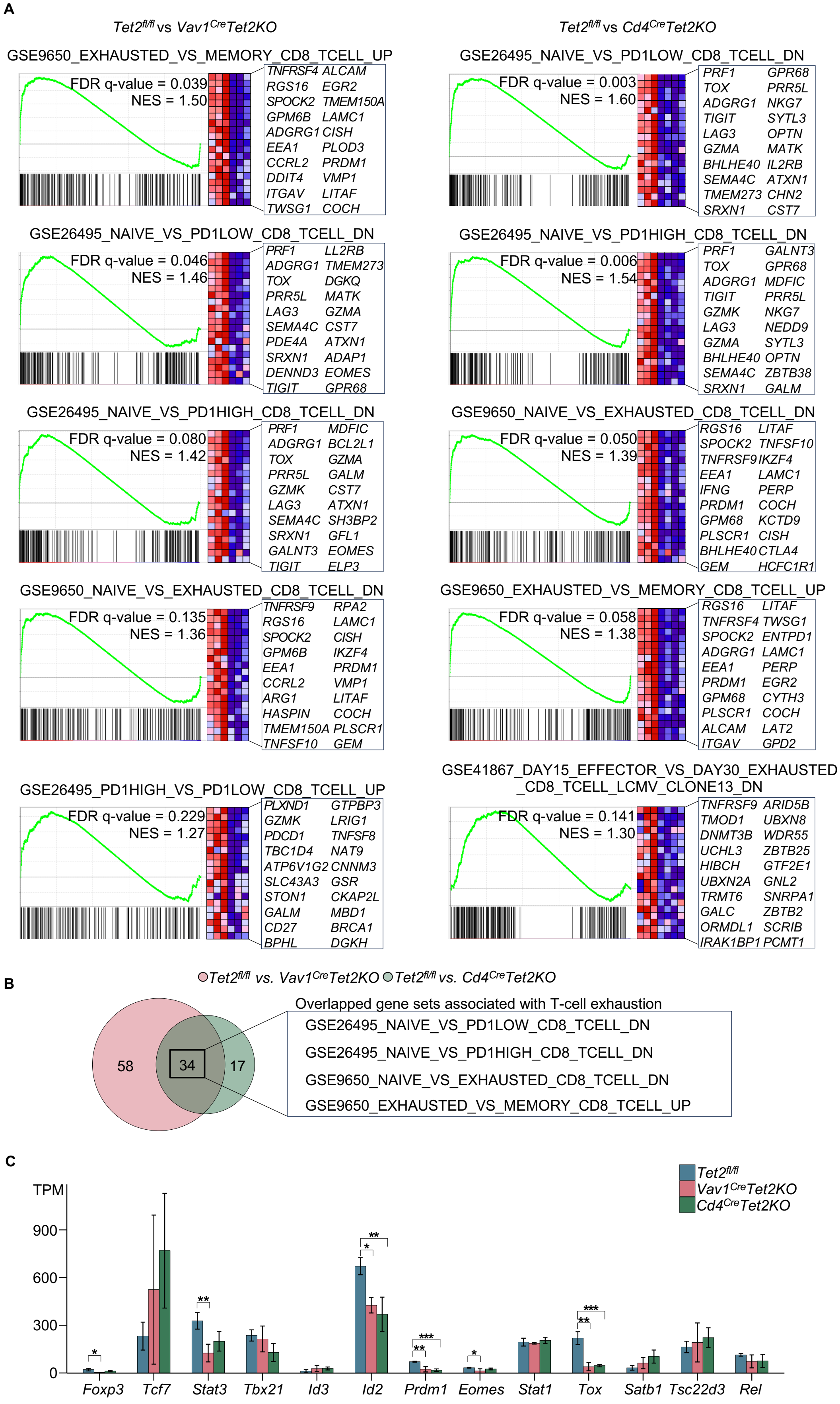

A
Tet2fl/fl vs Vav1CreTet2KO
Tet2fl/fl vs Cd4CreTet2KO
GSE9650_EXHAUSTED_VS_MEMORY_CD8_TCELL_UP
FDR q-value = 0.039
NES = 1.50
TNFRSF4 ALCAM
RGS16 EGR2
SPOCK2 TMEM150A
GPM6B LAMC1
ADGRG1 CISH
EEA1 PLOD3
CCRL2 PRDM1
DDIT4 VMP1
ITGAV LITAF
TWSG1 COCH
GSE26495_NAIVE_VS_PD1LOW_CD8_TCELL_DN
FDR q-value = 0.003
NES = 1.60
PRF1 GPR68
TOX PRR5L
ADGRG1 NKG7
TIGIT SYTL3
LAG3 OPTN
GZMA MATK
BHLHE40 IL2RB
SEMA4C ATXN1
TMEM273 CHN2
SRXN1 CST7
GSE26495_NAIVE_VS_PD1LOW_CD8_TCELL_DN
FDR q-value = 0.046
NES = 1.46
PRF1 LL2RB
ADGRG1 TMEM273
TOX DGKQ
PRR5L MATK
LAG3 GZMA
SEMA4C CST7
PDE4A ATXN1
SRXN1 ADAP1
DENND3 EOMES
TIGIT GPR68
GSE26495_NAIVE_VS_PD1HIGH_CD8_TCELL_DN
FDR q-value = 0.006
NES = 1.54
PRF1 GALNT3
TOX GPR68
ADGRG1 MDFIC
TIGIT PRR5L
GZMK NKG7
LAG3 NEDD9
GZMA SYTL3
BHLHE40 OPTN
SEMA4C ZBTB38
SRXN1 GALM
GSE26495_NAIVE_VS_PD1HIGH_CD8_TCELL_DN
FDR q-value = 0.080
NES = 1.42
PRF1 MDFIC
ADGRG1 BCL2L1
TOX GZMA
PRR5L GALM
GZMK CST7
LAG3 ATXN1
SEMA4C SH3BP2
SRXN1 GFL1
GALNT3 EOMES
TIGIT ELP3
GSE9650_NAIVE_VS_EXHAUSTED_CD8_TCELL_DN
FDR q-value = 0.050
NES = 1.39
RGS16 LITAF
SPOCK2 TNFSF10
TNFRSF9 IKZF4
EEA1 LAMC1
IFNG PERP
PRDM1 COCH
GPM68 KCTD9
PLSCR1 CISH
BHLHE40 CTLA4
GEM HCFC1R1
GSE9650_NAIVE_VS_EXHAUSTED_CD8_TCELL_DN
FDR q-value = 0.135
NES = 1.36
TNFRSF9 RPA2
RGS16 LAMC1
SPOCK2 CISH
GPM6B IKZF4
EEA1 PRDM1
CCRL2 VMP1
ARG1 LITAF
HASPIN COCH
TMEM150A PLSCR1
TNFSF10 GEM
GSE9650_EXHAUSTED_VS_MEMORY_CD8_TCELL_UP
FDR q-value = 0.058
NES = 1.38
RGS16 LITAF
TNFRSF4 TWSG1
SPOCK2 ENTPD1
ADGRG1 LAMC1
EEA1 PERP
PRDM1 EGR2
GPM68 CYTH3
PLSCR1 COCH
ALCAM LAT2
ITGAV GPD2
GSE26495_PD1HIGH_VS_PD1LOW_CD8_TCELL_UP
FDR q-value = 0.229
NES = 1.27
PLXND1 GTPBP3
GZMK LRIG1
PDCD1 TNFSF8
TBC1D4 NAT9
ATP6V1G2 CNNM3
SLC43A3 GSR
STON1 CKAP2L
GALM MBD1
CD27 BRCA1
BPHL DGKH
GSE41867_DAY15_EFFECTOR_VS_DAY30_EXHAUSTED
_CD8_TCELL_LCMV_CLONE13_DN
FDR q-value = 0.141
NES = 1.30
TNFRSF9 ARID5B
TMOD1 UBXN8
DNMT3B WDR55
UCHL3 ZBTB25
HIBCH GTF2E1
UBXN2A GNL2
TRMT6 SNRPA1
GALC ZBTB2
ORMDL1 SCRIB
IRAK1BP1 PCMT1
B
Tet2fl/fl vs. Vav1CreTet2KO
Tet2fl/fl vs. Cd4CreTet2KO
58
34
17
Overlapped gene sets associated with T-cell exhaustion
GSE26495_NAIVE_VS_PD1LOW_CD8_TCELL_DN
GSE26495_NAIVE_VS_PD1HIGH_CD8_TCELL_DN
GSE9650_NAIVE_VS_EXHAUSTED_CD8_TCELL_DN
GSE9650_EXHAUSTED_VS_MEMORY_CD8_TCELL_UP
C
Tet2fl/fl
Vav1CreTet2KO
Cd4CreTet2KO
TPM
900
600
300
0
*
**
**
*
**
***
*
***
**
***
Foxp3
Tcf7
Stat3
Tbx21
Id3
Id2
Prdm1
Eomes
Stat1
Tox
Satb1
Tsc22d3
Rel

**Figure S3. Gene set enrichment analysis reveals T-cell exhaustion signatures in *Tet2*$^{fl/fl}$ CD8$^+$ TILs in comparison to *Vav1*$^{Cre}$*/Cd4*$^{Cre}$*Tet2KO* CD8$^+$ TILs.**

(A) GSEA plots showing enrichment of T-cell exhaustion-related gene sets in *Tet2*$^{fl/fl}$ CD8$^+$ TILs compared to *Vav1*$^{Cre}$*Tet2KO* (left) and *Cd4*$^{Cre}$*Tet2KO* CD8$^+$ TILs (right).

(B) Venn diagram showing overlap of T-cell exhaustion-associated gene sets between *Tet2*$^{fl/fl}$ *vs. Vav1*$^{Cre}$*Tet2KO* (pink) and *Tet2*$^{fl/fl}$ *vs. Cd4*$^{Cre}$*Tet2KO* (green) comparisons. 34 gene sets were commonly enriched in *Tet2*$^{fl/fl}$ CD8$^+$ TILs in both comparisons, including four gene sets associated with exhaustion transcriptional program.

(C) Expression levels (TPM) of transcription factors from IPA upstream regulator analysis.

TILs, tumor-infiltrating lymphocytes; GSEA, Gene Set Enrichment Analysis; TPM, transcripts per million; IPA, Ingenuity Pathway Analysis; NES, normalized enrichment score. Significantly enriched gene sets in (A) and (B) were identified by FDR $q$-value < 0.25 and nominal $P$ value < 0.05. Statistical significance was determined by One-way ANOVA with Tukey's HSD test. *$P$ < 0.05, **$P$ < 0.01, ***$P$ < 0.001, ****$P$ < 0.0001, ns = not significant.

### Figure S4

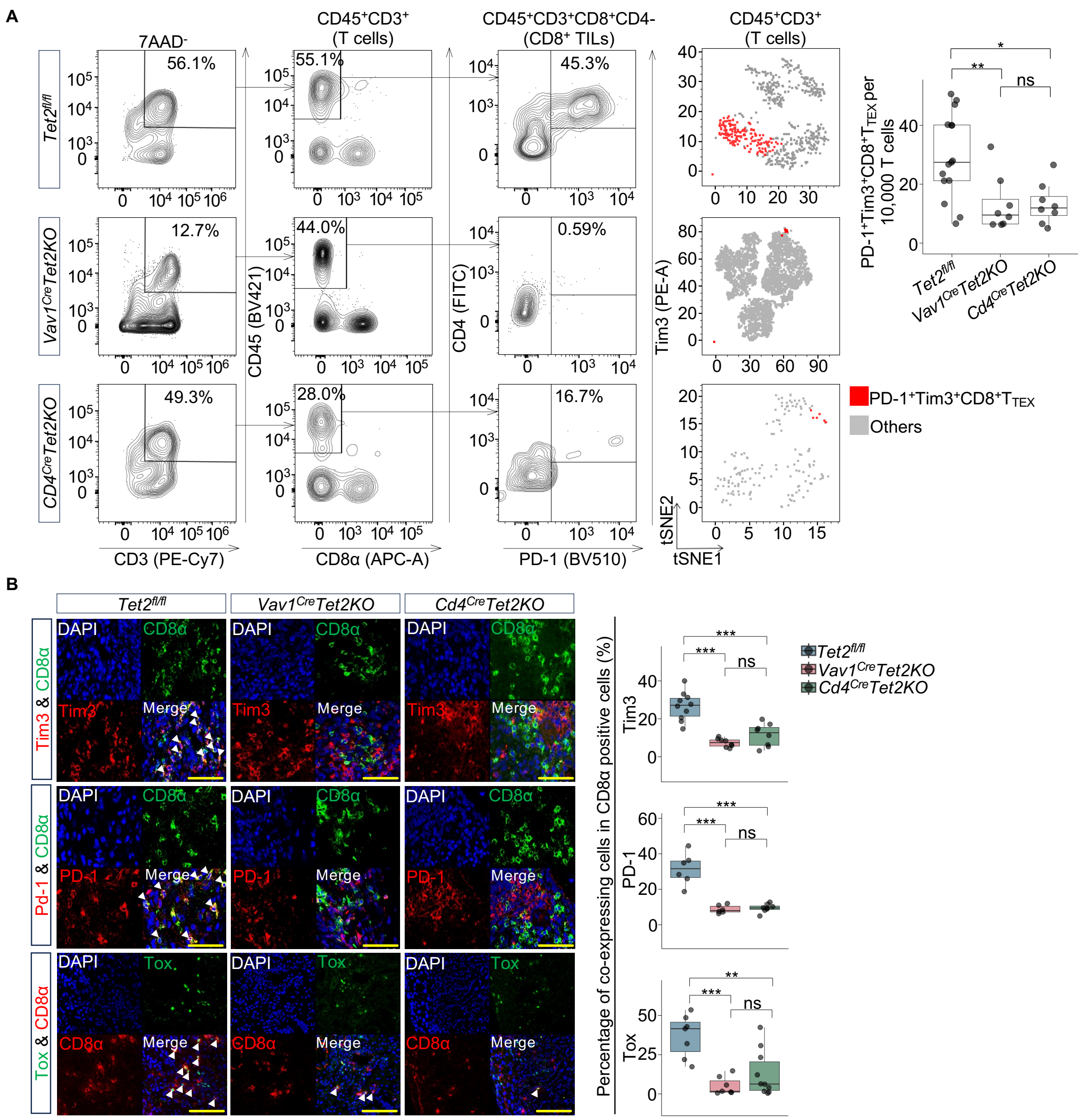


**Figure S4. Flow cytometry and immunofluorescence analysis of exhaustion marker co-expression on CD8+ TILs across *Tet2* genotypes.**

(A) Representative flow cytometry plots showing the proportion of PD-1+Tim3+$T_{TEX}$ in liver metastaic tumors of *Tet2fl/fl* ($n$ = 15), *Vav1CreTet2KO* ($n$ = 8), and *Cd4CreTet2KO* ($n$ = 8) groups. Box plots display the frequencies of PD-1+Tim3+$T_{TEX}$ across the three groups. See also Table S14A for detailed quantification and statistical comparisons.

(B) Representative images of indirect immunofluorescent staining performed on tumor samples demonstrating co-expression of CD8α with Tim3, PD-1, and Tox in *Tet2fl/fl*, *Vav1CreTet2KO*, and *Cd4CreTet2KO* groups. Box plots displaying percentage of cells co-expressing Tim3 (top), and PD-1 (middle), and Tox (bottom) within CD8α+ population. See also Table S14B for detailed quantification and statistical comparisons.

TILs, tumor-infiltrating lymphocytes; $T_{TEX}$, terminally exhausted T cells. Data in (A) and (B) are compiled of at least 2 independent experiments. Statistical significance was determined by ANOVA with Tukey's HSD test. *$P < 0.05$, **$P < 0.01$, ***$P < 0.001$, ****$P < 0.0001$, ns = not significant.

### Figure S5

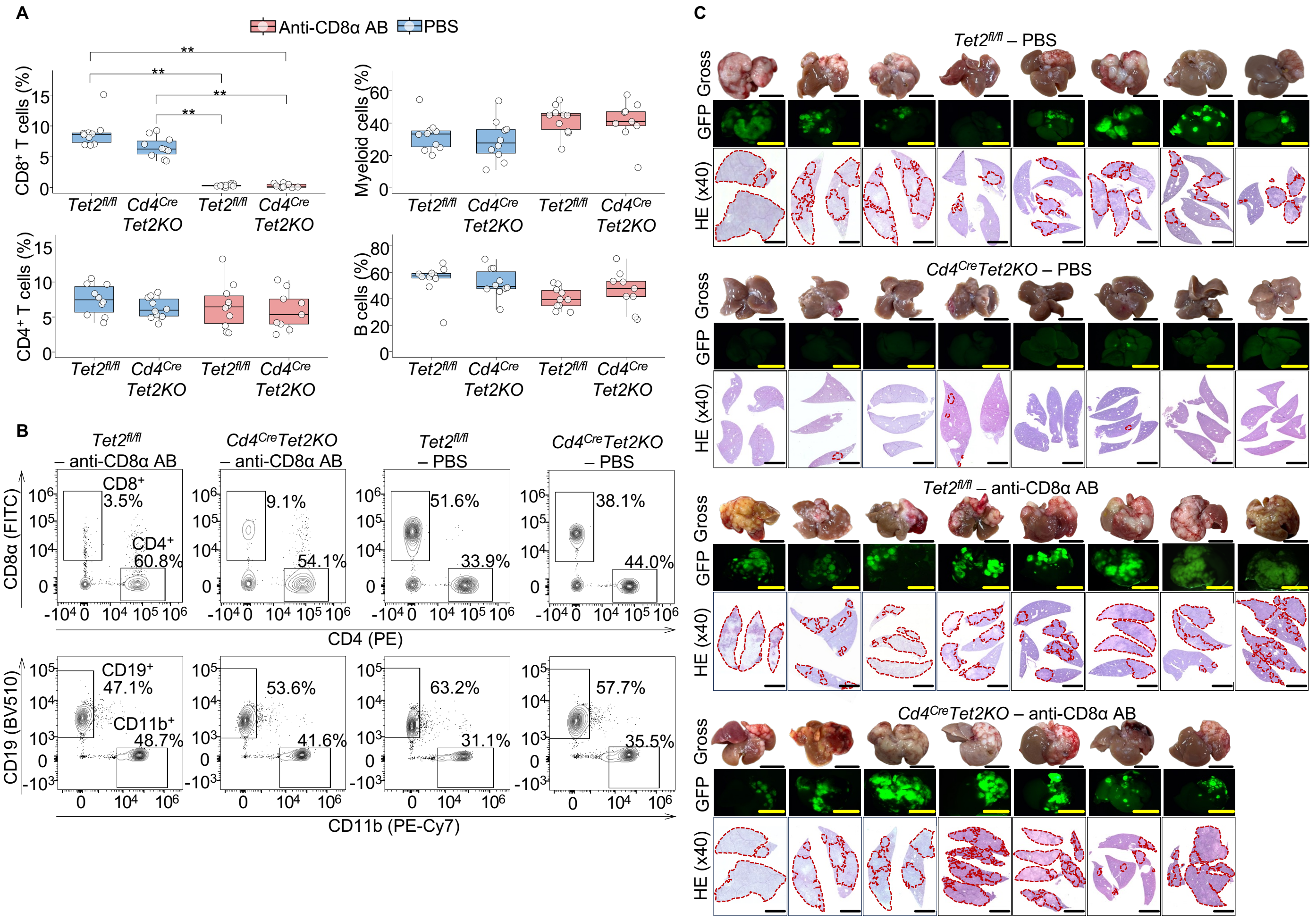


**Figure S5. CD8 T-cell depletion validation and liver metastasis assessment in *Cd4$^{Cre}$Tet2KO* and *Tet2$^{fl/fl}$* mice.**

(A) Box plots show the percentages of CD8 T cells, CD4 T cells, myeloid, and B cells among CD45$^{+}$ viable cells in peripheral blood collected at the end of the anti-CD8α antibody treatment course. Data represent cumulative results from 2 independent experiments with four experimental groups: *Tet2$^{fl/fl}$* - PBS ($n$ = 10), *Cd4$^{Cre}$Tet2KO* - PBS ($n$ = 10), *Tet2$^{fl/fl}$* - anti-CD8α AB ($n$ = 10), and *Cd4$^{Cre}$Tet2KO* - anti-CD8α AB ($n$ = 9).

(B) Representative flow cytometry plots of liver-infiltrating immune cells show the distribution of CD8 T cells, CD4 T cells (upper row, gating on 7-AAD$^{-}$CD45$^{+}$CD3$^{+}$Nk1.1$^{-}$ population), B cells (CD19+), and myeloid cells (CD11b$^{+}$) (lower row, gating on 7-AAD$^{-}$CD45$^{+}$ population), at the end of treatment. Representative plots are shown from one liver sample per group collected at the experimental endpoint.

(C) Liver metastatic burden assessment following CD8 T cell depletion. Each column represents one biological replicate across experimental groups. Top row: Macroscopic appearance of liver samples. Middle row: GFP fluorescence imaging. Bottom row: Representative HE-stained sections (40x); red dashed outlines delineate tumor regions. Scale bars: 1 cm (macroscopic and GFP); 5 mm (HE).

AB, antibody; GFP, green fluorescent protein; PBS, phosphate-buffered saline; Tet2fl/fl - PBS, Tet2fl/fl treated with PBS; Cd4CreTet2KO - PBS, Cd4CreTet2KO treated with PBS; Tet2fl/fl - anti-CD8α AB, Tet2fl/fl treated with anti-CD8α antibody; Cd4CreTet2KO - anti-CD8α AB, Cd4CreTet2KO treated with anti-CD8α antibody. Statistical significance in (A) was determined by pairwise Mann-Whitney U tests with Bonferroni correction for multiple comparisons, only significant comparisons are indicated in the plots. *$P$ < 0.05, **$P$ < 0.01, ***$P$ < 0.001, ****$P$ < 0.0001.

### Figure S6

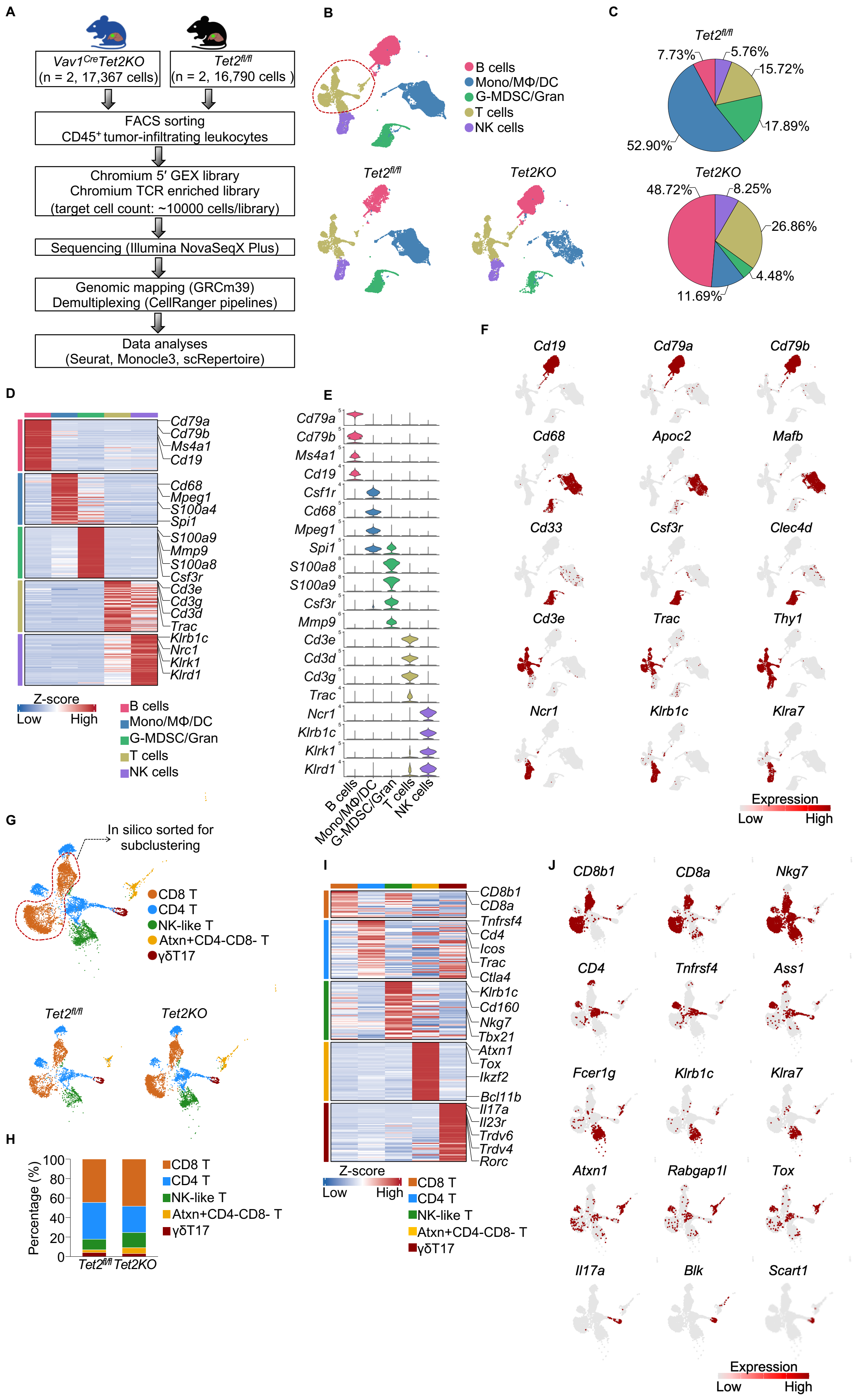
A
Vav1CreTet2KO (n = 2, 17,367 cells)
Tet2fl/fl (n = 2, 16,790 cells)
FACS sorting
CD45+ tumor-infiltrating leukocytes
Chromium 5′ GEX library
Chromium TCR enriched library
(target cell count: ~10000 cells/library)
Sequencing (Illumina NovaSeqX Plus)
Genomic mapping (GRCm39)
Demultiplexing (CellRanger pipelines)
Data analyses
(Seurat, Monocle3, scRepertoire)
B
B cells
Mono/MΦ/DC
G-MDSC/Gran
T cells
NK cells
Tet2fl/fl
Tet2KO
C
Tet2fl/fl
7.73%
5.76%
15.72%
17.89%
52.90%
Tet2KO
48.72%
8.25%
26.86%
4.48%
11.69%
D
Cd79a
Cd79b
Ms4a1
Cd19
Cd68
Mpeg1
S100a4
Spi1
S100a9
Mmp9
S100a8
Csf3r
Cd3e
Cd3g
Cd3d
Trac
Klrb1c
Nrc1
Klrk1
Klrd1
Z-score
Low
High
B cells
Mono/MΦ/DC
G-MDSC/Gran
T cells
NK cells
E
Cd79a
Cd79b
Ms4a1
Cd19
Csf1r
Cd68
Mpeg1
Spi1
S100a8
S100a9
Csf3r
Mmp9
Cd3e
Cd3d
Cd3g
Trac
Ncr1
Klrb1c
Klrk1
Klrd1
B cells
Mono/MΦ/DC
G-MDSC/Gran
T cells
NK cells
F
Cd19
Cd79a
Cd79b
Cd68
Apoc2
Mafb
Cd33
Csf3r
Clec4d
Cd3e
Trac
Thy1
Ncr1
Klrb1c
Klra7
Expression
Low
High
G
In silico sorted for
subclustering
CD8 T
CD4 T
NK-like T
Atxn+CD4-CD8- T
γδT17
Tet2fl/fl
Tet2KO
H
Percentage (%)
100
80
60
40
20
0
Tet2fl/fl
Tet2KO
CD8 T
CD4 T
NK-like T
Atxn+CD4-CD8- T
γδT17
I
CD8b1
CD8a
Tnfrsf4
Cd4
Icos
Trac
Ctla4
Klrb1c
Cd160
Nkg7
Tbx21
Atxn1
Tox
Ikzf2
Bcl11b
Il17a
Il23r
Trdv6
Trdv4
Rorc
Z-score
Low
High
CD8 T
CD4 T
NK-like T
Atxn+CD4-CD8- T
γδT17
J
CD8b1
CD8a
Nkg7
CD4
Tnfrsf4
Ass1
Fcer1g
Klrb1c
Klra7
Atxn1
Rabgap1l
Tox
Il17a
Blk
Scart1
Expression
Low
High

**Figure S6. scRNA-seq characterization of CD45$^+$ tumor-infiltrating leukocytes and T cell subsets in *Vav1$^{Cre}$Tet2KO* and *Tet2$^{fl/fl}$* mice.**

(A) Overview of scRNA-seq for CD45$^+$ tumor-infiltrating leukocytes extracted from *Vav1$^{Cre}$Tet2KO* ($n$ = 2) and *Tet2$^{fl/fl}$* ($n$ = 2) mice.

(B) UMAP plot of integrated data (upper), and UMAP plots of *Vav1$^{Cre}$Tet2KO* and *Tet2$^{fl/fl}$* CD45$^+$ tumor-infiltrating leukocytes shown separately (bottom left and bottom right, respectively). Five lineage clusters (B cells, T cells, Mono/MΦ/DC, G-MDSC/Gran, and NK cells) are colored by cell types. T cells (red dash line) were extracted for further sub-clustering in (G).

(C) Pie graphs displaying percentages of each cluster in *Vav1$^{Cre}$Tet2KO* and *Tet2$^{fl/fl}$* CD45$^+$ tumor-infiltrating leukocytes.

(D) Heatmap showing the expression of top 50 conserved markers in each lineage cluster of CD45$^+$ tumor-infiltrating leukocytes.

(E) Stacked violin plot showing the expression of specific markers extracted from top 50 conserved markers of each lineage cluster.

(F) Feature plots displaying the expressing distribution of specific markers used to defined lineage clusters in (B).

(G) UMAP plot of integrated data of T cells in-silico sorted from (B) (upper), and UMAP plots of *Vav1$^{Cre}$Tet2KO* and *Tet2$^{fl/fl}$* T cells shown separately (bottom left and bottom right, respectively). Five T-cell clusters are colored by cell types. CD8$^+$ TILs (red dash line) were sorted for further analysis (see Figure 4).

(H) Stacked bar plot showing the proportion of each T cell cluster in *Vav1$^{Cre}$Tet2KO* and *Tet2$^{fl/fl}$*.

(I) Heatmap of the top 50 conserved markers in each T cell cluster from (G).

(J) Feature plots displaying the expression of top three conserved markers used to defined T cells clusters in (G).

scRNA-seq, single-cell RNA sequencing; UMAP, uniform manifold approximation and projection; Mono/MΦ/DC, monocytes/ macrophages/ or dendritic cells; G-MDSC/Gran, granulocytic myeloid-derived suppressor cells or granulocytes; NK cells, natural killer cells; γδT17, Interleukin-17–producing gamma delta T cells; TILs, tumor-infiltrating lymphocytes.

### Figure S7

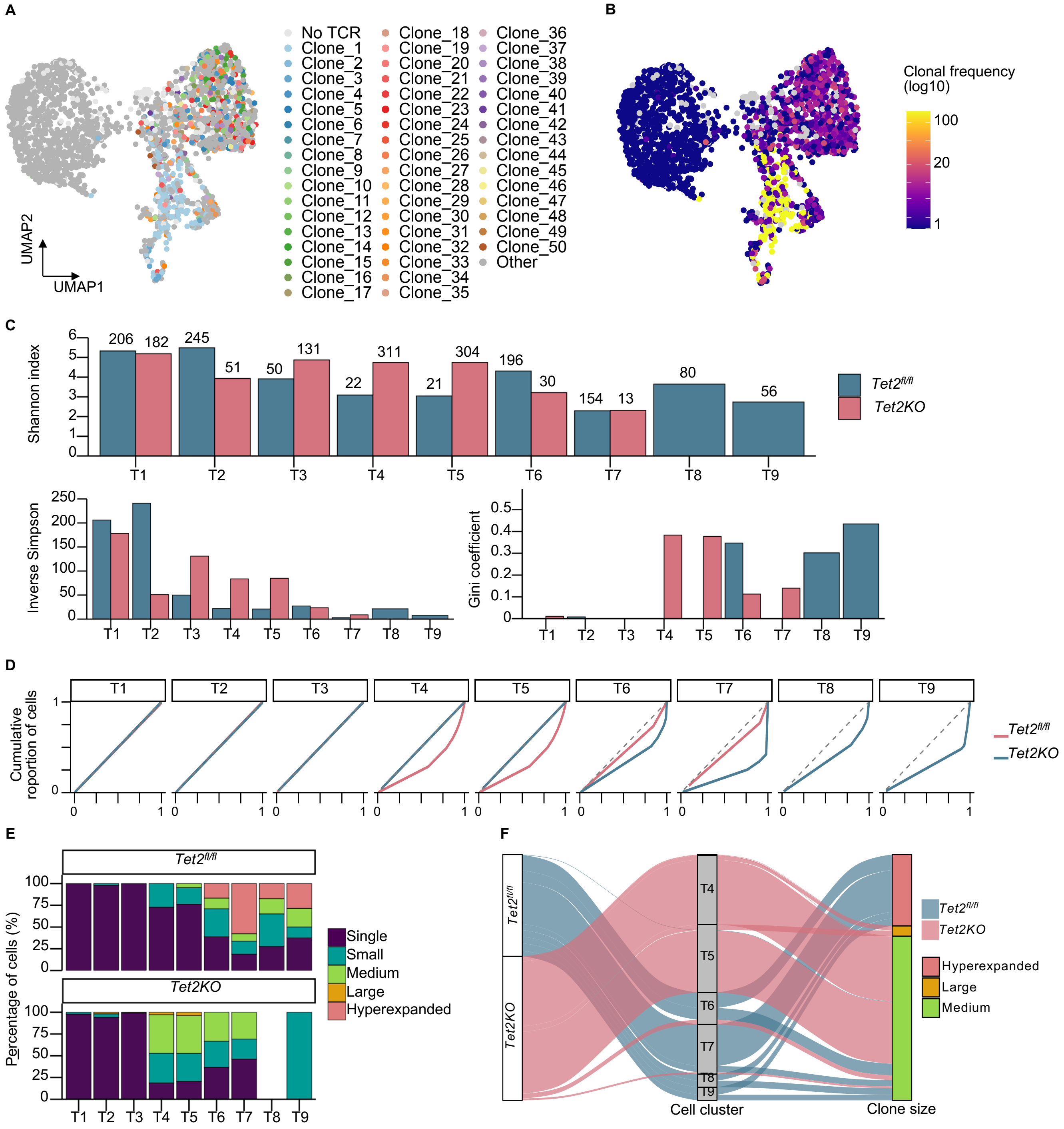
A
UMAP2
UMAP1
No TCR
Clone_1
Clone_2
Clone_3
Clone_4
Clone_5
Clone_6
Clone_7
Clone_8
Clone_9
Clone_10
Clone_11
Clone_12
Clone_13
Clone_14
Clone_15
Clone_16
Clone_17
Clone_18
Clone_19
Clone_20
Clone_21
Clone_22
Clone_23
Clone_24
Clone_25
Clone_26
Clone_27
Clone_28
Clone_29
Clone_30
Clone_31
Clone_32
Clone_33
Clone_34
Clone_35
Clone_36
Clone_37
Clone_38
Clone_39
Clone_40
Clone_41
Clone_42
Clone_43
Clone_44
Clone_45
Clone_46
Clone_47
Clone_48
Clone_49
Clone_50
Other
B
Clonal frequency (log10)
100
20
1
C
Shannon index
206 182 245 51 50 131 22 311 21 304 196 30 154 13 80 56
T1 T2 T3 T4 T5 T6 T7 T8 T9
Tet2fl/fl
Tet2KO
Inverse Simpson
Gini coefficient
D
Cumulative roportion of cells
T1 T2 T3 T4 T5 T6 T7 T8 T9
Tet2fl/fl
Tet2KO
E
Tet2fl/fl
Tet2KO
Percentage of cells (%)
Single
Small
Medium
Large
Hyperexpanded
F
Tet2fl/fl
Tet2KO
Cell cluster
Clone size
Hyperexpanded
Large
Medium

**Figure S7. TCR repertoire analysis of CD8$^+$ TILs shows *Tet2$^{fl/fl}$* mice exhibit hyperexpanded clones in exhausted states while *Vav1$^{Cre}$Tet2KO* maintains clonal diversity.**

(A) UMAP visualization of the top 50 expanded T cell clones (ranked by clone size) across all samples. Each color represents a distinct clonotype based on paired TCRα and TCRβ CDR3 amino acid sequences (CTaa). Grey dots indicate cells without TCR information.

(B) UMAP plot colored by clonal frequency (log10 scale), showing the distribution and expansion status of T-cell clones.

(C) Clonal diversity metrics across T-cell differentiation states (T1-T9) by condition. Top: Shannon diversity index showing the number of unique clones contributing to each cluster (numbers above bars indicate total cells); higher Shannon values indicate greater clonal diversity. Bottom left: Inverse Simpson index representing the effective number of clones contributing to each population; higher values indicate more clones contributing equally. Bottom right: Gini coefficient measuring clonal inequality within each cluster; 0 = perfect equality, 1 = complete dominance by single clone.

(D) Lorenz curves showing clonal inequality across differentiation states. Each panel represents a different T cell state (T1-T9). The x-axis shows cumulative proportion of clones, and y-axis shows cumulative proportion of cells. Curves closer to the diagonal (dashed line) indicate more equal clone size distributions.

(E) Relative abundance of clone size categories between *Tet2$^{fl/fl}$* and *Tet2KO* conditions.

(F) Alluvial (Sankey) diagram showing the distribution of expanded clones (>5 cells) across conditions and T cell differentiation states. Left panel shows condition (*Tet2$^{fl/fl}$ vs. Tet2KO*), middle panel shows cell cluster distribution, and right panel shows clone size categories. Flow width represents the number of cells.

TCR, T-cell receptor; TILs, tumor-infiltrating lymphocytes; UMAP, uniform manifold approximation and projection. Clone size in (H) and (I) is defined as; Hyperexpanded ($100 < X \leq 500$), Large ($20 < X \leq 100$), Medium ($5 < X \leq 20$), Small ($1 < X \leq 5$), Single ($0 < X \leq 1$).

#### Figure S8

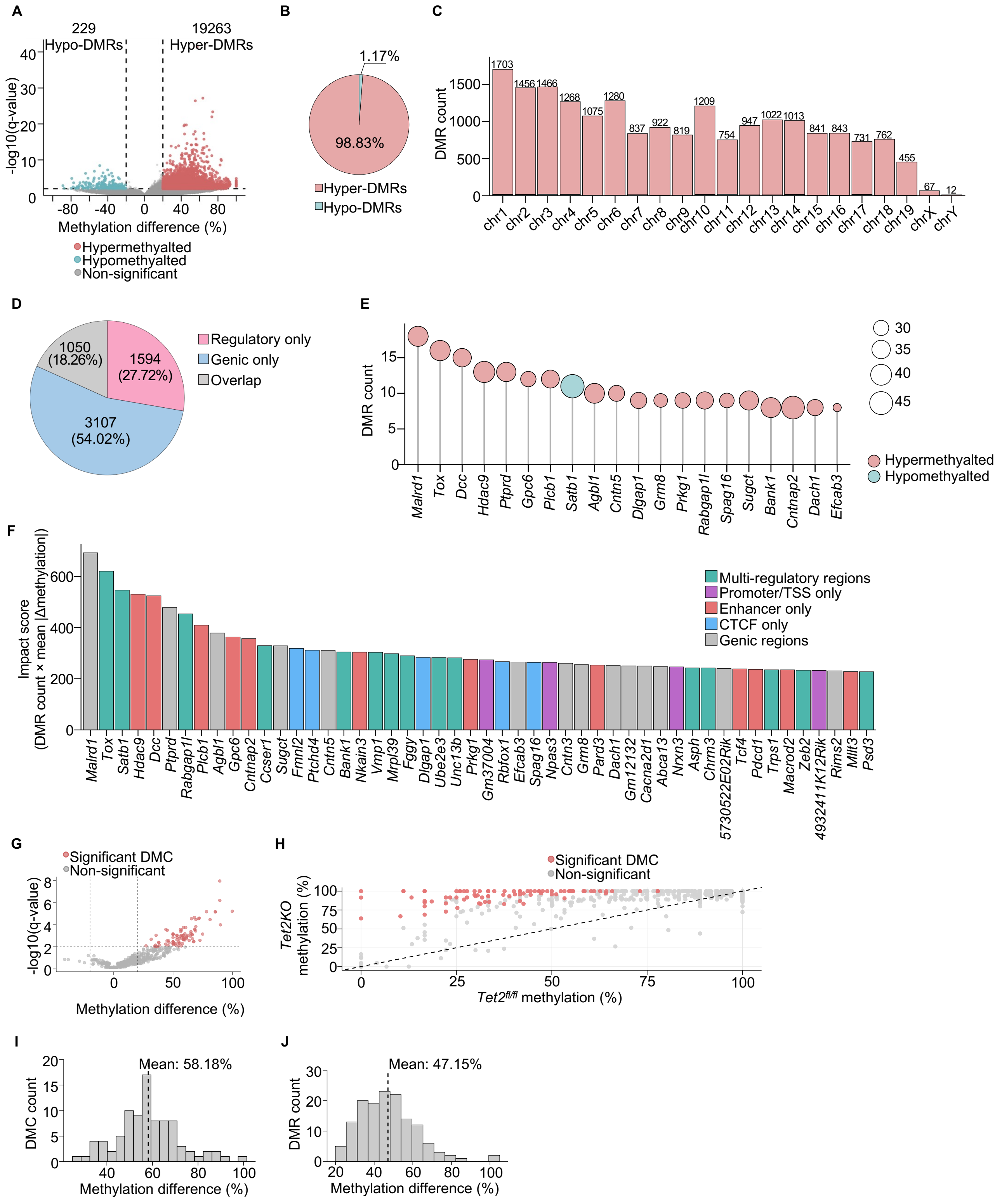
A
229
Hypo-DMRs
19263
Hyper-DMRs
-log10(q-value)
Methylation difference (%)
Hypermethyalted
Hypomethyalted
Non-significant
B
1.17%
98.83%
Hyper-DMRs
Hypo-DMRs
C
DMR count
1703
1456
1466
1268
1075
1280
837
922
819
1209
754
947
1022
1013
841
843
731
762
455
67
12
chr1
chr2
chr3
chr4
chr5
chr6
chr7
chr8
chr9
chr10
chr11
chr12
chr13
chr14
chr15
chr16
chr17
chr18
chr19
chrX
chrY
D
1050
(18.26%)
1594
(27.72%)
3107
(54.02%)
Regulatory only
Genic only
Overlap
E
DMR count
Malrd1
Tox
Dcc
Hdac9
Ptprd
Gpc6
Plcb1
Satb1
Agbl1
Cntn5
Dlgap1
Grm8
Prkg1
Rabgap1l
Spag16
Sugct
Bank1
Cntnap2
Dach1
Efcab3
30
35
40
45
Hypermethyalted
Hypomethyalted
F
Impact score
(DMR count × mean |Δmethylation|)
Multi-regulatory regions
Promoter/TSS only
Enhancer only
CTCF only
Genic regions
Malrd1
Tox
Satb1
Hdac9
Dcc
Ptprd
Rabgap1l
Plcb1
Agbl1
Gpc6
Cntnap2
Ccser1
Sugct
Fmnl2
Ptchd4
Cntn5
Bank1
Nkain3
Vmp1
Mrpl39
Fggy
Dlgap1
Ube2e3
Unc13b
Prkg1
Gm37004
Rbfox1
Efcab3
Spag16
Npas3
Cntn3
Grm8
Pard3
Dach1
Gm12132
Cacna2d1
Abca13
Nrxn3
Asph
Chrm3
5730522E02Rik
Tcf4
Pdcd1
Trps1
Macrod2
Zeb2
4932411K12Rik
Rims2
Mllt3
Psd3
G
Significant DMC
Non-significant
-log10(q-value)
Methylation difference (%)
H
Significant DMC
Non-significant
Tet2KO
methylation (%)
Tet2fl/fl methylation (%)
I
Mean: 58.18%
DMC count
Methylation difference (%)
J
Mean: 47.15%
DMR count
Methylation difference (%)

**Figure S8. Genome-wide DNA methylation profiling by WGBS reveals widespread hypermethylation in *Tet2KO* PD-1⁺CD8⁺ TILs and extensive methylation changes at the *Tox* locus.**

(A) Volcano plot showing genome-wide differential methylation analysis of *Tet2KO* PD-$1^+$CD$8^+$ TILs vs. *Tet2$^{fl/fl}$* PD-$1^+$CD$8^+$ TILs. Each point represents a 1-kb genomic tile. Red points indicate significantly hypermethylated regions (19,263 hyper-DMRs), blue points indicate hypomethylated regions (229 hypo-DMRs), and gray points represent non-significant regions.

(B) Pie chart showing the proportion of hypermethylated (98.83%) versus hypomethylated (1.17%) DMRs among all DMRs in *Tet2KO* PD-$1^+$CD$8^+$ TILs.

(C) Chromosome distribution of significant DMRs. Bar plot shows the number of hyper-DMRs and hypo-DMRs across all mouse chromosomes. Numbers above bars indicate total DMRs per chromosome.

(D) Categorization of DMRs by genomic annotaion. Pie chart showing the distribution of 5,751 unique DMRs with direct overlap to annotated genomic features. DMRs are classified as regulatory (enhancers, promoters, TSS, CTCF binding sites) or genic regions (introns, exons, UTRs, CDS).

(E) Top 20 genes ranked by number of associated DMRs. Genes are ordered by total DMR counts (indicated by circle size). Circle color indicates methylation status.

(F) Top 50 genes ranked by methylation impact score. Bars are colored by DMR location patterns: multi-regulatory, promoters, TSS, and CTCF sites, promoter/TSS only, enhancer only, CTCF only, and genic-only regions.

(G) DMC analysis of *Tox* gene at single-base resolution. Volcano plot shows 735 individual CpG sites with 89 significant DMCs. Red points indicate significant DMCs, gray points are non-significant.

(H) Correlation between *Tet2$^{fl/fl}$* and *Tet2KO* methylation levels across all CpG sites of *Tox*. Red points represent significant DMCs, gray points are non-significant. The deviation of significant DMCs above the diagonal line confirms hypermethylation in *Tet2KO* PD-$1^+$CD$8^+$ TILs.

(I) Histogram shows the distribution of methylation differences across 89 significant DMCs in the *Tox* locus.

(J) Histogram shows the distribution of methylation differences across 56 rDMRs in the *Tox* locus.

WGBS, whole genome bisulfite sequencing; DMR, differentially methylated region; TSS, transcription start site; CTCF, CCCTC-binding factor; CDS, coding sequence; UTR, untranslated region; DMC, differentially methylated cytosine; meth.diff, methylation difference (%). Analysis based on 3 biological replicates per group. Statistical significance to identify DMRs and DMCs determined by *methylKit* using $q$-value < 0.01 and $|\text{meth.diff}| \geq 20\%$. For (F), the multi-regulatory pattern includes genes with DMRs in ≥2 types of regulatory regions, the genic-only pattern includes genes with DMRs located exclusively outside of regulatory regions.

**Figure S9**

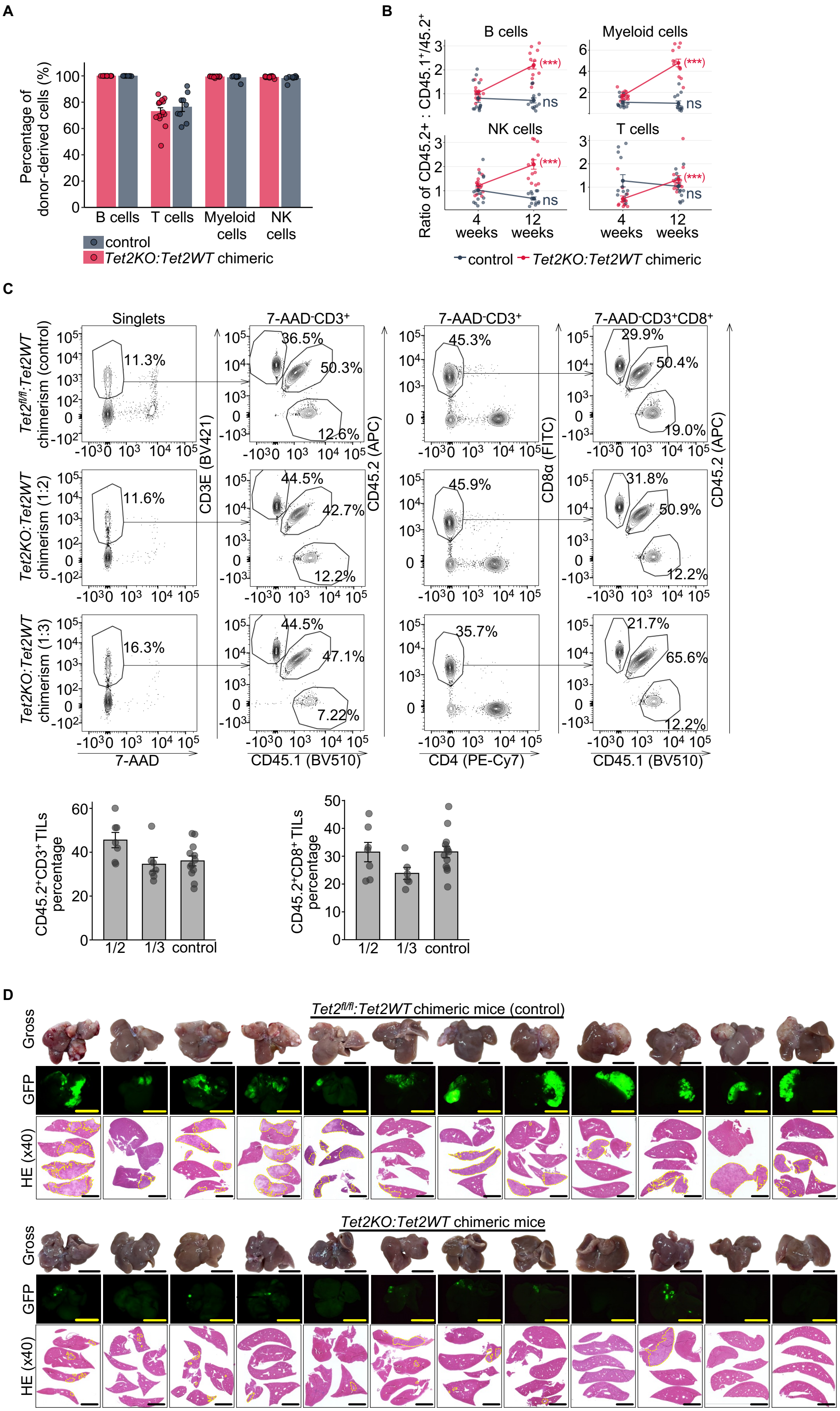

A
Percentage of donor-derived cells (%)
B cells
T cells
Myeloid cells
NK cells
control
Tet2KO:Tet2WT chimeric
B
Ratio of CD45.2+ : CD45.1+/45.2+
B cells
Myeloid cells
NK cells
T cells
(***)
ns
4 weeks
12 weeks
control
Tet2KO:Tet2WT chimeric
C
Singlets
7-AAD-CD3+
7-AAD-CD3+CD8+
Tet2fl/fl:Tet2WT chimerism (control)
Tet2KO:Tet2WT chimerism (1:2)
Tet2KO:Tet2WT chimerism (1:3)
11.3%
36.5%
50.3%
12.6%
45.3%
29.9%
50.4%
19.0%
11.6%
44.5%
42.7%
12.2%
45.9%
31.8%
50.9%
12.2%
16.3%
44.5%
47.1%
7.22%
35.7%
21.7%
65.6%
12.2%
CD3E (BV421)
CD45.2 (APC)
CD8α (FITC)
7-AAD
CD45.1 (BV510)
CD4 (PE-Cy7)
CD45.2+CD3+ TILs percentage
CD45.2+CD8+ TILs percentage
1/2
1/3
control
D
Tet2fl/fl:Tet2WT chimeric mice (control)
Tet2KO:Tet2WT chimeric mice
Gross
GFP
HE (x40)

**Figure S9. Competitive bone marrow transplantation (BMT) mouse model with colorectal cancer liver metastasis.**

(A) Hematopoietic reconstitution at 4 weeks post-BMT. Flow cytometric analysis confirms successful multilineage donor chimerism. B cells, myeloid cells, and NK cells showed near-complete donor-derived reconstitution (>99%). Donor-derived T cells accounted for approximately 73-76% of the total T cell pool, with the remainder representing radioresistant recipient $CD45.1^{+}$ T cells.

(B) Progressive expansion of donor-derived B cells, myeloid cells, NK cells, and T cells from 4 to 12 weeks post-BMT. The expansion was assessed by the longitudinal increase in the ratio of donor $CD45.2^{+}$ to $CD45.1^{+}/CD45.2^{+}$ cells (see also Table S25A and S25B for detailed quantification and statistical comparisons). Each dot represents one mouse; lines connect paired samples.

(C) Liver TIL analysis at 8 weeks post-tumor injection. Top: Representative flow cytometry plots from control, 1:2, and 1:3 chimeras showing gating strategy. Bottom: Quantification of donor-derived $CD45.2^{+}CD3^{+}$ TILs and $CD45.2^{+}CD8^{+}$ TILs (see also Table S25C for detailed quantification).

(D) Whole liver samples collected at the endpoint of the experiment.

TILs, tumor-infiltrating lymphocytes. Data in (A), (B), (C), and (D) are compiled of 3 independent experiments. Statistical significance in (B) was determined by Wilcoxon signed-rank test, and P values were adjusted for multiple comparisons using the Benjamini-Hochberg (FDR) method. *$P < 0.05$, **$P < 0.01$, ***$P < 0.001$, ****$P < 0.0001$, ns = not significant.

#### Figure S10

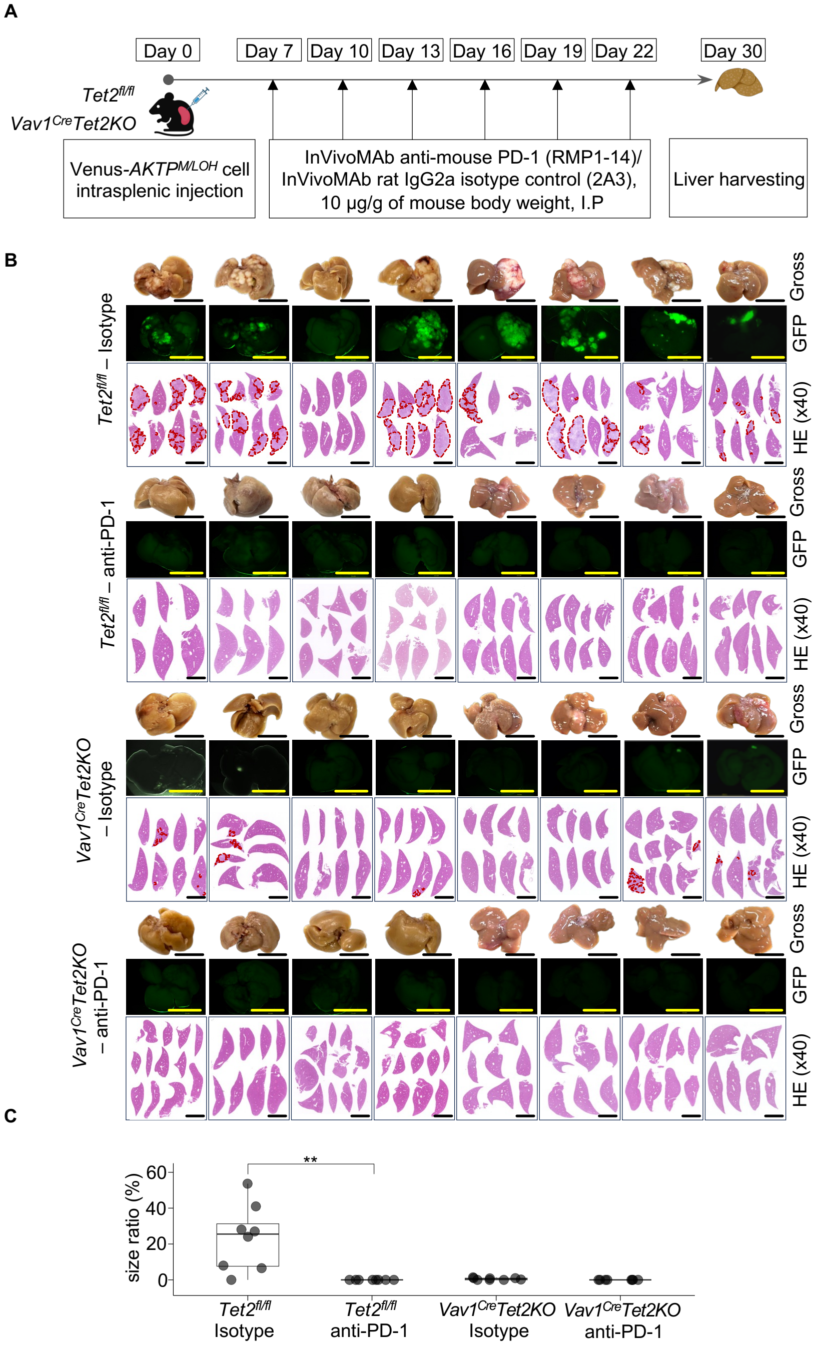


**Figure S10. Anti-PD-1 antibody treatment experiment.**

(A) Experimental schema of anti-PD-1 treatment. *Tet2*$^{fl/fl}$ and *Vav1*$^{Cre}$*Tet2KO* mice received intrasplenic injection of Venus-*AKTP*$^{M/LOH}$ cells on day 0. Starting at day 7, mice were treated intraperitoneally with either InVivoMAb anti-mouse PD-1 (10 µg/g body weight) or rat IgG2a isotype control (10 µg/g) every 3 days for 6 doses.

(B) Whole liver samples in four experimental groups (*n* = 8 per group). Each column represents one biological replicate. For each specimen: macroscopic appearance (top), GFP fluorescence (middle), and HE-stained sections at 40× magnification (bottom, red dashed lines outline tumor regions). Scale bars: 1 cm (macroscopic and GFP), 5 mm (HE 40x).

(C) Box plots displaying LMTB of four experimental groups at the end of anti-PD-1 antibody treatment course.

LMTB, liver metastatic tumor burden; GFP, green fluorescent protein. Data in (B) and (C) are compiled of 2 independent experiments. Statistical significance was determined by Mann-Whitney U test. $*p < 0.05$, $**p < 0.01$, $***p < 0.001$, $****p < 0.0001$, ns = not significant.
